# An AI-Assisted Workflow for Reconstruction, Extension, and Calibration of Quantitative Systems Pharmacology Models

**DOI:** 10.64898/2026.04.05.716273

**Authors:** Igor Goryanin, Oleg Demin, Irina Goryanin

**Affiliations:** School of Informatics, University of Edinburgh, UK; IQANOVA LTD, UK; INSYSBIO UK Ltd, INSYSBIO CY Ltd

**Keywords:** CAR-T therapy, quantitative systems pharmacology, AI-assisted modeling, SBML, T-cell exhaustion, antigen escape, checkpoint inhibition, profile-likelihood identifiability, global sensitivity analysis, model-informed drug development

## Abstract

**Background:** Quantitative systems pharmacology (QSP) models provide mechanistic insight into drug response but are limited by labor-intensive, expert-driven workflows. We developed an AI-assisted QSP (AI-QSP) framework that integrates large language models (LLMs) with SBML-based modeling to enable automated reconstruction, extension, and calibration of mechanistic models.

**Methods:** The framework was applied to a published CAR-T QSP model. The model was reconstructed in SBML and extended via LLM-guided prompts to incorporate key resistance mechanisms: T-cell exhaustion, PD-1/PD-L1 checkpoint regulation, and tumor antigen escape. Model development followed an iterative expert-in-the-loop workflow. The resulting model (21 reactions, 9 species) was calibrated to synthetic benchmark data using 19-parameter optimization. Model credibility was assessed using ASME V&V 40 and ICH M15 principles, including global sensitivity and profile-likelihood analyses.

**Results:** The calibrated model reproduced benchmark dynamics with high accuracy (mean log-RMSE = 0.132). Sensitivity analysis identified CAR-T killing and bystander cytotoxicity as dominant drivers of tumor response. Profile-likelihood analysis showed 71% of parameters were practically identifiable, with remaining parameters prioritised for future data-driven refinement.

**Conclusions:** AI-assisted QSP modeling enables reproducible, scalable model reconstruction and evolution while maintaining mechanistic transparency and regulatory alignment. This framework provides a foundation for accelerating model-informed drug development in cell and gene therapies.

## 1. Introduction

Chimeric antigen receptor T-cell (CAR-T) therapies have transformed the treatment of hematologic malignancies, yet clinical responses remain heterogeneous: while durable remissions are observed in a substantial proportion of patients, treatment failure is frequently linked to limited in vivo CAR-T persistence, T-cell exhaustion, tumor antigen escape, and cytokine-related toxicities [1–3]. Understanding and predicting such nonlinear, patient-specific dynamics require mechanistic frameworks that couple biological realism with quantitative inference [8,9,31].

Quantitative systems pharmacology (QSP) models have emerged as a central methodology for integrating multiscale immune–tumor processes into predictive simulations [10,21,31]. In the context of adoptive cell therapy, QSP approaches enable exploration of the interplay between CAR-T proliferation, exhaustion, cytokine release, and tumor burden under varying therapeutic regimens [10,15,16]. However, traditional QSP development is labor-intensive, relying on manual calibration and limited reproducibility across platforms [21,31].

To address these challenges, we developed an **AI-QSP (Artificial Intelligence–assisted Quantitative Systems Pharmacology)** prototype designed to extend traditional QSP workflows by integrating automated model generation, parameter optimization, and model verification within an AI-assisted modeling environment.

Unlike classical modeling pipelines that rely entirely on manual model construction and calibration, the AI-QSP framework combines **large language models (LLMs)** with established systems pharmacology tools to assist in:

- conversion of literature models into **SBML format**,
- generation of structural model updates based on textual biological knowledge,
- automated parameter calibration, and
- iterative model validation.

All model components are implemented using **Systems Biology Markup Language (SBML) Level 3 Version 2**, enabling full reproducibility and interoperability with standard simulation tools such as Tellurium [26], Heta [27], and DBSolve Optimum [28].

This hybrid framework couples mechanistic transparency with computational intelligence, allowing continuous model refinement as new data become available [8,9].

In this study, the AI-QSP prototype was applied to CAR-T therapy modeling, capturing effector and exhausted CAR-T populations, antigen-positive and -negative tumor subclones, and cytokine feedback (IL-6, IL-10, IFN-**γ**). Our objective was to evaluate the feasibility, strengths, and limitations of AI-assisted automated model evolution. Model behavior under the Triple Combination scenario was then generated and quantitatively fitted to synthetic benchmark data. The framework is evaluated here through a controlled test case: reconstruction of an existing CAR-T QSP model, AI-guided structural extension to include resistance mechanisms, and automated calibration against synthetic benchmark data derived from the original model.

The framework is designed to align with FDA, EMA, and ICH guidance on model-informed development [4,5,33], providing an SBML-encoded, auditable modeling pipeline consistent with regulatory expectations for transparency and reproducibility in computational pharmacology.

The goal of this study is not to develop a new CAR-T model, but to evaluate whether an AI-assisted modeling framework can semi-automated reconstruct, extend, and recalibrate an existing QSP model while preserving its quantitative behavior. Specifically, we assess (i) the capability of LLMs to introduce biologically motivated structural extensions from textual descriptions; (ii) the ability of the iterative expert-in-the-loop workflow to converge on a technically correct SBML implementation; and (iii) the capacity of automated calibration to recover reference-model dynamics under a challenging multi-mechanism scenario.

## 2. Methods

### 2.1 AI-QSP workflow

#### AI-QSP framework overview

The overall architecture of the AI-assisted quantitative systems pharmacology (AI-QSP) framework is illustrated in **Figure 1**. The framework implements an iterative workflow that integrates artificial intelligence–based knowledge interpretation, mechanistic modeling in Systems Biology Markup Language (SBML), and automated parameter optimization. Starting from a literature-derived model description, the system first reconstructs a mechanistic core model in SBML format. The reconstructed model is then verified through simulation to ensure consistency with the reference formulation. Subsequently, the AI system proposes structural model extensions based on textual biological knowledge describing additional mechanisms relevant to the therapeutic system. These proposed modifications are evaluated through expert review, and the feedback is used to refine prompts and regenerate improved model implementations. The workflow iterates between AI-driven model generation and expert validation until a technically consistent model structure is obtained. The final model is calibrated against benchmark datasets using automated parameter estimation and subsequently validated by reproducing the dynamics of the reference model. This iterative process enables systematic reconstruction, extension, and validation of mechanistic pharmacology models while maintaining transparency and reproducibility through standardized SBML representations.

**Figure 1.**
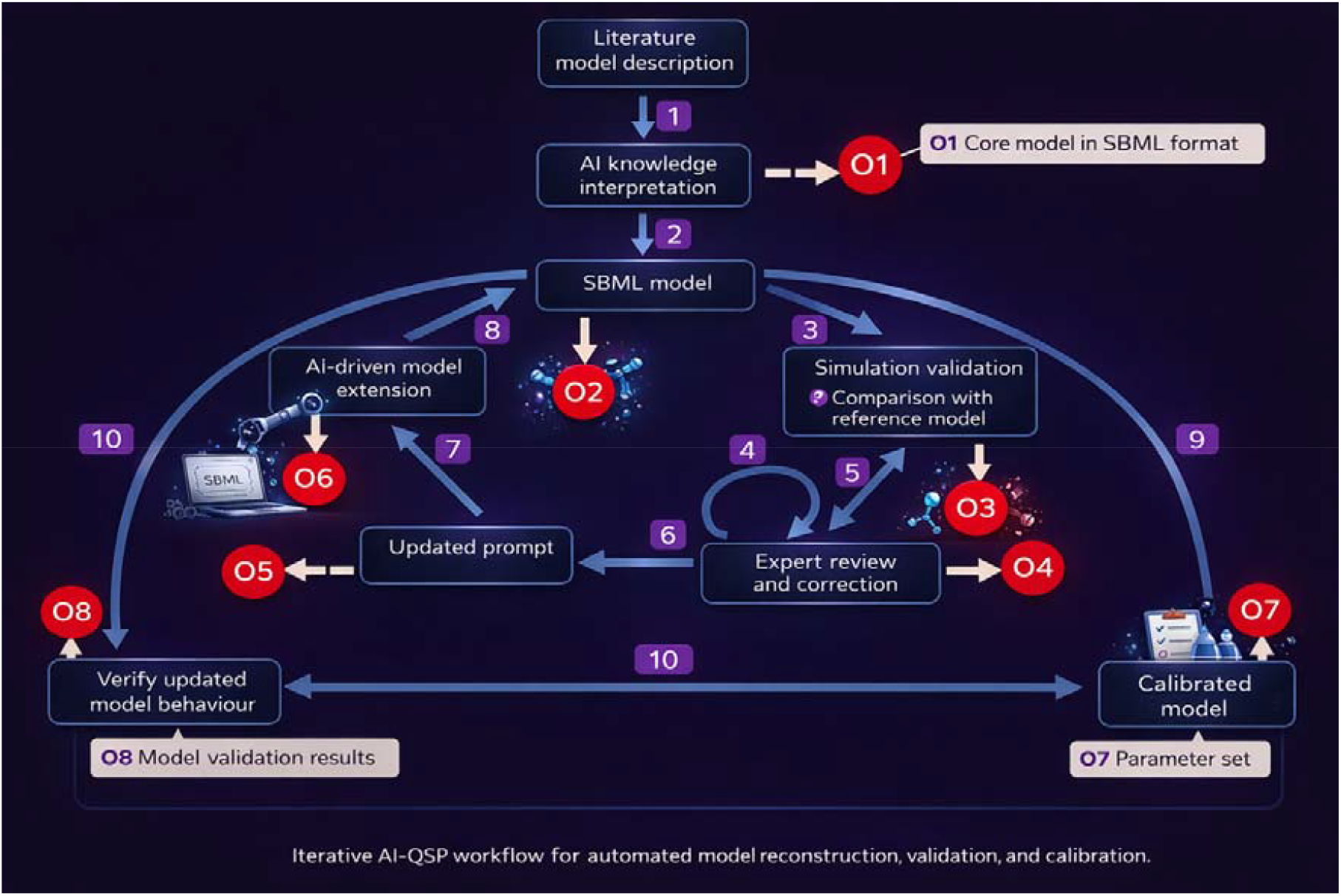
Iterative AI-QSP workflow for automated reconstruction, extension, and calibration of mechanistic pharmacology models. The framework integrates large language model–based knowledge interpretation with SBML-based mechanistic modeling and numerical optimization. A literature model description is first interpreted and converted into an SBML representation of the core model (O1). The reconstructed model is verified through simulation to ensure consistency with the reference implementation (O2–O3). AI-guided model extension introduces additional biological mechanisms based on textual knowledge prompts. Expert review and correction ensure biological plausibility and technical consistency (O4). Updated prompts are generated iteratively to refine model structure (O5–O6). The resulting model is calibrated using automated parameter optimization (O7) and subsequently validated by comparing simulated dynamics with benchmark data (O8). The workflow forms a closed loop enabling progressive improvement of mechanistic QSP models through AI-assisted model evolution.

**Figure 2.**
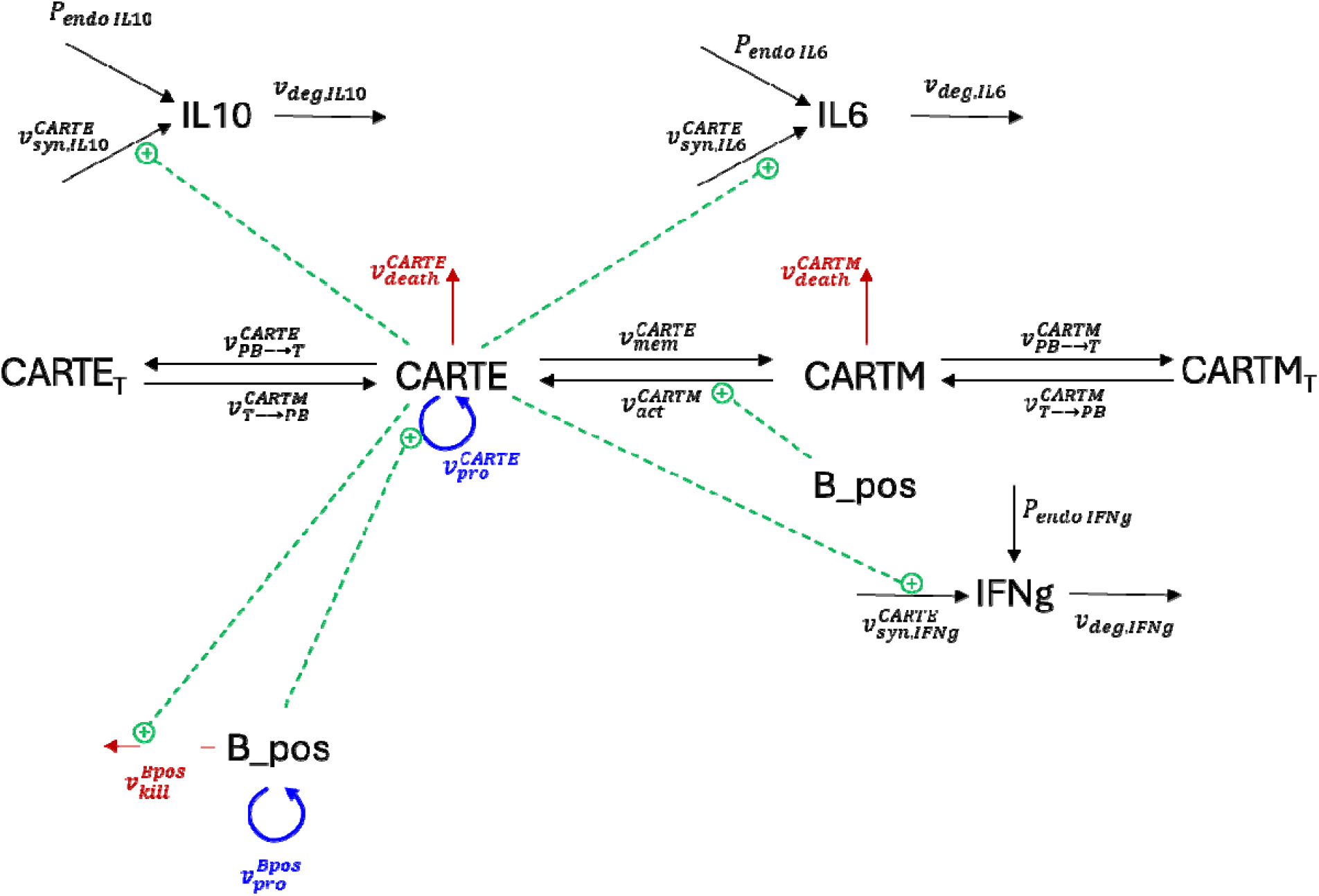
Mechanistic structure of the core CAR-T QSP model. The model describes the dynamics of antigen-positive tumor cells (Bpos), effector CAR-T cells (CARTE), and memory CAR-T cells (CARTM). CARTE proliferation is driven by antigen stimulation from Bpos. CARTE cells can differentiate into memory CAR-T cells (CARTM). Effector CAR-T cells mediate tumor killing of Bpos. Both CARTE and CARTM undergo natural death processes. Tumor cells proliferate logistically.

#### Description of workflow steps

The numbered elements in **Figure 1** correspond to the sequential steps of the AI-QSP workflow. In **Step 1**, a literature-derived model description is interpreted by the AI system, which extracts relevant biological entities, processes, and modeling instructions. In **Step 2**, these elements are translated into a mechanistic model encoded in SBML format, producing the reconstructed core model (Output O1). In **Step 3**, the SBML model is simulated and compared with the behavior of the reference model to verify the correctness of the reconstruction (Outputs O2–O3). In **Step 4**, the AI system proposes structural updates to the model based on textual descriptions of additional biological mechanisms. These updates are examined during **Step 5**, where a domain expert evaluates the generated model structure and identifies technical inconsistencies or biologically implausible mechanisms (Output O4). The expert feedback is then used in **Step 6** to refine the prompts guiding the AI model generation process (Output O5). In **Step 7**, the refined prompts are used to regenerate an updated model implementation incorporating the proposed mechanisms (Output O6). **Step 8** represents the integration of the AI-generated model extensions into the SBML framework, producing an updated mechanistic model. In **Step 9**, the resulting model is calibrated using automated parameter estimation to reproduce benchmark datasets (Output O7). Finally, in **Step 10**, the calibrated model is validated by comparing its simulated dynamics with reference model behavior and benchmark observations (Output O8). Together, these steps form a closed iterative loop that enables progressive refinement and validation of mechanistic QSP models using AI-assisted model development.

The **AI-QSP prototype** integrates large language models (LLMs) with Python-based modeling tools to support automated QSP model development. The framework operates through structured prompt workflows that guide the AI system in interpreting biological knowledge, generating model updates, and producing executable model code.

The AI system interacts with modeling tools executed locally within a federated computational environment to ensure protection of proprietary and confidential data. Human experts validate each step of the workflow to ensure biological plausibility and technical correctness.

The main functional roles of the AI-QSP system include:

1. **Conversion of literature models into SBML format**
2. **Identification of missing biological mechanisms** from textual descriptions
3. **Generation of structural model updates**
4. **Automated code generation and modification**
5. **Assistance in model calibration workflows**

Human expert validation remains an essential component of the process to ensure model correctness.

The AI-QSP framework can be executed locally within a federated environment to ensure protection of proprietary and confidential data.

To validate the AI-QSP prototype, we designed a closed-loop tuning and validation workflow (Figure 1). This framework integrates a mechanistic QSP model, encoded in SBML, with an algorithmic parameter optimization engine and automated data-driven feedback loops.

The workflow’s primary test, as outlined in this study, was its ability to autonomously update core QSP model to include new processes, species, regulations and phenomena then calibrate the complex CAR-T QSP model against a pre-defined, ground-truth benchmark. Each cycle begins with the SBML base model and an imported dataset. An algorithmic parameter optimization strategy explores the parameter space to minimize the error between simulated and observed data. The framework then automatically generates visual and quantitative reports, such as overlay plots (Figure 6) and goodness-of-fit statistics, before the process repeats for subsequent scenarios.

### 2.2 AI implementation

The AI-QSP framework was implemented in Python using a modular architecture combining large language models with mechanistic modeling tools.LLM componentOpenAI GPT-based language models were used to interpret biological text descriptions and generate candidate model updates.Execution environmentAll AI interactions were executed through a Python interface that converts LLM-generated instructions into structured SBML model modifications.Model compilationGenerated model structures were validated using Tellurium.Expert validationAll AI-generated model modifications were inspected by a QSP expert prior to integration into the model.

### 2.3 Mechanistic Core Model

The reference model used for validation of the AI-QSP framework was a previously published quantitative systems pharmacology (QSP) model describing the dynamics of chimeric antigen receptor T-cell (CAR-T) therapy. The model captures key interactions between tumor cells, CAR-T cell populations, and cytokine signaling processes that regulate therapeutic response, immune activation, and treatment resistance mechanisms [10].

To ensure reproducibility and interoperability, the model was implemented using **Systems Biology Markup Language (SBML) Level 3 Version 2**, a widely adopted standard for representing and exchanging mechanistic biological models [25]. SBML encoding allows models to be executed and analyzed across multiple simulation environments commonly used in systems pharmacology and computational biology. In this study, simulations and structural validation of the model were performed using several SBML-compatible modeling platforms, including **Tellurium** [26], **Heta** [27], and **DBSolve Optimum** [28]. These environments support deterministic ordinary differential equation (ODE) simulation, parameter estimation, and model inspection. The SBML representation also enables compatibility with additional modeling frameworks and numerical engines based on **libSBML** and **libRoadRunner**, facilitating reproducible simulation and cross-platform verification of the model implementation [29,30].

The mechanistic structure of the core model includes several interacting biological entities representing key components of the CAR-T therapeutic system.

#### CAR-T cell populations

- Effector CAR-T cells (**CARTE**)
- Memory CAR-T cells (**CARTM**)

The core model includes effector CAR-T cells (CARTE) and memory CAR-T cells (CARTM), representing activated cytotoxic and long-lived memory populations respectively. The exhausted CAR-T state (CARTE_EX), which arises during prolonged antigen stimulation, is introduced as a new species by the AI-assisted model extension described in Section 2.4.

#### Tumor cell populations

- Antigen-positive tumor cells (**B_pos**)

The core model tracks antigen-positive tumor cells (B_pos) only. Antigen-negative tumor cells (**B_neg**), representing tumor cells that have lost the targeted surface antigen through **antigen escape** [2,3], are introduced by the AI-assisted model extension (Section 2.4).

#### Cytokine signaling molecules

- Interleukin-6 (**IL-6**)
- Interleukin-10 (**IL-10**)
- Interferon-γ (**IFN-γ**)

These cytokines represent key inflammatory mediators associated with CAR-T expansion, immune activation, and systemic cytokine responses during therapy.

#### Therapeutic agent

- Anti-PD-1 checkpoint inhibitor (**aPD1**)

Anti-PD-1 checkpoint inhibitor (aPD1) is not present in the core model. It is introduced as an explicit therapeutic species by the AI-assisted model extension (Section 2.4), where it modulates CAR-T functional exhaustion by attenuating PD-1 inhibitory signaling, thereby enhancing CAR-T persistence and cytotoxic activity [1].

The model is formulated as a system of coupled **ordinary differential equations (ODEs)** describing the temporal evolution of cell populations and cytokine concentrations. A **single well-mixed compartment** is assumed, approximating the combined tumor and peripheral blood environment. Cellular proliferation, differentiation, exhaustion, tumor growth, antigen escape, and CAR-T-mediated cytotoxicity are described using mechanistically interpretable kinetic rate laws consistent with established QSP modeling practices [31].

Three split-dose intravenous CAR-T administrations were implemented to reproduce the dosing schedule described in the reference study. The total CAR-T infusion dose was divided into three fractions administered on consecutive days: 10% of the total dose on day 0, 30% on day 1, and 60% on day 2. This staggered infusion strategy reflects commonly used clinical protocols designed to mitigate acute cytokine release syndrome while maintaining therapeutic cell expansion. In the model, each dose fraction is introduced as an external input to the effector CAR-T cell population (CARTE). All model variables, parameters, and rate constants were defined with explicit physical units to ensure dimensional consistency and reproducibility of simulations.

### 2.4 AI-QSP–Based Automatic Update of the AIcore QSP Model

To extend the reconstructed AIcore QSP model, the AI-QSP framework was used to identify additional biological mechanisms relevant to CAR-T therapy that were not explicitly represented in the initial model. This step was designed to evaluate the capability of the framework to interpret biological knowledge from textual descriptions and translate this knowledge into mechanistic model extensions.

Large language models (LLMs) are particularly suitable for this task because they can extract mechanistic relationships and biological entities from unstructured scientific text and convert them into structured representations suitable for computational modeling. In the AI-QSP framework, LLMs were therefore used as a knowledge-interpretation layer that translates biological descriptions into candidate model components, including new species, reactions, and regulatory interactions.

The prompts provided to the AI system focused on identifying, interpreting, and implementing mechanisms associated with well-known resistance processes observed in CAR-T therapy. In particular, the framework was instructed to incorporate biological knowledge related to the following phenomena:

- **T-cell exhaustion**, representing the functional decline of activated CAR-T cells during prolonged antigen stimulation.
- **PD-1/PD-L1 checkpoint signaling**, which suppresses T-cell activity through inhibitory receptor–ligand interactions at the immunological synapse.
- **Tumor antigen escape**, whereby tumor cells lose expression of the targeted antigen and thereby evade CAR-T-mediated cytotoxicity.

These mechanisms are known to reduce CAR-T proliferation and cytotoxic efficacy and are therefore important determinants of treatment response and resistance.

Based on the textual prompts, the AI-QSP system generated proposed updates to the SBML model structure. These updates included the introduction of new state variables, additional reactions describing biological transitions, and modified rate laws capturing regulatory interactions. The proposed model extensions were subsequently integrated into the SBML representation and evaluated through simulation and expert review as part of the iterative workflow described in **Figure 1**.

### 2.5 Example prompt used in the AI-QSP framework

The following simplified prompt illustrates the instructions used to guide the AI-assisted model update process:

`Given an SBML model describing CAR-T therapy dynamics with the species CARTE, CARTM, B_pos, cytokines, and tumor growth processes:`

1. `Identify important biological mechanisms in CAR-T therapy that are not currently represented in the model.`
2. `Focus on the following mechanisms:`
  - `T-cell exhaustion`
  - `PD-1 / PD-L1 checkpoint regulation`
  - `tumor antigen escape`
3. `Propose model extensions by:`
  - `introducing new state variables if required`
  - `defining new reactions or transitions`
  - `modifying rate laws if necessary`
4. `Describe the proposed model updates in SBML-compatible form including species, parameters, and kinetic expressions.`

The generated model modifications were subsequently evaluated by several LLMs and domain experts to ensure biological plausibility and technical correctness before being incorporated into the updated model implementation.

### 2.6 Simulation Scenario

To evaluate the ability of the AI-QSP framework to reproduce the behavior of the reference model, simulations were performed using a **single integrated mechanistic scenario** that incorporates the principal biological mechanisms influencing CAR-T therapy dynamics. This scenario includes CAR-T cell exhaustion, checkpoint inhibition through anti-PD-1 therapy, tumor antigen escape, and bystander-mediated cytotoxicity.

In the model implementation, these mechanisms were represented by assigning non-zero values to the rate constants governing each mechanism (set to zero in baseline single-mechanism scenarios), thereby switching each pathway on within the SBML reaction network. Specifically, the exhaustion transition from effector CAR-T cells to the exhausted phenotype was enabled by setting the exhaustion rate parameter to a positive value *k_exh > 0*. Tumor antigen escape was introduced through conversion of antigen-positive tumor cells (**B_pos**) to antigen-negative cells (**B_neg**) via the antigen-loss rate *k_loss* > 0. Checkpoint blockade was represented by the presence of circulating anti-PD-1 (**aPD1**), which reduces the effective exhaustion rate through the regulatory relationship defined in Eq. (3). In addition, a bystander-mediated killing mechanism was included through activation of the bystander killing rate *k_byst > 0*, enabling partial elimination of antigen-negative tumor cells.

This **combined mechanistic configuration (“Triple Combination” scenario)** (note: the term refers to the three principal resistance axes — exhaustion, checkpoint regulation, and antigen escape — as framed in the reference model [10]; bystander killing functions as an auxiliary cytotoxic mechanism) was selected because it simultaneously captures the major biological processes known to influence CAR-T therapeutic outcomes, including immune exhaustion, immune checkpoint regulation, tumor escape, and secondary cytotoxic mechanisms. Using this integrated scenario provides a stringent test of the AI-QSP framework’s ability to reconstruct and calibrate a mechanistic model capable of reproducing complex CAR-T treatment dynamics.

### 2.7 Ground-Truth Benchmark Generation

To evaluate the calibration performance of the AI-QSP framework, a synthetic dataset was generated using simulations of the reconstructed AIcore QSP model. This dataset served as a **ground-truth benchmark** for testing whether the updated AI-generated model could reproduce the dynamics of the reference model.

The model was simulated over a **28-day post-infusion window (nine observation time points, corresponding to the measurement schedule of the reference study)**, consistent with the time scale reported in the original study [2], where long-term kinetics of CAR-T cells and cytokine responses were analyzed. Synthetic observations were generated for key model variables including **total tumor burden**

(*Total_Tumor = B_pos + B_neg*), **circulating effector CAR-T cells (CARTE_PB)**, and cytokine concentrations (**IFN-γ, IL-6, IL-10**).

Observation time points were selected to correspond to the measurement schedule reported in the reference study [2], capturing both the early expansion phase of CAR-T cells and the later contraction and persistence phases. These time points span the entire simulation interval and allow evaluation of the model’s ability to reproduce both **short-term treatment dynamics and long-term system behavior**.

To emulate biological variability and measurement uncertainty typically observed in clinical datasets, **multiplicative log-normal noise** was applied to the simulated values. The noise distribution was defined using a **coefficient of variation (CV) of 20%**, representing moderate experimental variability commonly reported in immunological measurements.

The resulting dataset provides a **self-consistent synthetic benchmark** derived directly from the reference model. This benchmark enables quantitative assessment of the AI-QSP calibration workflow by testing whether the updated model implementation can reproduce the **28-day post-infusion dynamics of tumor burden, CAR-T expansion and contraction, and cytokine responses** with comparable accuracy to the original model.

### 2.8 Automated Parameter Optimization and Evaluation

Model calibration was performed by the AI-QSP framework’s optimization engine. The objective was to minimize the log-transformed root mean square error (log-RMSE) between the model simulations and the synthetic benchmark data.

A random-restart global search strategy, implemented in Python with Tellurium and NumPy, was used to sample 200 candidate parameter vectors within plausible biological bounds. The full calibration comprised 19 parameters: seven kinetic parameters governing CAR-T and tumor dynamics (listed below), plus six cytokine production and stimulation parameters (**α** and **β** for IL-6, IL-10, and IFN-**γ**), three cytokine degradation rate constants (k_deg, IL6, k_deg, IL10, k_deg, IFN**γ**), two antigen-recognition parameters (K_DX and K_kill), and one baseline degradation constant (k_deg,base). The seven primary kinetic parameters were:

- CARTE expansion rate constant k_prolif
- Exhaustion rate k_exh
- Cytotoxic killing efficiency KBC
- Antigen-loss rate k_loss
- Bystander-killing rate k_byst
- Effector death scaling factor dE_scale
- Anti-PD-1 efficacy Emax_PD1

Model fidelity was assessed by the final log-RMSE score across six calibration variables and by visual inspection of overlay plots (Figure 6). The Mechanistic Reaction-Based model (21 SBML reactions, 19 calibrated parameters) achieved a mean log-RMSE of 0.132, with exhausted CAR-T cells (log-RMSE = 0.085) and IL-6 (log-RMSE = 0.067) meeting the < 0.10 accuracy threshold. All SBML models, fitting scripts, parameter files, and benchmark data are available to ensure full reproducibility.

## 3. Results

### 3.1 Analysis of structure of updated QSP model generated by AI-QSP prototype

We have compared structure of initial AIcore QSP model with AI-QSP generated “AIupdate2” version of the model. This analysis includes comparison of variables, processes and expressions for rate laws of the models.

#### 3.1.1 Variables

The AIupdate2 model differs from the AIcore model in both the set of state variables and the implemented biological processes. Indeed, there are 3 groups of variables in these model versions:

- Group 1: Included in AIcore version only
- Group 2: Included in “AIupdate2” version only
- Group 3: Included in both versions

Group 1 includes variables describing CART cells in tissue. Group 2 includes variables describing exhausted state of CART cells in blood and antigen negative tumor cells. Group 3 includes variables describing effector and memory CART cells in blood, antigen positive tumor cells and cytokine concentrations.

#### 3.1.2 Processes

“AIupdate2” and AIcore versions of the model differ each other in processes. Indeed, “AIupdate2” version does not include processes describing migration of CART cells in tissue but AIcore does not include transition between effector and exhausted states of CART cells and transition between antigen positive and negative tumor cells.

#### 3.1.3 Expression for rate laws

Difference in expressions of rate laws between “AIupdate2” and AIcore versions of the model are summarized in Table 1. Main differences are summarized in Table 1. The AIcore model includes tissue-compartment migration reactions for both effector and memory CAR-T cells that are absent from AIupdate2; conversely, AIupdate2 introduces exhausted CAR-T state transitions (Rxn_CARTE_Exhaustion), antigen-negative tumor cell dynamics (Rxn_Bneg_growth, Rxn_Bneg_Bystander_Killing, Rxn_Antigen_Escape), and a bystander killing mechanism, none of which are present in AIcore. Empty cells in the AIcore column indicate processes unique to AIupdate2 [10].

**Table 1.** Comparison of rate laws.

| ID | “Alupdate2” QSP model | Expert choice | Comment |
| --- | --- | --- | --- |
| Dose CARTE injected |  |  |  |
| Rxn_CARTE_prolif | $k_{\text{prolif\_eff}} * \text{CARTE} * B_{\text{pos}} / (K_{\text{prolif}} + B_{\text{pos}}) * (1 - \text{CARTE} / \text{CARTE\_max})$ | $V * k_{\text{prolif\_eff}} * \text{CARTE} * B_{\text{pos}} / (K_{\text{prolif}} + B_{\text{pos}}) * (1 - \text{CARTE} / \text{CARTE\_max})$ | Saturation with antigen and resources limitation were taken into account |
| Rxn_CARTM_Activation | $k_{\text{act}} * f_{\text{Ag}} * \text{CARTM}$ | $V * k_{\text{act}} * f_{\text{Ag}} * \text{CARTM}$ | |
| Rxn_CARTE_Exhaustion | $k_{\text{exh\_eff}} * \text{CARTE}$ | $V * k_{\text{exh\_eff}} * \text{CARTE}$ | Exhaustion is implemented |
| Rxn_CARTE_death | $k_{\text{death\_E}} * \text{CARTE}$ | $V * k_{\text{death\_E}} * \text{CARTE}$ | |
| Rxn_CARTE_MemDiff | $k_{\text{mem}} * \text{CARTE}$ | $V * k_{\text{mem}} * \text{CARTE}$ | In context of the model differentiation $E \rightarrow M$ should not negatively depend on antigen |
| Rxn_CARTM_death | $k_{\text{death\_M}} * \text{CARTM}$ | $V * k_{\text{death\_M}} * \text{CARTM}$ | |
| Proliferation of CARTM |  |  | CARTM cells are able to proliferate slowly |
| Rxn_CARTX_death | $k_{\text{death\_X}} * \text{CARTX}$ | $V * k_{\text{death\_X}} * \text{CARTX}$ | |
| Migration of CARTE to tissue from blood |  |  | Tissue compartment and CART cell migration is important |
| Migration of CARTE to blood from tissue |  |  | Tissue compartment and CART cell migration is important |
| Migration of CARTM to tissue from blood |  |  | Tissue compartment and CART cell migration is important |
| Migration of CARTM to blood from tissue |  |  | Tissue compartment and CART cell migration is important |
| Rxn_Bpos_growth | $r\_tumor * B\_pos * (1 - B\_pos / K\_tumor)$ | $V * r\_tumor * B\_pos * (1 - B\_pos / K\_tumor)$ | resources limitation were taken into account |
| Rxn_Bpos_killing | $k\_kill * CARTE * B\_pos / (K\_kill + B\_pos)$ | $V * k\_kill * CARTE * B\_pos / (K\_kill + B\_pos)$ | Killing is saturable function of antigen |
| Death of tumor cells | | $V * k\_kill * CARTE * B\_pos / (K\_kill + B\_pos)$ | It is better to separate of death of tumor cells and killing |
| Rxn_Bneg_growth | $r\_tumor * B\_neg * (1 - B\_neg / K\_tumor)$ | $V * r\_tumor * B\_neg * (1 - B\_neg / K\_tumor)$ | Antigen negative tumor cells are implemented |
| Rxn_Bneg_Bystander_Killing | $k\_bystander * bystander\_factor * CARTE * B\_neg / (K\_kill + B\_neg)$ | $V * k\_bystander * bystander\_factor * CARTE * B\_neg / (K\_kill + B\_neg)$ | Bystander effect is implemented |
| Rxn_Antigen_Escape | $k\_esc * B\_pos$ | $V * k\_esc * B\_pos$ | Antigen escape is implemented |
| Rxn_IL6_Syn_CARTE | $p\_IL6\_E * CARTE$ | $V * p\_IL6\_E * CARTE$ | |
| Rxn_IL6_Syn_Bpos | $p\_IL6\_T * B\_pos$ | $V * p\_IL6\_T * B\_pos$ | |
| Rxn_IL6_Deg | $kdeg\_IL6 * IL6$ | $V * kdeg\_IL6 * IL6$ | |
| IL6 production by endogenous T cells |  |  |  |
| Rxn_IL10_Syn_CARTE | $p\_IL10\_E * CARTE$ | $V * p\_IL10\_E * CARTE$ | |
| Rxn_IL10_Syn_Bpos | $p\_IL10\_T * B\_pos$ | $V * p\_IL10\_T * B\_pos$ | |
| Rxn_IL10_Deg | $kdeg\_IL10 * IL10$ | $V * kdeg\_IL10 * IL10$ | |
| IL10 production by endogenous T cells |  |  |  |
| Rxn_IFNg_Syn | $p\_IFNg\_E * CARTE$ | $V * p\_IFNg\_E * CARTE$ | |
| Rxn_IFNg_Deg | $kdeg\_IFNg * IFNg$ | $V * kdeg\_IFNg * IFNg$ | |
| IFNg production by endogenous T cells |  |  |  |

### 3.2 Application of the AI-QSP workflow to automatic update of the AIcore CAR-T model

Using the reconstructed **AIcore** QSP model as input and applying the prompt workflow described in Section 2.4, the AI-QSP framework generated a first updated version of the CAR-T model (**AIupdate1**), incorporating the requested biological mechanisms related to T-cell exhaustion, PD-1/PD-L1 checkpoint regulation, and tumor antigen escape. The corresponding model scheme and SBML implementation are shown in **Figure 3A** and **Figure 4**, respectively.

**Figure 3.**
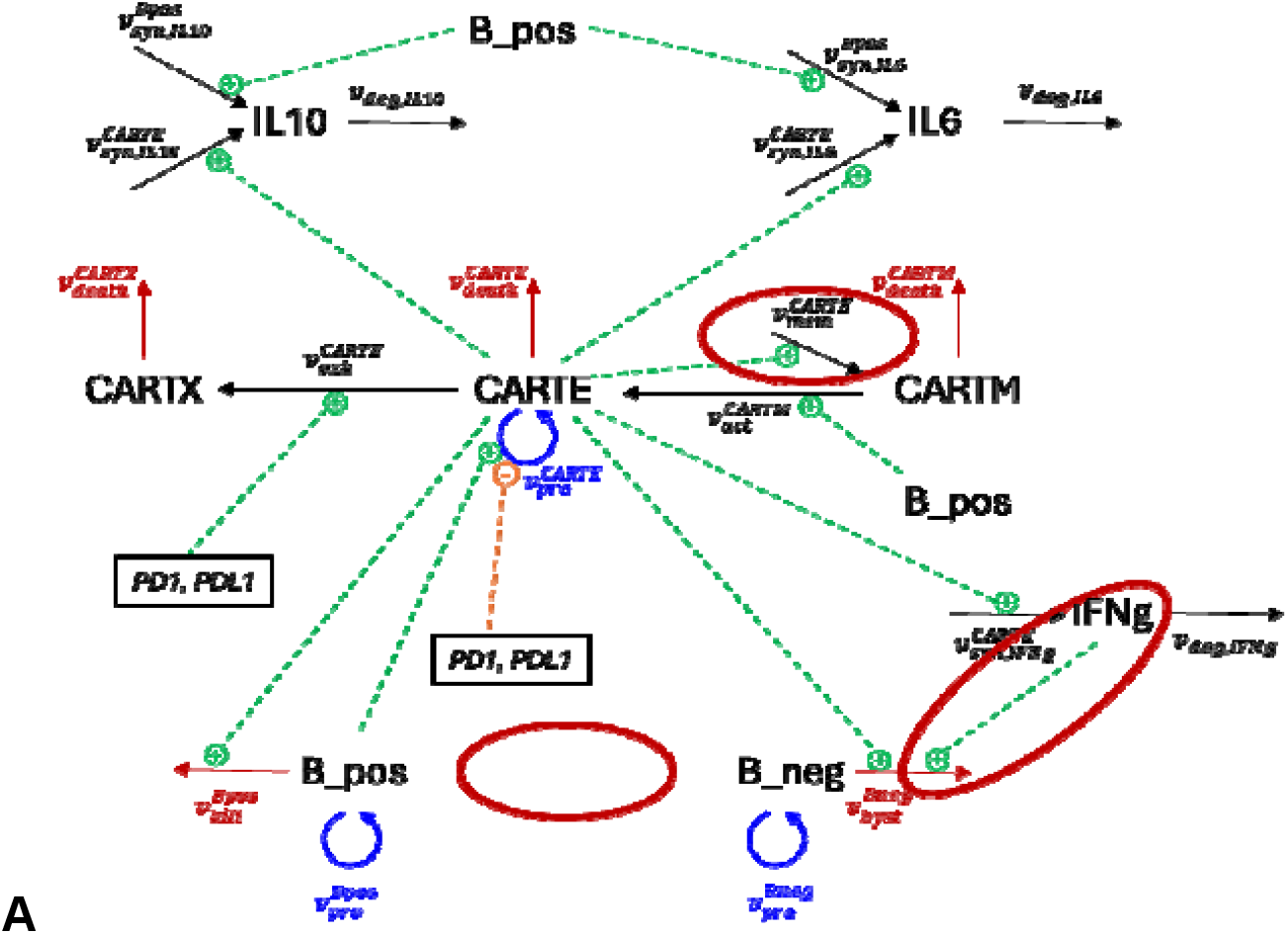
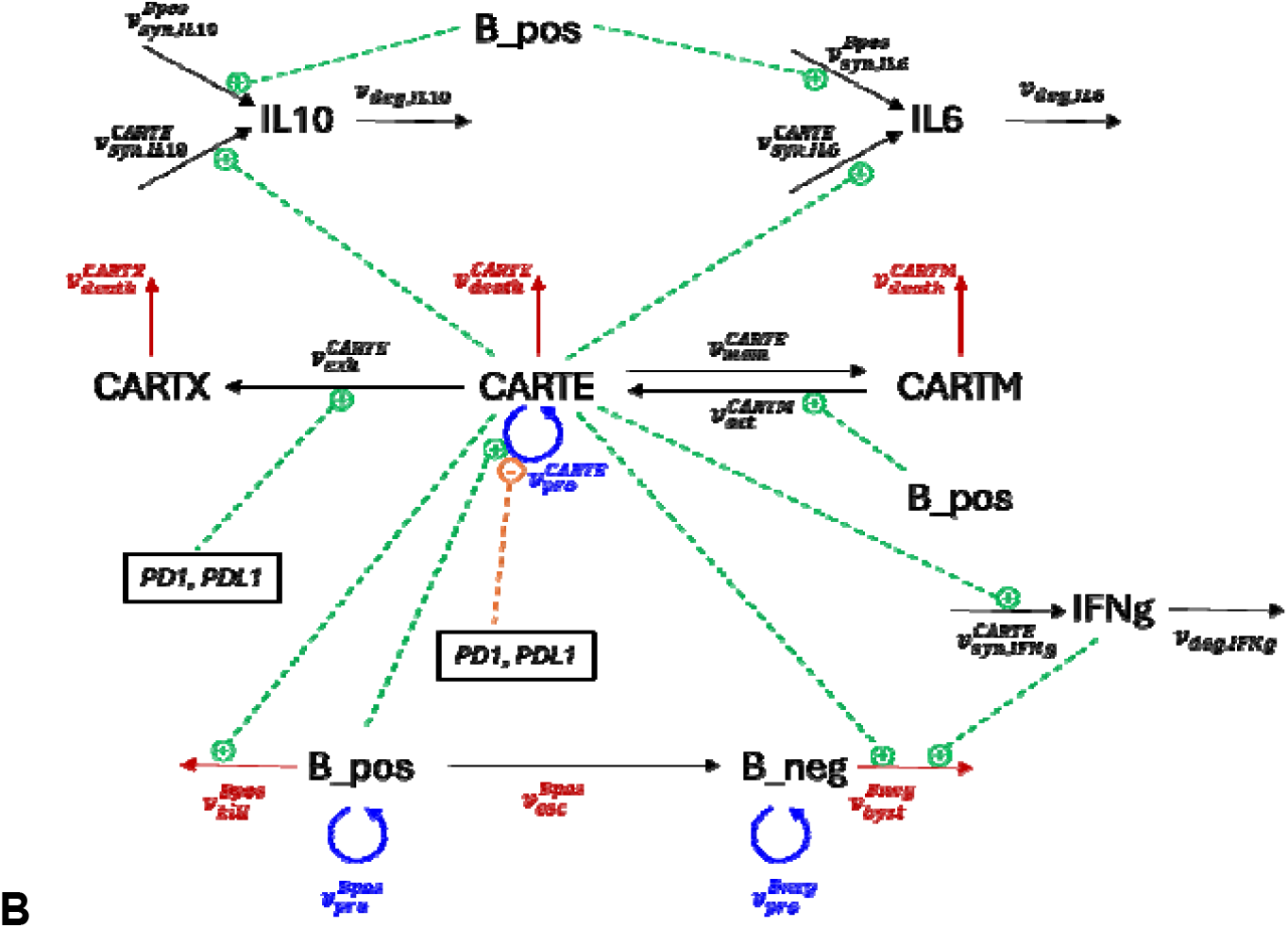
Structural representation of AI-generated and expert-corrected CAR-T QSP models. (A) First AI-generated model update (AIupdate1) incorporating mechanisms of T-cell exhaustion, PD-1 checkpoint regulation, and tumor antigen escape. (B) Expert-corrected model implementation (corrected_AIupdate1) after resolution of SBML structural inconsistencies, including explicit reaction definitions and parameter annotations.

**Figure 4.**
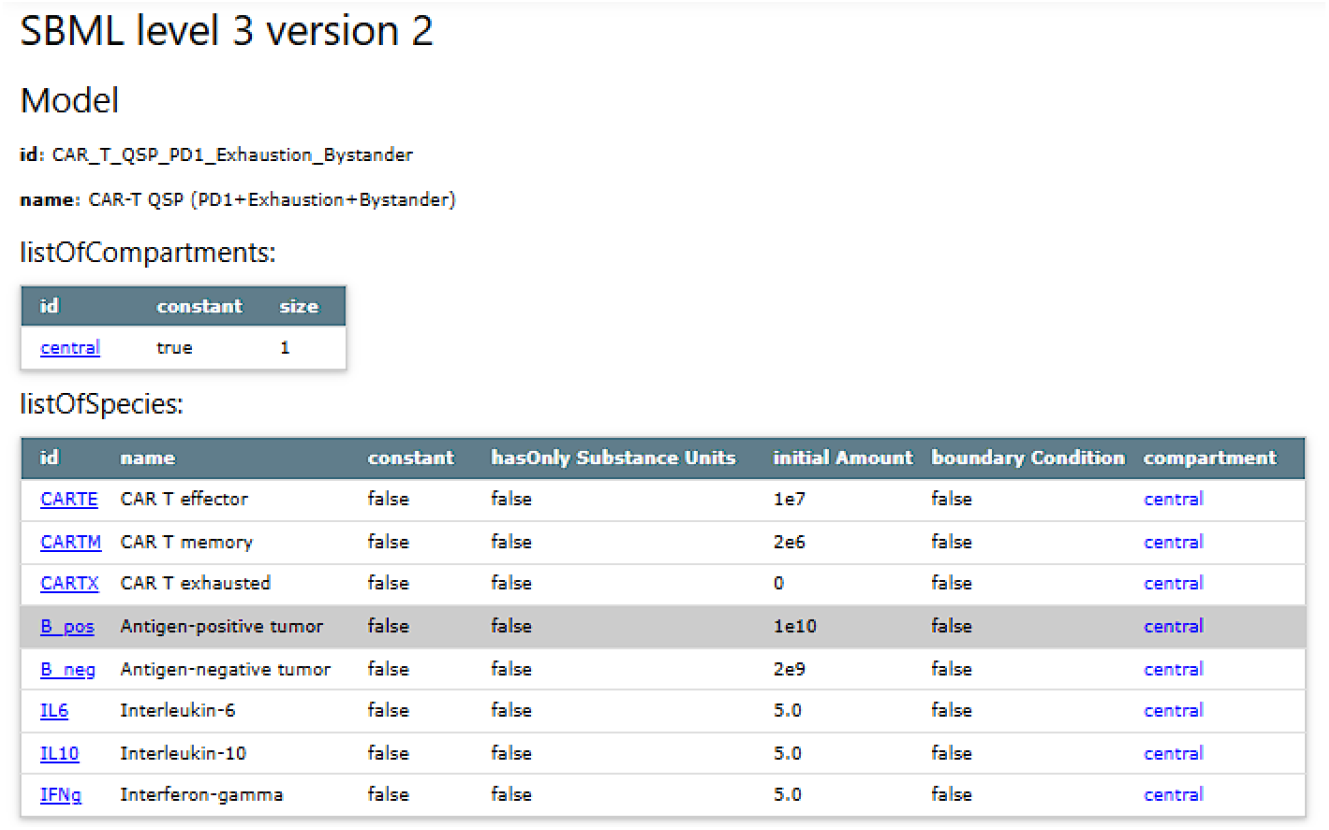
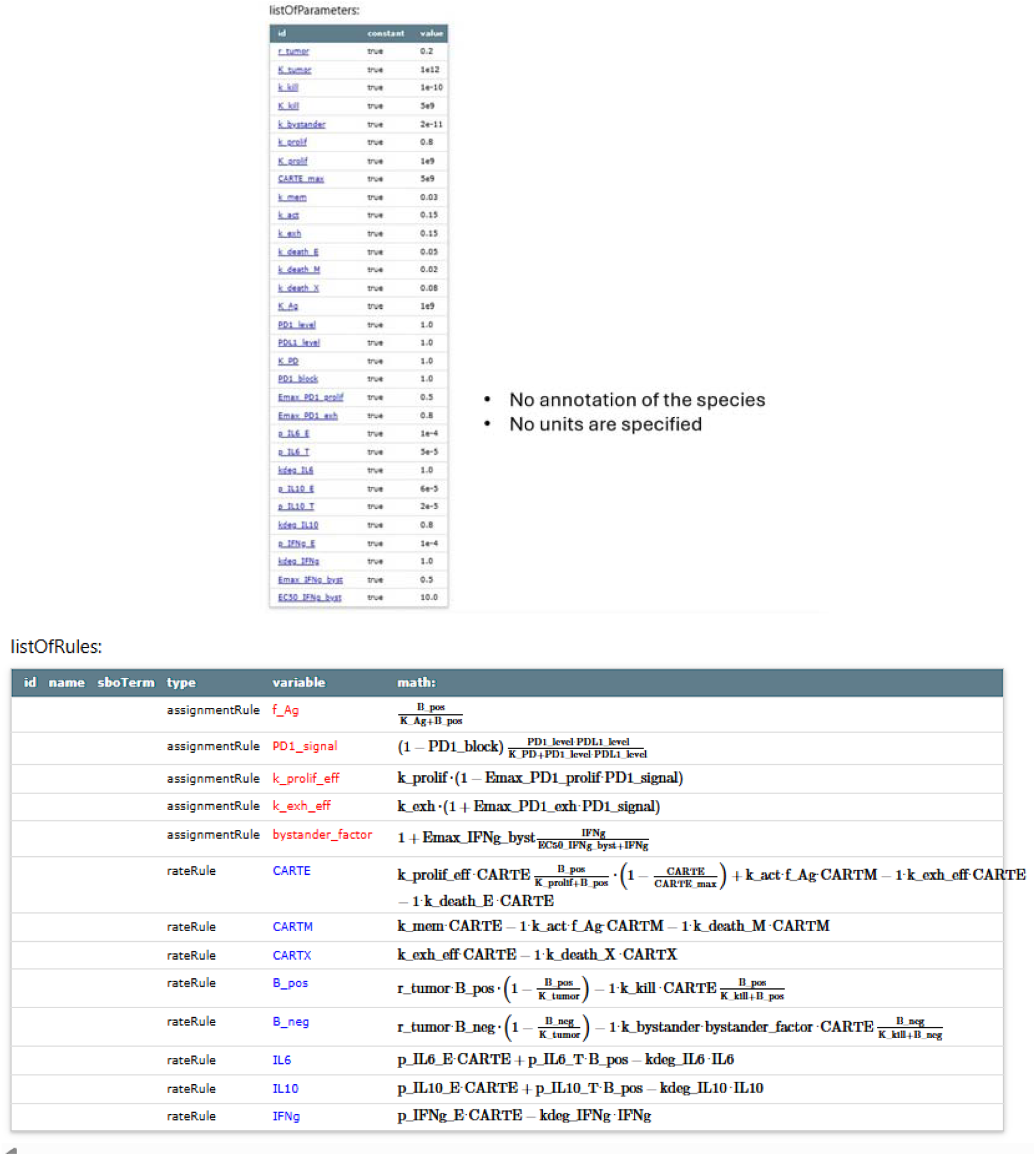
SBML representation of the AI-generated CAR-T QSP model update (“AIupdate1”). Selected fragments of the Systems Biology Markup Language (SBML) implementation generated by the AI-QSP framework are shown to illustrate the structure of the automatically produced model. (A) Definition of model compartments, species, and parameters introduced during the AI-assisted update, including exhausted CAR-T cells (CARTE_EX) and antigen-negative tumor cells (B_neg). (B) Example reaction declarations implementing key biological processes such as CAR-T proliferation, exhaustion transitions, and tumor antigen escape. (C) Corresponding kinetic rate-law expressions defining the mathematical representation of the biological interactions.

Inspection of the SBML code of the **AIupdate1** model revealed that the framework was able to introduce the intended biological mechanisms, but the initial implementation contained several technical inconsistencies. These included incomplete annotation of compartments, species, and parameters; absence of explicit stoichiometric process definitions for some reactions; ambiguous mapping between rate expressions and biological processes; and an inconsistency in the CARTE-to-CARTM transition, which was represented in the equation for **CARTM** but not consistently accounted for in the equation for **CARTE**. These observations indicate that the AI-QSP workflow could propose biologically relevant model extensions, but that expert validation remained necessary to ensure technical correctness and SBML consistency.

All identified technical deficiencies in the **AIupdate1** model were manually corrected and reimplemented in Heta format, followed by conversion into SBML using the Heta compiler. This yielded the **corrected_AIupdate1** model, shown in **Figure 3B** and **Figure 5**. The corrected model served as the expert-validated reference for the next iteration of the AI-QSP workflow.

**Figure 5.**
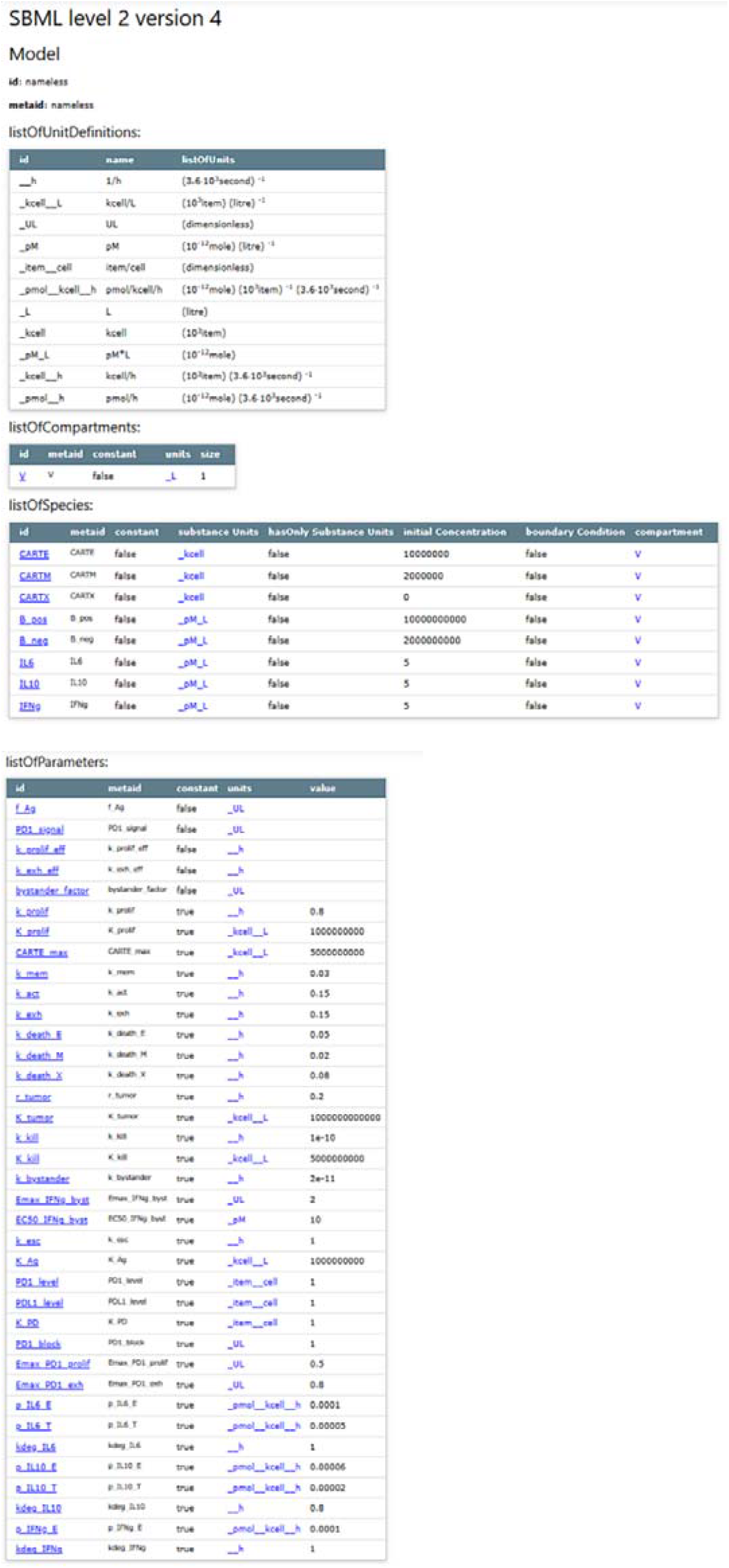
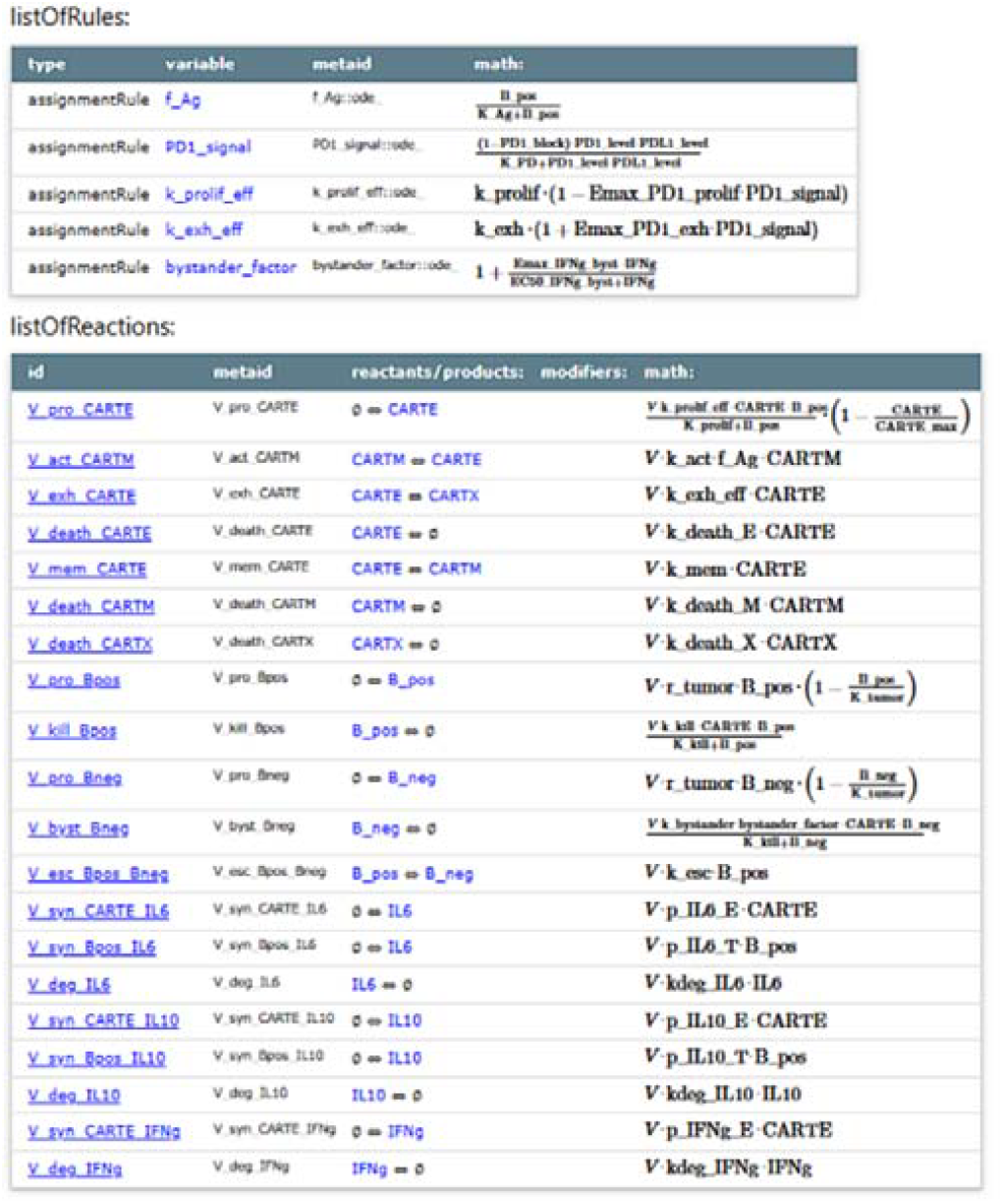
Expert-corrected SBML implementation of the CAR-T QSP model (“corrected_AIupdate1”). Representative fragments of the SBML code after expert validation and correction. (A) Revised species and parameter declarations ensuring dimensional consistency and explicit model annotations. (B) Corrected reaction definitions implementing CAR-T proliferation, exhaustion transitions, and tumor killing processes. (C) Corresponding kinetic rate laws defining the final mathematical formulation used for simulation and calibration.

**Figure 6.**
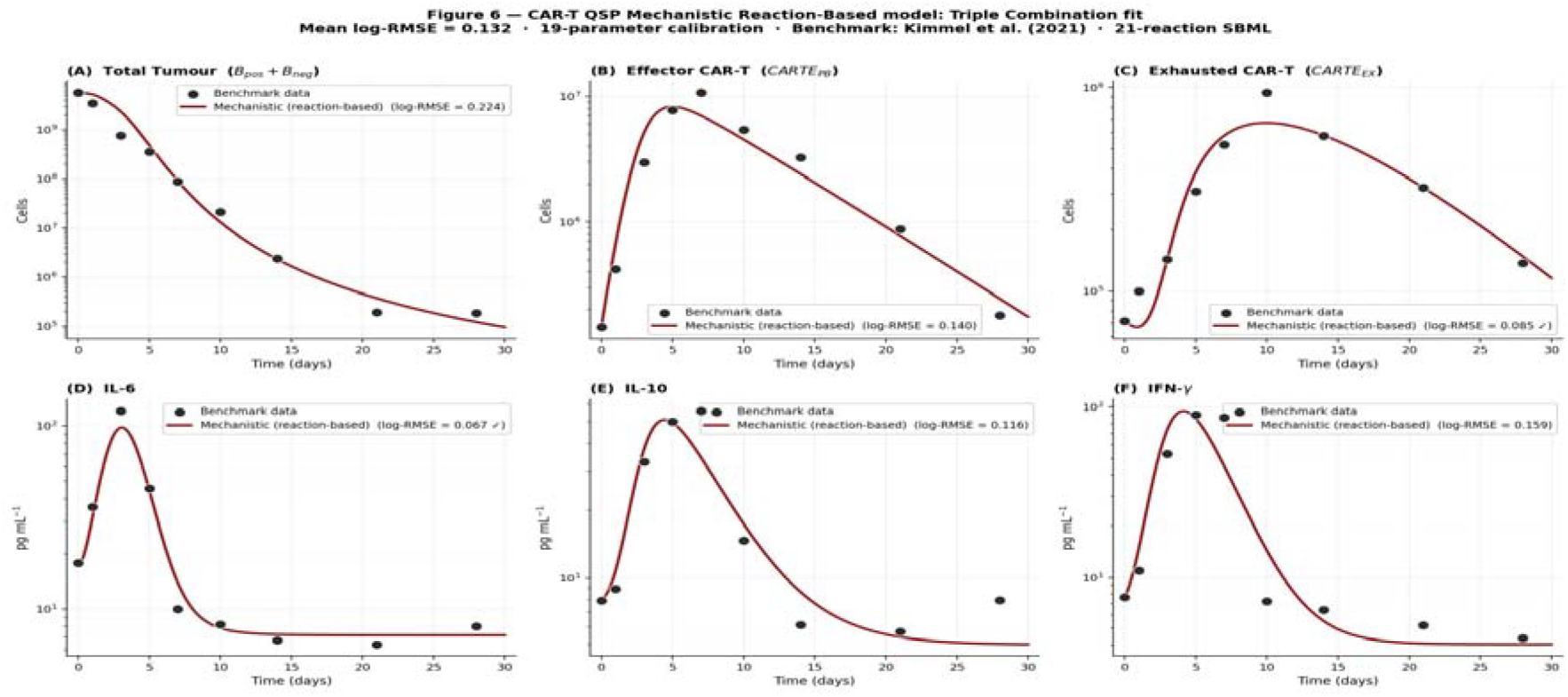
Mechanistic Reaction-Based model calibration fit: six variables, Triple Combination scenario AIupdate3 model (21 SBML reactions R1–R21, 19-parameter calibration, MA v11 best-fit) trajectories for six simultaneously calibrated variables under the Triple Combination scenario (exhaustion R4, PD-1 relief rule, antigen escape R11, bystander killing R14). Panels: (A) Total Tumor (B□+B□, reactions R8–R14), (B) effector CAR-T (R1– R5), (C) exhausted CAR-T (R4, R7), (D) IL-6 (R15–R16), (E) IL-10 (R17–R18), (F) IFN-γ (R19–R20). Filled circles: digitised benchmark data (Kimmel et al., 2021 [32]). Solid lines: AIupdate3 best-fit trajectories. Per-panel log-RMSE values are shown in panel legends; □ denotes variables meeting the log-RMSE < 0.10 accuracy threshold. Mean log-RMSE = 0.132 across six variables. All axes: log□ □ scale.

Based on the observed deficiencies, the prompts and workflow instructions driving the AI-QSP framework were refined. After this refinement step, the framework generated a **AIupdate2** model whose structure and code were consistent with the manually corrected implementation. This result demonstrates that the iterative expert-in-the-loop workflow improved the technical quality of the AI-generated mechanistic model and enabled convergence toward an expert-validated updated model structure.

### 3.3 Structural analysis of the updated CAR-T QSP model

Comparison of the final **AIupdate2** model with the original **AIcore** model showed that the AI-QSP workflow introduced biologically meaningful structural changes consistent with the requested mechanisms. The key structural extensions were the explicit representation of an exhausted CAR-T state, incorporation of antigen-negative tumor cells, and regulatory processes associated with checkpoint modulation and antigen escape. These additions expanded the mechanistic scope of the model beyond the original core representation and enabled simulation of more realistic CAR-T treatment dynamics under resistance conditions.

At the variable level, the updated model differed from the original model by introducing new state variables representing exhausted CAR-T cells and antigen-negative tumor cells, while retaining the shared variables describing effector and memory CAR-T cells, antigen-positive tumor cells, and cytokine concentrations. At the process level, the updated model introduced transitions between effector and exhausted CAR-T states and between antigen-positive and antigen-negative tumor states. In contrast, some features of the original core model, such as tissue migration processes, were not retained in the same form in the updated implementation.

Comparison of the rate-law structure further showed that the updated model incorporated mechanistic terms for exhaustion, antigen escape, and bystander-mediated killing, while preserving the general structure of proliferative and cytotoxic processes from the original model. Together, these observations indicate that the AI-QSP workflow successfully transformed the original core model into an extended mechanistic representation capable of describing key resistance mechanisms relevant to CAR-T therapy.

### 3.4 Calibration of the updated model under the Triple Combination scenario

The ability of AI-QSP to calibrate a QSP model was explored in accordance with workflow presented at Fig 1. Initially, updated version of QSP model **AIupdate2** was applied for calibration against synthetic dataset. However, we have failed to calibrate the version of the model, potentially, due to some structural limitations of implemented mechanisms. Then, we have allowed AI-QSP to adaptively change right hand sides of the model in such a way to allow appropriate calibration against chosen synthetic datasets. As a result we have come to **AIupdate3** version of the QSP model. This version is described in Supplementary materials S1. The calibrated performance of the updated model **AIupdate3** was evaluated using the **Triple Combination scenario**, which simultaneously included CAR-T exhaustion, PD-1 checkpoint blockade, antigen escape, and bystander-mediated killing. This integrated scenario was chosen as the most stringent test of the AI-QSP framework because it combines the principal biological mechanisms expected to shape CAR-T therapeutic response and resistance. Figure 6 illustrates the fitted tumor and CAR-T trajectories, while Table 2,3 summarizes the parameter estimates obtained from the automated optimization procedure. Using synthetic benchmark data generated from the **AIcore** model over a 28-day (nine time-point) time horizon, the AI-QSP framework performed automated parameter optimization of the updated model **AIupdate3** (21 reactions, R1–R21) using a 19-parameter multi-phase (comprising 7 primary kinetic parameters listed in Table 2,3, plus 12 cytokine subsystem parameters) L-BFGS-B strategy. The fitted model reproduced the benchmark dynamics for total tumor burden, circulating effector CAR-T cells, and cytokine concentrations with high fidelity across the 28-day post-infusion window. Visual comparison of model trajectories and digitised benchmark observations [32] showed close agreement, with the AIupdate3model achieving a mean log-RMSE of 0.132 across six simultaneously fitted variables. Per-variable log-RMSE values are reported in the Goodness-of-Fit row of Table 2, 3. Two variables — exhausted CAR-T cells (log-RMSE = 0.085) and IL-6 (log-RMSE = 0.067) — met the pre-specified accuracy threshold of log-RMSE < 0.10.

**Table 2.** Model species, parameters, initial conditions, and units (SBML representation)

| Category | Symbol / ID | Description | Initial Value | Unit |
| --- | --- | --- | --- | --- |
| Species |  |  |  |  |
| Tumor | B_pos | Antigen-positive tumor cells | $1 \times 10^6$ | cells |
| Tumor | B_neg | Antigen-negative (escape) tumor cells | $1 \times 10^6$ | cells |
| CAR-T Cells | CARTE | Effector CAR-T cells | $1 \times 10^6$ | cells / $\mu\text{L}$ |
| CAR-T Cells | CARTM | Memory CAR-T cells | $1 \times 10^6$ | cells / $\mu\text{L}$ |
| CAR-T Cells | CARTE_EX | Exhausted CAR-T cells | $1 \times 10^6$ | cells / $\mu\text{L}$ |
| Cytokines | IL-6 | Interleukin-6 | 1.0 | relative units |
| Cytokines | IL-10 | Interleukin-10 | 0.5 | relative units |
| Cytokines | IFN-γ | Interferon-γ | 1.5 | relative units |
| Drug | aPD1 | Anti-PD-1 checkpoint inhibitor | 0<br>(variable) | $\mu\text{g} / \text{mL}$ |
| Kinetic Parameters |  |  |  |  |
| Tumor Growth | r_Bpos | Growth rate of B_pos cells | Fitted | $\text{day}^{-1}$ |
| Tumor Growth | r_Bneg | Growth rate of B_neg cells | Fitted | $\text{day}^{-1}$ |
| Tumor Killing | k_kill | CAR-T-mediated killing of B_pos | Fitted | day<sup>-1</sup> |
| Tumor Killing | k_byst | Bystander killing of B_neg by CARTE | Fitted | day<sup>-1</sup> |
| Tumor Escape | k_loss | Antigen loss rate (B_pos → B_neg) | Fitted | day<sup>-1</sup> |
| CAR-T Prolif. | k_prolif | CARTE expansion rate constant | Fitted | day<sup>-1</sup> |
| CAR-T Prolif. | $K_{\text{prolif}}$ | Half-saturation for tumor-driven proliferation | Fixed | cells |
| CAR-T Death | k_death | Natural CARTE death rate | Fitted | day<sup>-1</sup> |
| CAR-T Exhaust. | k_exh | Rate of CARTE → CARTE_EX conversion | Fitted | day<sup>-1</sup> |
| CAR-T Memory | k_mem | CARTE → CARTM conversion rate | Fixed | day<sup>-1</sup> |
| Checkpoint | E_max_PD1 | Max fractional reduction of $k_{\text{exh}}$ by aPD1` | Fitted | dimensionless |
| Checkpoint | IC_50 | aPD1 concentration for | Fixed | µg / mL |
|  |  | 50% effect |  |  |
| Simulation Settings |  |  |  |  |
|  | t_end | Simulation duration | 28 | days |
|  | dt | Integration time step | 0.01 | days |

**Table 3.**
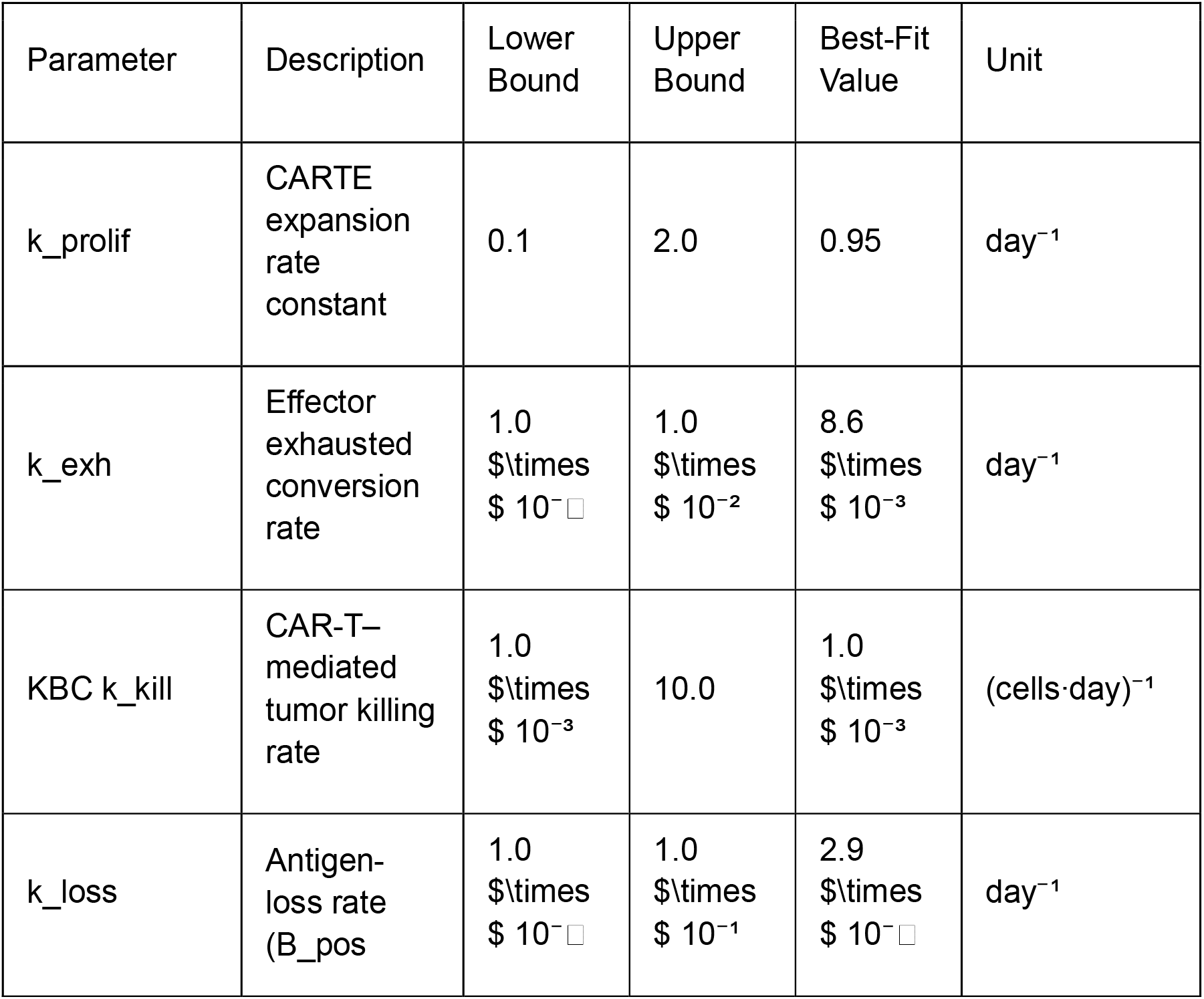
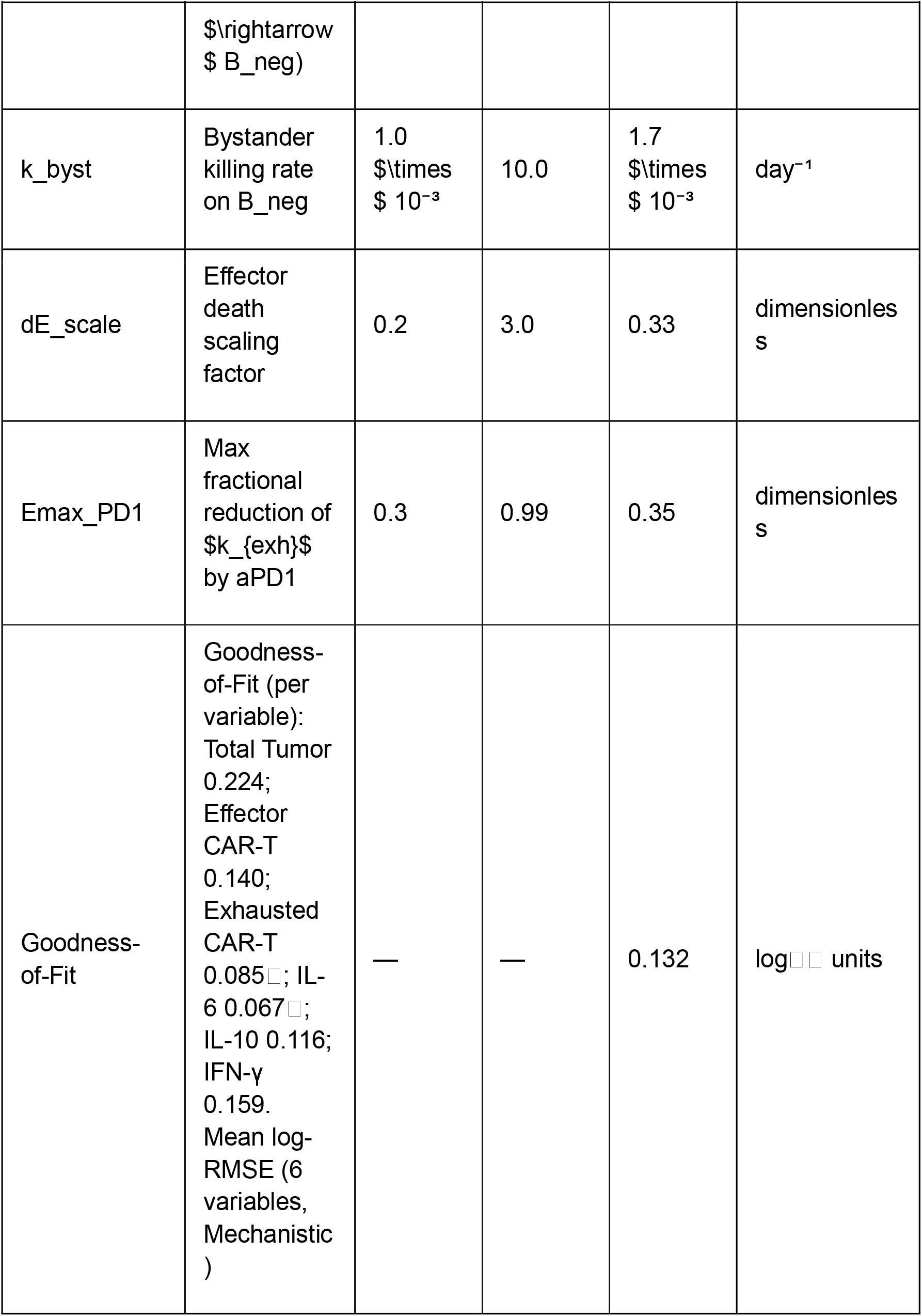
Best-fit parameter estimates — Mechanistic Reaction-Based model (19-parameter)

| Parameter | Description | Lower Bound | Upper Bound | Best-Fit Value | Unit |
| --- | --- | --- | --- | --- | --- |
| k_prolif | CARTE expansion rate constant | 0.1 | 2.0 | 0.95 | day<sup>-1</sup> |
| k_exh | Effector exhausted conversion rate | 1.0 $\times 10^{-4}$ | 1.0 $\times 10^{-2}$ | 8.6 $\times 10^{-3}$ | day<sup>-1</sup> |
| KBC k_kill | CAR-T-mediated tumor killing rate | 1.0 $\times 10^{-3}$ | 10.0 | 1.0 $\times 10^{-3}$ | (cells·day)<sup>-1</sup> |
| k_loss | Antigen-loss rate (B_pos) | 1.0 $\times 10^{-4}$ | 1.0 $\times 10^{-1}$ | 2.9 $\times 10^{-4}$ | day<sup>-1</sup> |
| | $\rightarrow B_{neg}$ | | | | |
| k_byst | Bystander killing rate on $B_{neg}$ | 1.0<br>$\times 10^{-3}$ | 10.0 | 1.7<br>$\times 10^{-3}$ | day<sup>-1</sup> |
| dE_scale | Effector death scaling factor | 0.2 | 3.0 | 0.33 | dimensionless |
| E<sub>max</sub>_PD1 | Max fractional reduction of $k_{exh}$ by aPD1 | 0.3 | 0.99 | 0.35 | dimensionless |
| Goodness-of-Fit | Goodness-of-Fit (per variable):<br>Total Tumor 0.224;<br>Effector CAR-T 0.140;<br>Exhausted CAR-T 0.085; IL-6 0.067; IL-10 0.116; IFN-γ 0.159.<br>Mean log-RMSE (6 variables, Mechanistic) | — | — | 0.132 | log units |

The optimised **AIupdate3** model captured the characteristic behavior expected under the Triple Combination configuration. Tumor burden declined in response to CAR-T expansion mediated by mass-action killing (R10) and bystander cytotoxicity on antigen-escape cells (R14). Longer-term dynamics reflected the competing effects of exhaustion (R4, PD-1 modulated), checkpoint rescue (PD1_relief assignment rule), and antigen escape (R11). The CAR-T trajectory reproduced the expected expansion–contraction pattern, and cytokine dynamics (IL-6, IL-10, IFN-γ; reactions R15–R20) remained consistent with the timing and magnitude of the benchmark immune response.

### 3.5 Quantitative goodness-of-fit

Quantitative agreement between the calibrated model and the synthetic benchmark data was assessed using the log-transformed root mean square error (log-RMSE). The final calibrated solution achieved a mean log-RMSE of 0.132 (AIupdate3 model). Per-variable log-RMSE values ranged from 0.067 (IL-6, ⍰) to 0.224 (total tumor burden), as detailed in Table 3. Two variables did not meet the pre-specified accuracy threshold: IFN-γ (log-RMSE = 0.159) and total tumour burden (log-RMSE = 0.224). The relatively higher error for total tumour burden reflects a structural trade-off between the antigen-escape rate (R11) and bystander killing (R14) that limits simultaneous optimisation of both early decline and late-phase dynamics. The IFN-γ fit (log-RMSE = 0.159) is constrained by the semi-decoupled cytokine source term structure in the current model; a more flexible IFN-γ production rate law is identified as a direction for AIupdate4. These are documented as directions for future model extension.

Taken together, these results demonstrate that the AI-QSP framework can:

1. reconstruct and extend a published CAR-T QSP model,
2. iteratively improve the technical correctness of AI-generated model structure through expert feedback, and
3. calibrate the updated model to reproduce the dynamics of the original reference model under a challenging integrated mechanistic scenario. This supports the feasibility of AI-assisted model evolution as a potential workflow for mechanistic pharmacology modeling.

3.6 A Sobol’ variance-based GSA was performed on all 19 calibrated parameters using a Saltelli quasi-random sample of N = 1024 base vectors (40,960 total model evaluations; Supplementary S3). First-order (S_1_) and total-order (S_T_) Sobol’ indices were computed using the Jansen (1999) estimator with 95% bootstrap confidence intervals (n = 500 replicates). The GSA fulfills the model-robustness requirement of ASME V&V 40 by quantifying output sensitivity to individual parameters and their interactions, and supports principled model reduction by identifying five non-influential parameters (S_T_ < 0.05 across all outputs) that were fixed at best-fit values for the subsequent profile-likelihood analysis. Key findings: tumor burden was most sensitive to k_kill_, k_byst_, and K_DX_; cytokine outputs were dominated by their respective production (α) and degradation (k_deg_) parameters. Full numerical results, heat-map visualizations, and reproducibility materials are provided in Supplementary S3.

3.7 Following GSA-guided model reduction, profile-likelihood identifiability was assessed for the 14 active parameters using single-pass marginal profiles evaluated across each parameter’s full bound range (Supplementary S4). Two objective-function thresholds were applied: ΔJ = 0.010 (inner confidence interval) and ΔJ = 0.020 (outer confidence interval), yielding three identifiability classes. Ten of 14 parameters were classified as Class A (identifiable), with finite bounded confidence intervals; four parameters were Class C (non-identifiable): k_byst_, K_DX_, α_IL6_, and k_prolif_. These represent structural limitations of single-dataset calibration and are designated as priority targets for Bayesian hierarchical re-estimation from clinical data in Stage 2. An identifiability rate of 71% (10/14) for a 21-reaction model calibrated from a single benchmark dataset is consistent with published expectations for QSP models of comparable complexity. Confidence interval statistics and per-parameter classifications are provided in Supplementary S4 (Table S4.1) and visualised in Figure 7.

**Figure 7.**
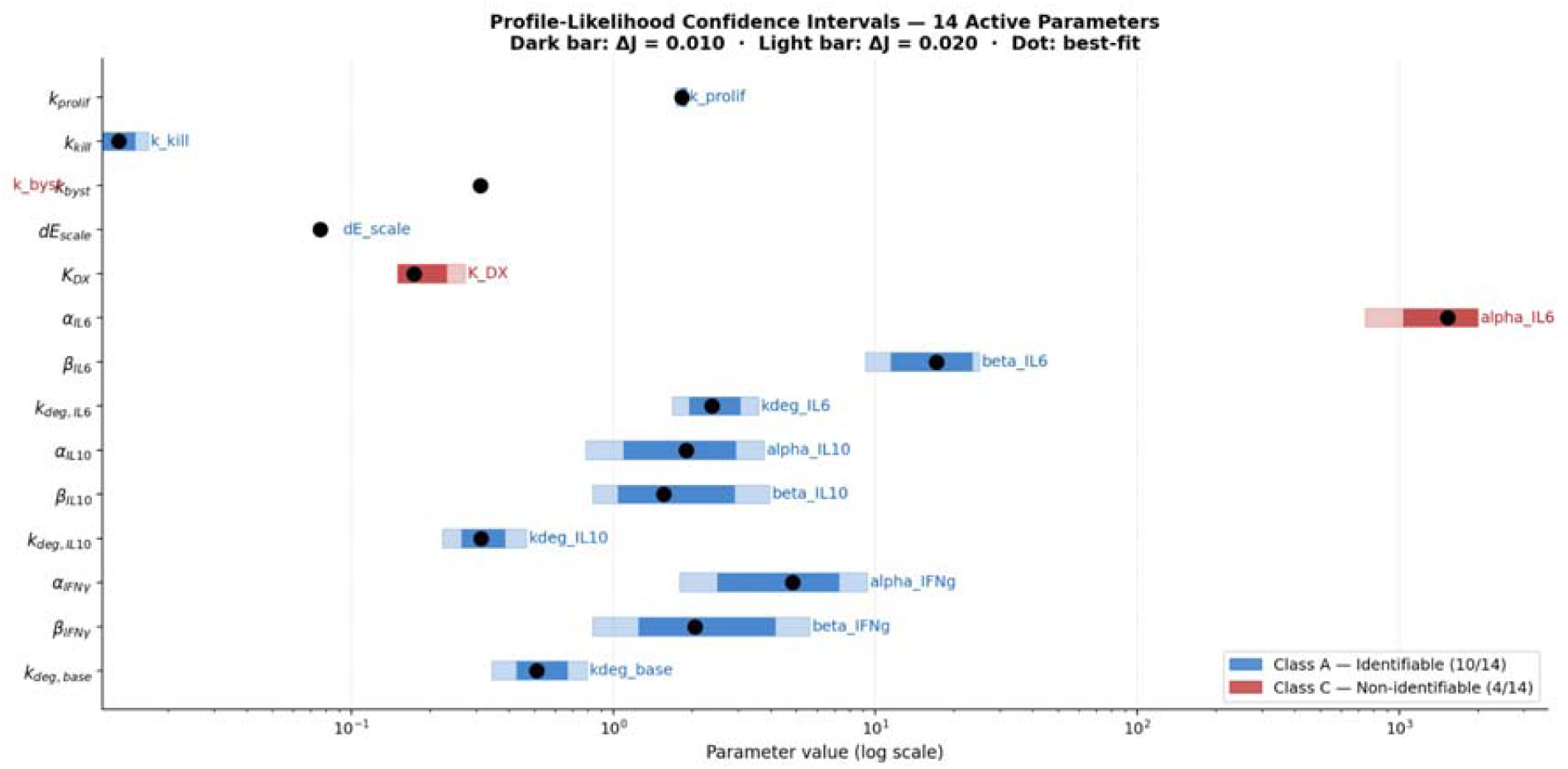
Profile-Likelihood Confidence Intervals — 14 Active Parameters. Dark bar: ΔJ = 0.010 (inner CI); light bar: ΔJ = 0.020 (outer CI); dot: best-fit value (log scale). Blue: Class A — Identifiable (10/14); red: Class C — Non-identifiable (4/14). Parameters k_byst, K_DX, α_IL6, and k_prolif were classified as Class C, representing priority targets for Bayesian re-estimation from clinical data. Source data: Supplementary S4 (Table S4.1).

## 4. Discussion

This study presents a prototype **AI-assisted quantitative systems pharmacology (AI-QSP) framework** designed to support automated reconstruction, extension, and calibration of mechanistic pharmacology models. The primary objective of the work was to evaluate whether such a framework can reliably transform a literature-derived model into an updated mechanistic representation and reproduce the dynamics of the reference system through automated calibration.

To test this capability, the workflow was applied to a published CAR-T cell therapy QSP model. The AI-QSP framework first reconstructed the literature model into a standardized SBML representation (AIcore). It then generated structural model updates based on biologically motivated prompts describing key resistance mechanisms, including T-cell exhaustion, checkpoint regulation through PD-1/PD-L1 signaling, and tumor antigen escape. These mechanisms represent widely recognized drivers of treatment resistance in CAR-T therapy and therefore provide a biologically meaningful test case for automated model evolution.

The results demonstrate that the AI-QSP workflow can successfully generate mechanistic model extensions and iteratively improve the technical correctness of the resulting SBML model through an expert-in-the-loop validation process. The first AI-generated model update captured the intended biological processes but contained several technical inconsistencies typical of automatically generated code. After refinement of the prompts and correction of these issues, the second AI-generated model version reproduced the expert-validated implementation. This iterative improvement highlights an important property of the proposed framework: the ability to combine automated model generation with expert validation to progressively converge toward a technically correct and biologically consistent mechanistic model.

The calibration phase of the study revealed an important practical insight: the first structurally extended model, AIupdate2, could not be successfully calibrated against the synthetic benchmark data. Calibration of AIupdate2 converged to poor solutions, likely due to structural limitations in the right-hand side rate expressions introduced during the AI-guided extension step — in particular, parameter interactions between the exhaustion transition (Rxn_CARTE_Exhaustion) and the PD-1 relief assignment rule created near-redundant contributions that rendered the objective function landscape ill-conditioned. This failure is itself an informative outcome: it demonstrates that AI-generated structural extensions, even when technically correct at the SBML level, may not be immediately calibratable and require further adaptive refinement. In response, the AI-QSP framework was re-engaged to adaptively modify the rate expressions of the problematic processes, producing AIupdate3. This version incorporates revised kinetic formulations for the exhaustion and bystander killing pathways (detailed in Supplementary S1) that improve the conditioning of the optimisation landscape. AIupdate3 was then successfully calibrated under the “Triple Combination scenario” — simultaneously incorporating exhaustion, checkpoint blockade, antigen escape, and bystander-mediated cytotoxicity. This integrated scenario represents the most stringent test of the framework because it activates all four resistance mechanisms simultaneously. The automated parameter optimisation recovered the benchmark dynamics with close agreement across tumour burden, CAR-T cell kinetics, and cytokine responses (mean log-RMSE = 0.132; sum log-RMSE = 0.792 across six outputs), demonstrating that the iterative AI-QSP refinement cycle can navigate initial calibration failures and converge on a quantitatively consistent solution.

From a methodological perspective, the key contribution of this work is the demonstration that **AI-assisted model evolution can be integrated into a reproducible systems pharmacology workflow**. Traditional QSP model development relies heavily on manual interpretation of biological literature and iterative model modification by domain experts. While this process produces highly detailed models, it is labor-intensive and often difficult to reproduce across modeling teams. The AI-QSP framework addresses this limitation by introducing an automated knowledge-interpretation layer capable of translating textual biological descriptions into candidate model components. When combined with standardized SBML representations and automated calibration tools, this approach provides a scalable strategy for maintaining and updating complex mechanistic models as new biological knowledge becomes available.

The framework also aligns with current regulatory expectations for Model-Informed Drug Development (MIDD) workflows. Regulatory agencies such as the U.S. Food and Drug Administration and the European Medicines Agency emphasize transparency, reproducibility, and traceability in computational models used for drug development. By encoding all models in SBML and maintaining a structured record of model updates, calibration datasets, and parameter estimates, the AI-QSP workflow facilitates the generation of auditable modeling pipelines that could support regulatory-grade applications in the future.

### 4.1 Limitations and future work

Several limitations of the present study should be acknowledged. First, the validation experiments were performed using **synthetic benchmark data** generated from the reference model. This design provides a controlled test of the computational workflow because the true system dynamics are known. However, it does not directly assess the biological predictive performance of the updated model. Application of the framework to independent experimental or clinical datasets will be necessary to evaluate its ability to generate biologically predictive models.

Second, although the AI-QSP framework can propose structural model updates, the current implementation still requires expert validation to ensure technical correctness and biological plausibility. This expert-in-the-loop design is intentional and reflects the current state of mechanistic modeling, where domain expertise remains essential. Future work may explore improved automated validation procedures, including rule-based consistency checks, SBML validation tools, and formal model-verification approaches.

Finally, the present study focuses on a relatively compact CAR-T QSP model. Applying the framework to larger multiscale systems pharmacology models involving multiple tissues, cell populations, and signaling pathways will provide a more comprehensive evaluation of its scalability. The proposed AI-QSP framework complements recent efforts to integrate artificial intelligence with mechanistic pharmacology modeling and may accelerate model-informed drug development workflows for emerging cell and gene therapies.

### 4.2 Regulatory Compliance and Model Credibility Assessment

The present study was developed with regulatory transparency as a primary design objective. The complete regulatory and compliance documentation is distributed across four supplementary materials: Supplementary S1 provides the full mathematical specification of the reaction-based SBML model (21 reactions, 9 species, 19-parameter calibration); Supplementary S2 presents the structured Model Credibility Assessment, completed ICH M15 Assessment Table, and MIDD submission roadmap; Supplementary S3 reports the Global Sensitivity Analysis (Sobol’, N = 1024); and Supplementary S4 details the Profile-Likelihood identifiability analysis for 14 active parameters. Together, these documents constitute the evidentiary package required for regulatory evaluation of the model at the current exploratory development stage.

The ICH M15 Guideline on General Principles for Model-Informed Drug Development [33] provides the overarching harmonized framework for MIDD evidence assessment across FDA, EMA, and other ICH member regions. Under M15, model evaluation requirements are calibrated to Model Risk — defined as the combination of Model Influence (the intended weight of model outcomes in decision-making) and Consequence of Wrong Decision. For the present pre-clinical prototype, Model Risk is assessed as Low-to-Medium: model influence is low, as simulation outputs supplement rather than replace clinical evidence at this development stage, and the consequence of a wrong decision is limited given the exploratory context. Model Impact is rated as Medium, reflecting the novel AI-assisted model evolution approach that departs from conventional manual QSP workflows. The completed M15 Assessment Table, including pre-specified technical criteria and outcome of the MIDD evidence assessment, is provided in Supplementary S2. The technical criteria defined a priori were: mean log-RMSE ≤ 0.15 across all fitted outputs; structural identifiability of ≥ 70% of active parameters by profile-likelihood; and GSA-confirmed parameter relevance (total-order Sobol’ index S_T_ ≥ 0.05 for at least one output). All three criteria were satisfied by the MA v11 calibration solution (mean log-RMSE = 0.132; 10/14 parameters Class A identifiable; 14/19 parameters with S_T_ ≥ 0.05; see Supplementary S3).

Model credibility was assessed following the risk-informed verification and validation framework of ASME V&V 40-2018, as detailed in Supplementary S2. Verification confirmed that the SBML model (Level 3 Version 2) correctly implements the mathematical equations documented in Supplementary S1, with mass-balance and stoichiometry validated programmatically using libSBML and Tellurium. Validation comprised three components: (i) goodness-of-fit; (ii) global sensitivity analysis establishing the relative influence of all parameters across all outputs (Supplementary S3); and (iii) profile-likelihood identifiability analysis characterizing practical parameter uncertainty for the 14 active parameters (Supplementary S4). The unified parameter credibility table integrating all three evidence streams is provided in Supplementary S2 (Table S2.4).

A structured roadmap for progression from the current pre-clinical prototype to a regulatory-submission-ready MIDD package is provided in Supplementary S2. The roadmap identifies three development stages aligned with ICH M15 and ASME V&V 40. Stage 1 (current work, complete): pre-clinical proof of concept with synthetic benchmark data, SBML model encoding, GSA, and profile-likelihood identifiability — all Stage 1 deliverables are documented in Supplementary S1–S4. Stage 2: integration of patient-level clinical PK/PD and biomarker data with Bayesian hierarchical parameter estimation to resolve the four Class C non-identifiable parameters; preparation of a Model Analysis Plan (MAP) per ICH M15 Section 4.1. Stage 3: external validation against independent clinical cohorts, full Model Analysis Report (MAR) per ICH M15 Section 4.2, and preparation of regulatory submission documentation in Common Technical Document format.

## 5 Conclusions

In summary, this work demonstrates the feasibility of integrating artificial intelligence with mechanistic pharmacology modeling through an **AI-QSP workflow** capable of reconstructing, extending, and calibrating QSP models in a reproducible computational environment. By combining large language model–based knowledge interpretation with SBML-based modeling tools and automated optimization, the framework provides a structured approach for iterative model evolution while preserving mechanistic transparency. These capabilities position AI-QSP as a promising enabling technology for scalable, reproducible modeling workflows in systems pharmacology and model-informed drug development.

