## Supplementary File 1 for "An AI-Assisted Workflow for Reconstruction, Extension, and Calibration of Quantitative Systems Pharmacology Models"

**Supplementary Material S1**

**Mathematical Equations of the CAR-T QSP Mechanistic Reaction-Based Model**

*SBML source: CAR_T_QSP_triple_true_mechanistic_reaction_based.xml · 21 reactions, 9 species, 1 assignment rule · 19-parameter calibration (MA v11 best-fit)*

### **S1.1 Model Overview**

The Mechanistic Reaction-Based CAR-T QSP model encodes the dynamics of nine state variables as 21 explicit SBML reactions together with one algebraic assignment rule. Unlike the Rule-Based formulation (which uses rate rules), each biological process is represented as a stoichiometrically defined reaction with an explicit kinetic law, improving biological transparency and facilitating SBML-level analysis and exchange. The model was encoded in SBML Level 3 Version 2 and calibrated against digitised benchmark data (Kimmel et al., 2021) using a 19-parameter multi-phase L-BFGS-B optimisation strategy. The best-fit solution (MA v11) achieved a mean log-RMSE of **0.132** across six simultaneously fitted variables.

### **S1.2 State Variables**

Table S1 lists the nine SBML *species* with biological descriptions and calibration initial conditions (ICs) from the digitised benchmark (Kimmel et al., 2021).

| **SBML ID** | **Symbol** | **Description** | **IC (calibration)** | **Units** | **Reactions** |
| --- | --- | --- | --- | --- | --- |
| CARTE | C_E | Effector CAR-T cells (peripheral blood) | 1.451×10⁵ cells | cells | R1–R4, R5 |
| CARTM | C_M | Memory CAR-T cells | 1.451×10⁴ cells | cells | R3, R5–R6 |
| CARTE_EX | C_X | Exhausted CAR-T cells (peripheral blood) | 7.135×10⁴ cells | cells | R4, R7 |
| B_pos | B⁺ | Antigen-positive tumour cells | 5.667×10⁹ cells | cells | R1, R5, R8–R11 |
| B_neg | B⁻ | Antigen-negative tumour cells (escape phenotype) | 5.667×10⁴ cells | cells | R11–R14 |
| IL6 | L₆ | Interleukin-6 concentration | 17.93 pg mL⁻¹ | pg mL⁻¹ | R15–R16 |
| IL10 | L₁₀ | Interleukin-10 concentration | 7.90 pg mL⁻¹ | pg mL⁻¹ | R17–R18 |
| IFNg | γ | Interferon-γ concentration | 7.65 pg mL⁻¹ | pg mL⁻¹ | R19–R20 |
| aPD1 | A | Anti-PD-1 antibody concentration (normalised) | 5.0 (norm.) | norm. | R21 |

### **S1.3 Fixed Parameters**

Fixed parameters are declared in the SBML *listOfParameters* with *constant="true"*. They are not subject to optimisation. The normalisation constants *S_C = C_E(0)* and *S_B = B⁺(0)* are derived from the benchmark initial conditions and encode the physiological cell-count scales used in the kinetic laws.

| **No.** | **Symbol** | **SBML ID** | **Description** | **Value** | **Units** |
| --- | --- | --- | --- | --- | --- |
| 1 | S_C | — | Effector CAR-T normalisation scale = C_E(0) | 1.451×10⁵ | cells |
| 2 | S_B | — | Tumour normalisation scale = B⁺(0) | 5.667×10⁹ | cells |
| 3 | s | scale_1e9 | Explicit 10⁹ scale in SBML kinetic laws | 1×10⁹ | cells |
| 4 | r_E | rE_per_1e9 | CAR-T effector mass-action expansion coeff. | 16.05 | d⁻¹ per 10⁹ cells |
| 5 | δ_E | dCARTE_rate | CAR-T effector & exhausted base death rate | 1.44 | d⁻¹ |
| 6 | α_M | aM_per_1e9 | Effector → memory differentiation rate coeff. | 0.024 | d⁻¹ per 10⁹ cells |
| 7 | r_M | rM | Memory reactivation rate (MM numerator) | 0.0509 | d⁻¹ |
| 8 | δ_M | dCARTM_rate | Memory CAR-T death rate | 0.408 | d⁻¹ |
| 9 | r_B | rB | Tumour B⁺ intrinsic growth rate | 0.0093 | d⁻¹ |
| 10 | δ_B | dB | Tumour B⁺/B⁻ natural death rate | 0.0076 | d⁻¹ |
| 11 | K_{BC} | KBC_per_1e9 | MM half-saturation for B⁺ in memory reactivation R5 | 0.19×10⁹ | cells |
| 12 | k_l | k_loss | Antigen-escape conversion rate B⁺ → B⁻ | 0.0002 | d⁻¹ |
| 13 | r_{Bn} | k_growth_neg | B⁻ (escape tumour) growth rate | 0.002 | d⁻¹ |
| 14 | k_b | k_bystander_per_1e9 | Bystander killing coeff. on B⁻ | 0.15 | d⁻¹ per 10⁹ cells |
| 15 | P_{IL6} | PIL6 | IL-6 production proportionality constant (SBML default) | 0.0018 | pg mL⁻¹ d⁻¹ per 10⁹ cells |
| 16 | P_{IL10} | PIL10 | IL-10 production proportionality constant (SBML default) | 0.0007 | pg mL⁻¹ d⁻¹ per 10⁹ cells |
| 17 | P_{IFNγ} | PINFg | IFN-γ production proportionality constant (SBML default) | 0.0003 | pg mL⁻¹ d⁻¹ per 10⁹ cells |
| 18 | k_{deg,L6} | kdeg_IL6 | IL-6 first-order degradation rate (SBML default) | 1.0 | d⁻¹ |
| 19 | k_{deg,L10} | kdeg_IL10 | IL-10 first-order degradation rate (SBML default) | 1.0 | d⁻¹ |
| 20 | k_{deg,γ} | kdeg_IFNg | IFN-γ first-order degradation rate (SBML default) | 1.0 | d⁻¹ |
| 21 | E_{max} | Emax_aPD1 | Maximum fractional PD-1 inhibition of exhaustion | 0.9 | — |
| 22 | IC₅₀ | IC50_aPD1 | aPD1 half-effect concentration | 5.0 | norm. units |
| 23 | k_{elim} | k_elim_aPD1 | Anti-PD-1 antibody elimination rate | 0.05 | d⁻¹ |

### **S1.4 Algebraic Assignment Rule**

The SBML *listOfRules* contains one algebraic assignment rule that computes the PD-1 checkpoint relief factor *f_{PD1}* (SBML variable *PD1_relief*) used in reaction R4:

| **SBML variable** | **Formula** | **Biological meaning** |
| --- | --- | --- |
| PD1_relief | f_{PD1} = 1 − E_max · A_{PD1} / (IC₅₀ + A_{PD1}) | PD-1 inhibition factor on exhaustion rate |

When *A_{PD1} → 0*, *f_{PD1} → 1* (no inhibition, maximal exhaustion). When *A_{PD1} ≫ IC₅₀*, *f_{PD1} → 1 − E_max*, reducing exhaustion by the maximum fraction.

### **S1.5 SBML Reactions (R1–R21)**

Table S3 lists all 21 reactions from the SBML *listOfReactions* with their stoichiometries and kinetic laws exactly as encoded in the SBML file. All reactions are irreversible (*reversible="false"*). Calibrated cytokine parameters (Table S4) override the SBML default values for reactions R15–R20 in the Python implementation.

| **ID** | **Reaction name** | **Reactants** |  | **Products** | **Kinetic law (SBML)** | **Biological process** |
| --- | --- | --- | --- | --- | --- | --- |
| **R1** | CARTE_expansion_antigen_driven | *CARTE + B_pos* | → | *2 CARTE + B_pos* | *r_E · (C_E / s) · B⁺* | Antigen-driven CAR-T proliferation (mass-action) |
| **R2** | CARTE_death | *CARTE* | → | *∅* | *δ_E · dE_scale · C_E* | Effector CAR-T constitutive death |
| **R3** | CARTE_to_memory | *CARTE* | → | *CARTM* | *α_M · (C_E / s)* | Effector → memory differentiation |
| **R4** | CARTE_exhaustion | *CARTE* | → | *CARTE_EX* | *k_{exh} · f_{PD1} · C_E* | PD-1 checkpoint–modulated exhaustion |
| **R5** | CARTM_reactivation_by_antigen | *CARTM + B_pos* | → | *CARTE + B_pos* | *r_M · C_M · B⁺ / (K_{BC} + B⁺)* | Memory reactivation (Michaelis-Menten) |
| **R6** | CARTM_death | *CARTM* | → | *∅* | *δ_M · C_M* | Memory CAR-T constitutive death |
| **R7** | CARTE_EX_death | *CARTE_EX* | → | *∅* | *K_{DX} · C_X* | Exhausted CAR-T clearance |
| **R8** | Bpos_growth | *B_pos* | → | *2 B_pos* | *r_B · B⁺* | Tumour B⁺ exponential growth |
| **R9** | Bpos_natural_death | *B_pos* | → | *∅* | *δ_B · B⁺* | Tumour B⁺ natural death |
| **R10** | Bpos_CARTE_kill | *B_pos* | → | *∅* | *k_{kill} · (C_E / S_C) · B⁺* | Mass-action CAR-T killing of B⁺ |
| **R11** | Bpos_antigen_escape_to_Bneg | *B_pos* | → | *B_neg* | *k_l · B⁺* | Antigen-loss escape conversion B⁺ → B⁻ |
| **R12** | Bneg_growth | *B_neg* | → | *2 B_neg* | *r_{Bn} · B⁻* | Escape tumour B⁻ growth |
| **R13** | Bneg_natural_death | *B_neg* | → | *∅* | *δ_B · B⁻* | Escape tumour B⁻ natural death |
| **R14** | Bneg_bystander_kill | *B_neg* | → | *∅* | *k_b · (C_E / S_C) · B⁻* | Bystander killing of antigen-negative tumour |
| **R15** | IL6_production | *∅* | → | *IL6* | *α_{IL6} · (c̃_E / (K_{m,E} + c̃_E)) · b̃ + β_{IL6}* | IL-6 production (calibrated; MM saturated, source C_E + C_X in SBML) |
| **R16** | IL6_decay | *IL6* | → | *∅* | *k_{deg,IL6} · L₆* | IL-6 first-order degradation |
| **R17** | IL10_production | *∅* | → | *IL10* | *α_{IL10} · c̃_E · b̃ + β_{IL10}* | IL-10 production (calibrated; source C_M + C_X in SBML) |
| **R18** | IL10_decay | *IL10* | → | *∅* | *k_{deg,IL10} · L₁₀* | IL-10 first-order degradation |
| **R19** | IFNg_production | *∅* | → | *IFNg* | *α_{IFNγ} · c̃_E · b̃ + β_{IFNγ}* | IFN-γ production (calibrated; source C_E in SBML) |
| **R20** | IFNg_decay | *IFNg* | → | *∅* | *(k_{deg,base} + k_{up,IFNγ} · c̃_X) · γ* | IFN-γ degradation, enhanced by exhausted CAR-T |
| **R21** | aPD1_elimination | *aPD1* | → | *∅* | *k_{elim} · A_{PD1}* | Anti-PD-1 antibody first-order clearance |

Note: Cytokine kinetic laws for R15–R20 show the calibrated Python implementation. The SBML file uses simplified proportional production terms (*PIL6, PIL10, PINFg*) driven by normalised cell counts; the 19 calibrated parameters override these during optimisation to match the benchmark data.

### **S1.6 ODEs Derived from Reaction Stoichiometry**

The net rate of change for each species is obtained by summing stoichiometric contributions from all reactions in which that species participates (net = production − consumption):

| **ODE** | **Equation** | **Reactions contributing** |
| --- | --- | --- |
| dC_E/dt | = R1 − R2 − R3 − R4 + R5 | Expansion(R1) − Death(R2) − Differentiation(R3) − Exhaustion(R4) + Reactivation(R5) |
| dC_M/dt | = R3 − R5 − R6 | Gain from R3 − Reactivation loss(R5) − Death(R6) |
| dC_X/dt | = R4 − R7 | Exhaustion influx(R4) − Clearance(R7) |
| dB⁺/dt | = R8 − R9 − R10 − R11 | Growth(R8) − Natural death(R9) − CAR-T kill(R10) − Escape(R11) |
| dB⁻/dt | = R11 + R12 − R13 − R14 | Escape influx(R11) + Growth(R12) − Natural death(R13) − Bystander kill(R14) |
| dL₆/dt | = R15 − R16 | IL-6 production(R15) − Degradation(R16) |
| dL₁₀/dt | = R17 − R18 | IL-10 production(R17) − Degradation(R18) |
| dγ/dt | = R19 − R20 | IFN-γ production(R19) − Degradation(R20) [C_X-enhanced] |
| dA/dt | = −R21 | aPD1 elimination only; no source after dosing events |

### **S1.7 CAR-T Dosing Events**

CAR-T cells are administered as a split infusion over three days. The SBML *listOfEvents* defines three discrete injections that add cells to *CARTE*:

| **Event ID** | **Trigger (time ≥)** | **C_E increment** | **Fraction of D_total** |
| --- | --- | --- | --- |
| dose_d0 | 0 days | 0.10 × D_total | 10% |
| dose_d1 | 1 day | 0.30 × D_total | 30% |
| dose_d2 | 2 days | 0.60 × D_total | 60% |

Total dose: *D_total = 1.10 × 10⁹ cells* (SBML parameter *dose_total*). Calibration ICs taken from digitised benchmark data.

### **S1.8 Calibrated Parameters (19-Parameter MA v11 Best-Fit)**

The 19 parameters below were estimated by multi-phase L-BFGS-B optimisation against digitised benchmark data (Kimmel et al., 2021). They override SBML-default cytokine production constants (R15–R20) and provide additional mechanistic detail (exhaustion gating, IL-6 MM saturation, IFN-γ exhaustion-dependent degradation). Analytical pre-conditioning derived *k_{exh} ≈ 0.044 d⁻¹* and *K_{DX} ≈ 0.25 d⁻¹* from the benchmark CART_EX decay kinetics.

| **#** | **Symbol** | **Description** | **Lower** | **Upper** | **Best-fit (MA v11)** |
| --- | --- | --- | --- | --- | --- |
| 1 | k_{prolif} | CAR-T mass-action proliferation rate (R1) | 0.10 | 30.0 | 1.8281 d⁻¹ |
| 2 | k_{exh} | Exhaustion rate (R4; PD-1 gated) | 0.015 | 0.055 | 0.04025 d⁻¹ |
| 3 | k_{kill} | Mass-action killing rate on B⁺ (R10) | 0.01 | 10.0 | 0.01290 d⁻¹ cell⁻¹ |
| 4 | k_{loss} | Antigen-escape rate B⁺→B⁻ (R11) | 1×10⁻⁴ | 0.01 | 0.1417 d⁻¹ |
| 5 | k_{byst} | Bystander killing coeff. on B⁻ (R14) | 1×10⁻⁴ | 0.005 | 0.3099 d⁻¹ cell⁻¹ |
| 6 | dE_scale | Death rate scaling factor for R2 | 0.05 | 5.0 | 7.607×10⁻² |
| 7 | E_{max,PD1} | Max fractional PD-1 inhibition (assignment rule) | 0.10 | 0.99 | 0.9749 |
| 8 | K_{DX} | Exhausted CAR-T clearance rate (R7) | 0.15 | 0.50 | 0.1735 d⁻¹ |
| 9 | α_{IL6} | IL-6 CAR-T·tumour production coeff. (R15) | 0.1 | 2000 | 1530 pg mL⁻¹ d⁻¹ |
| 10 | K_{m,E} | IL-6 MM half-saturation for c̃_E (R15) | 0.5 | 200 | 65.42 (c̃_E units) |
| 11 | β_{IL6} | IL-6 basal production rate (R15) | 0 | 25 | 17.08 pg mL⁻¹ d⁻¹ |
| 12 | k_{deg,IL6} | IL-6 first-order degradation rate (R16) | 0.1 | 8.0 | 2.380 d⁻¹ |
| 13 | α_{IL10} | IL-10 bilinear production coeff. (R17) | 0.1 | 2000 | 1.896 pg mL⁻¹ d⁻¹ |
| 14 | β_{IL10} | IL-10 basal production rate (R17) | 0 | 25 | 1.552 pg mL⁻¹ d⁻¹ |
| 15 | k_{deg,IL10} | IL-10 first-order degradation rate (R18) | 0.1 | 5.0 | 0.3121 d⁻¹ |
| 16 | α_{IFNγ} | IFN-γ bilinear production coeff. (R19) | 0.1 | 2000 | 4.837 pg mL⁻¹ d⁻¹ |
| 17 | β_{IFNγ} | IFN-γ basal production rate (R19) | 0 | 25 | 2.051 pg mL⁻¹ d⁻¹ |
| 18 | k_{deg,base} | IFN-γ baseline degradation rate (R20) | 0.1 | 5.0 | 0.5100 d⁻¹ |
| 19 | k_{up,IFNγ} | IFN-γ exhaustion-driven up-regulation (R20) | 0 | 0.30 | 9.896×10⁻⁵ (c̃_X)⁻¹ d⁻¹ |

### **S1.9 Calibration Objective Function: Log-RMSE**

The objective function minimised during calibration is the mean log-transformed root mean square error (log-RMSE) across *N* = 6 observed variables and *T* = 9 benchmark time points (days 0, 1, 3, 5, 7, 10, 14, 21, 28 post-infusion):

J(θ) = 1/N · ∑_i=1_*^N^* RMSE_i_(θ)

RMSE_i_(θ) = √( 1/*T* · ∑_j=1_*^T^* [ log₁₀(ŷ_ij_(θ)) − log₁₀(y_ij_) ]² )

where: *θ* = 19 calibrated parameters; *ŷ*ᵢⱼ(θ) = model prediction for variable *i* at time *j*; *y*ᵢⱼ = benchmark value; *N* = 6 variables; *T* = 9 time points. Only observations with *y > 0* contribute. The minimisation used L-BFGS-B with box constraints initialised from 200 LHS seeds in log-parameter space.

### **S1.10 Per-Variable Log-RMSE (Mechanistic Reaction-Based Model, MA v11)**

| **i** | **Variable (y_i)** | **SBML species** | **SBML reactions** | **T** | **log-RMSE_i** |
| --- | --- | --- | --- | --- | --- |
| 1 | Total Tumour (B⁺+B⁻) | B_pos + B_neg | R8–R14 | 9 | 0.224 |
| 2 | Effector CAR-T (PB) | CARTE | R1–R5 | 9 | 0.140 |
| 3 | Exhausted CAR-T (PB) | CARTE_EX | R4, R7 | 9 | 0.085 ✓ |
| 4 | Interleukin-6 | IL6 | R15–R16 | 9 | 0.067 ✓ |
| 5 | Interleukin-10 | IL10 | R17–R18 | 9 | 0.117 |
| 6 | Interferon-γ | IFNg | R19–R20 | 9 | 0.159 |
| **Mean J(θ*) =** | **0.132** |  |  |  |  |

✓ Variables meeting the pre-specified log-RMSE < 0.10 accuracy threshold.
