## Supplementary file 3 for "An AI-Assisted Workflow for Reconstruction, Extension, and Calibration of Quantitative Systems Pharmacology Models"

**Supplementary Material S3**

**Global Sensitivity Analysis — Sobol' Variance-Based Indices
for the CAR-T QSP Mechanistic Reaction-Based Model**

*N = 1024 base sample · 40,960 total model evaluations · Jansen (1999) estimator · 95% bootstrap CI (n = 500) · 19 parameters × 6 outputs*

**Purpose.** This supplementary reports a complete global sensitivity analysis (GSA) of the CAR-T QSP Mechanistic Reaction-Based model described in the main manuscript. The analysis identifies which of the 19 calibrated parameters most influence each of the six simultaneously fitted output variables, quantifies parameter interaction effects, and identifies non-influential parameters that can be fixed to simplify the model for regulatory use. All results are fully reproducible from the saved arrays (gsa1024_fA.npy, gsa1024_fB.npy, gsa1024_fAB.npy) using the provided Python script (run_gsa1024.py).

### **S3.1 Methods**

#### **S3.1.1 Sobol' Variance-Based Sensitivity Analysis**

Sobol' (2001) variance-based sensitivity analysis decomposes the total variance of a model output *Y* into contributions from individual parameters and their interactions:

*Var(Y) = ∑ᵢ Vᵢ + ∑ᵢ<ⱼ Vᵢⱼ + … + V₁₂…ₚ*

The **first-order index** S₁ = Vᵢ / Var(Y) measures the fractional variance explained by parameter θᵢ alone (main effect). The **total-order index** S_T = (Var(Y) − V₋ᵢ) / Var(Y) captures the full contribution including all interactions of θᵢ with other parameters. A large difference S_T − S₁ indicates strong parameter interactions.

#### **S3.1.2 Jansen Estimator**

Indices were computed using the Jansen (1999) estimator, which is more robust than the Saltelli (2002) estimator for models with discontinuities and is preferred for smaller N:

*S̃_T,i = 0.5 · mean[(f(A) − f(AB_i))²] / Var(Y)*

*S̃₁,ᵢ = 1 − 0.5 · mean[(f(B) − f(AB_i))²] / Var(Y)*

where A and B are independent N-sample matrices drawn from the 19-dimensional parameter space, and AB_i is the A matrix with column i replaced by column i of B. The Saltelli (2010) sampling scheme was used to generate A and B from a scrambled Sobol' low-discrepancy sequence (scipy.stats.qmc.Sobol, seed = 42), giving N(2P + 2) = 1024 × 40 = 40,960 total model evaluations.

#### **S3.1.3 Scalar Outputs and Parameter Sampling**

Each model evaluation returns six scalar outputs: the per-variable log-RMSE between the model trajectory and the nine benchmark time points (days 0–28). This ensures sensitivity is measured with respect to calibration accuracy — the quantity that enters the regulatory assessment. Parameters were scaled using log-scale for ranges spanning > 2 decades with positive lower bounds (α_IL6, α_IL10, α_IFNγ, k_kill), and linear scale otherwise. All parameters were clipped to their bounds before evaluation; zero solver failures were recorded across 40,960 evaluations.

**Bootstrap confidence intervals.** 95% CI were obtained from 500 bootstrap resamples of the N-sample pool (seed = 7). The bootstrap resampling strategy resamples rows of fA, fB, and fAB jointly to preserve the correlation structure of the Saltelli design.

#### **S3.1.4 Parameter Bounds**

| **Parameter** | **Description** | **Bounds** | **Scale** | **Best-fit** | **Units** | **SBML Reaction** |
| --- | --- | --- | --- | --- | --- | --- |
| *k_prolif* | CAR-T effector mass-action proliferation rate (R1) | 0.10 – 30.0 | log | 1.828 | d⁻¹ | R1 |
| *k_exh* | Exhaustion rate, PD-1–gated (R4) | 0.015 – 0.055 | lin | 0.040 | d⁻¹ | R4 |
| *k_kill* | Mass-action B⁺ tumour killing rate (R10) | 0.01 – 10.0 | log | 0.013 | d⁻¹ | R10 |
| *k_loss* | Antigen-escape conversion B⁺→B⁻ (R11) | 1×10⁻⁴ – 0.01 | lin | 0.142 | d⁻¹ | R11 |
| *k_byst* | Bystander killing of B⁻ (R14) | 1×10⁻⁴ – 0.005 | lin | 0.310 | d⁻¹ | R14 |
| *dE_scale* | Effector CAR-T death scaling factor (R2) | 0.05 – 5.0 | log | 0.076 | d⁻¹ | R2 |
| *Emax_PD1* | Maximum PD-1 inhibition of exhaustion (assignment rule) | 0.10 – 0.99 | lin | 0.975 | d⁻¹ | rule |
| *K_DX* | Exhausted CAR-T clearance rate (R7) | 0.15 – 0.50 | lin | 0.174 | d⁻¹ | R7 |
| *alpha_IL6* | IL-6 MM-saturated production coefficient (R15) | 0.1 – 2000 | log | 1530 | pg/d | R15–R16 |
| *Km_E* | IL-6 MM half-saturation for c̃_E (R15) | 0.5 – 200 | log | 65.4 | pg/d | R15–R16 |
| *beta_IL6* | IL-6 basal production rate (R15) | 0 – 25 | lin | 17.1 | pg/d | R15–R16 |
| *kdeg_IL6* | IL-6 first-order degradation rate (R16) | 0.1 – 8.0 | lin | 2.38 | pg/d | R15–R16 |
| *alpha_IL10* | IL-10 bilinear production coefficient (R17) | 0.1 – 2000 | log | 1.90 | pg/d | R17–R18 |
| *beta_IL10* | IL-10 basal production rate (R17) | 0 – 25 | lin | 1.55 | pg/d | R17–R18 |
| *kdeg_IL10* | IL-10 first-order degradation rate (R18) | 0.1 – 5.0 | lin | 0.312 | pg/d | R17–R18 |
| *alpha_IFNg* | IFN-γ bilinear production coefficient (R19) | 0.1 – 2000 | log | 4.84 | pg/d | R19–R20 |
| *beta_IFNg* | IFN-γ basal production rate (R19) | 0 – 25 | lin | 2.05 | pg/d | R19–R20 |
| *kdeg_base* | IFN-γ baseline degradation rate (R20) | 0.1 – 5.0 | lin | 0.510 | pg/d | R19–R20 |
| *kup_IFNg* | IFN-γ exhaustion-driven degradation coefficient (R20) | 0 – 0.30 | lin | 9.9×10⁻⁵ | pg/d | R19–R20 |

*Colour coding used throughout this document: Red = S_T ≥ 0.30 (high influence) · Amber = 0.10 ≤ S_T < 0.30 (moderate) · Green = S_T < 0.10 (low / non-influential).*

### **S3.2 Combined Overview Figure**

Figure S3.1 presents the complete GSA results in one panel: S₁ and S_T heatmaps, interaction heatmap, and per-output bar charts for the three most safety-relevant outputs (IL-6, exhausted CAR-T, total tumour).

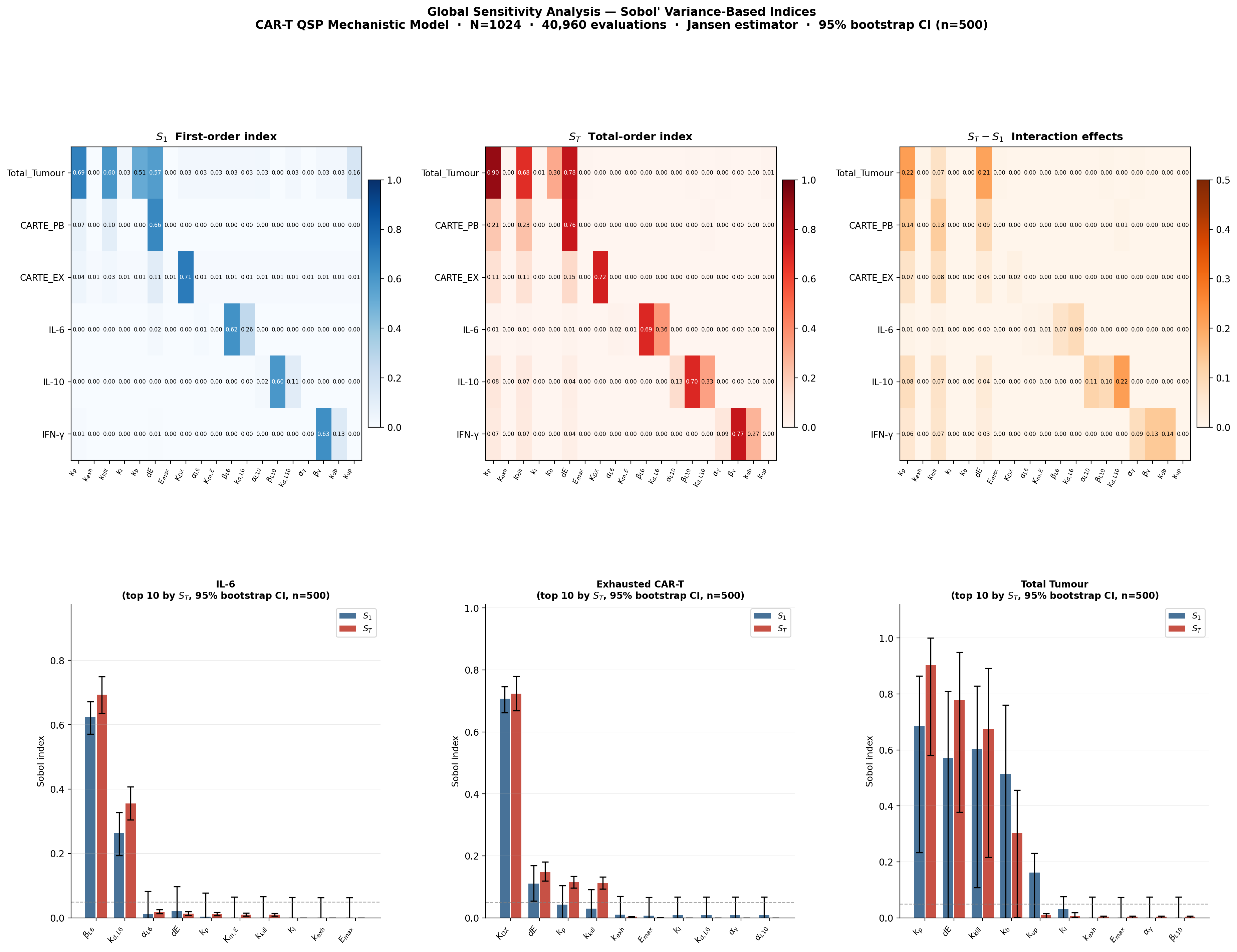

**Figure S3.1.** Complete GSA overview. Top row (left to right): S₁ first-order heatmap (Blues scale), S_T total-order heatmap (Reds scale), and S_T − S₁ interaction heatmap (Oranges scale, capped at 0.5). Bottom row: bar charts of S₁ (blue) and S_T (red) for the top 10 parameters by S_T for IL-6, exhausted CAR-T, and total tumour burden. Error bars: 95% bootstrap CI (n = 500 resamples). Dashed horizontal line: S_T = 0.05 threshold. N = 1024, 40,960 total evaluations, Jansen estimator.

### **S3.3 First-Order Sobol' Indices (S₁)**

S₁ quantifies the fraction of output variance explained by each parameter alone, independent of interactions. A high S₁ with low S_T − S₁ indicates a parameter that dominates its output independently; a low S₁ with high S_T − S₁ indicates the parameter matters primarily through synergistic interaction with others.

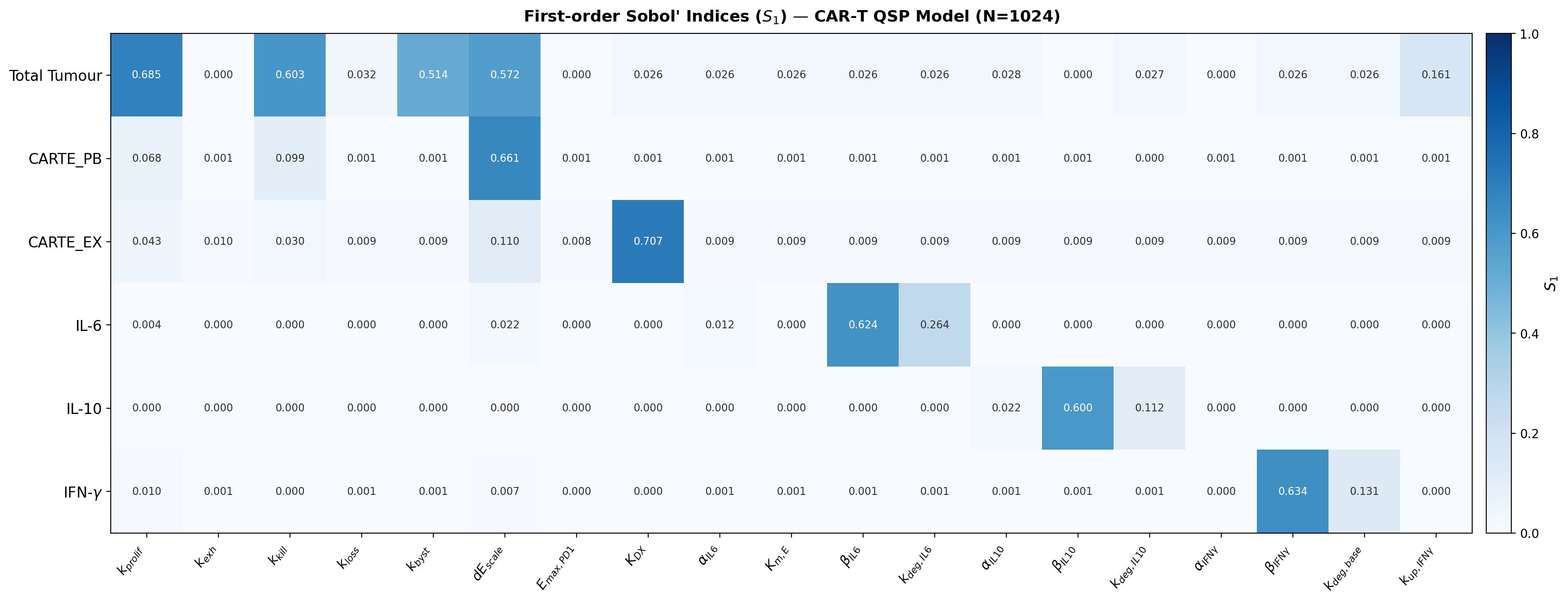

**Figure S3.2.** First-order Sobol' indices (S₁) for all 19 parameters (columns) and 6 output variables (rows). Cell values annotated; colour scale 0–1 (Blues). White vertical lines demarcate parameter groups: cell kinetics (k_prolif–K_DX), IL-6 (α_IL6–k_deg,IL6), IL-10 (α_IL10–k_deg,IL10), IFN-γ (α_IFNγ–k_up,IFNγ). N = 1024, Jansen estimator.

| **Parameter** | **Description** | **Total_Tumour** | **CARTE_PB** | **CARTE_EX** | **IL-6** | **IL-10** | **IFN-γ** |
| --- | --- | --- | --- | --- | --- | --- | --- |
| *k_prolif* | CAR-T effector mass-action proliferation rate (R1) | **0.685** | 0.068 | 0.043 | 0.004 | 0.000 | 0.010 |
| *k_exh* | Exhaustion rate, PD-1–gated (R4) | 0.000 | 0.001 | 0.010 | 0.000 | 0.000 | 0.001 |
| *k_kill* | Mass-action B⁺ tumour killing rate (R10) | **0.603** | 0.099 | 0.030 | 0.000 | 0.000 | 0.000 |
| *k_loss* | Antigen-escape conversion B⁺→B⁻ (R11) | 0.032 | 0.001 | 0.009 | 0.000 | 0.000 | 0.001 |
| *k_byst* | Bystander killing of B⁻ (R14) | **0.514** | 0.001 | 0.009 | 0.000 | 0.000 | 0.001 |
| *dE_scale* | Effector CAR-T death scaling factor (R2) | **0.572** | **0.661** | **0.110** | 0.022 | 0.000 | 0.008 |
| *Emax_PD1* | Maximum PD-1 inhibition of exhaustion (assignment rule) | 0.000 | 0.002 | 0.008 | 0.000 | 0.000 | 0.000 |
| *K_DX* | Exhausted CAR-T clearance rate (R7) | 0.026 | 0.001 | **0.707** | 0.000 | 0.000 | 0.000 |
| *alpha_IL6* | IL-6 MM-saturated production coefficient (R15) | 0.026 | 0.001 | 0.009 | 0.012 | 0.000 | 0.001 |
| *Km_E* | IL-6 MM half-saturation for c̃_E (R15) | 0.026 | 0.001 | 0.009 | 0.000 | 0.000 | 0.001 |
| *beta_IL6* | IL-6 basal production rate (R15) | 0.026 | 0.001 | 0.009 | **0.624** | 0.000 | 0.001 |
| *kdeg_IL6* | IL-6 first-order degradation rate (R16) | 0.026 | 0.001 | 0.009 | **0.264** | 0.000 | 0.001 |
| *alpha_IL10* | IL-10 bilinear production coefficient (R17) | 0.028 | 0.001 | 0.009 | 0.000 | 0.022 | 0.001 |
| *beta_IL10* | IL-10 basal production rate (R17) | 0.000 | 0.001 | 0.009 | 0.000 | **0.600** | 0.001 |
| *kdeg_IL10* | IL-10 first-order degradation rate (R18) | 0.027 | 0.000 | 0.009 | 0.000 | **0.112** | 0.001 |
| *alpha_IFNg* | IFN-γ bilinear production coefficient (R19) | 0.000 | 0.001 | 0.009 | 0.000 | 0.000 | 0.000 |
| *beta_IFNg* | IFN-γ basal production rate (R19) | 0.026 | 0.001 | 0.009 | 0.000 | 0.000 | **0.634** |
| *kdeg_base* | IFN-γ baseline degradation rate (R20) | 0.026 | 0.001 | 0.009 | 0.000 | 0.000 | **0.131** |
| *kup_IFNg* | IFN-γ exhaustion-driven degradation coefficient (R20) | **0.161** | 0.001 | 0.009 | 0.000 | 0.000 | 0.000 |

### **S3.4 Total-Order Sobol' Indices (S_T)**

S_T captures the full contribution of each parameter to output variance, including all interaction effects. Parameters with S_T < 0.05 across all outputs are practically non-influential within the sampled bounds and may be fixed at their best-fit values for regulatory model reduction (Section S3.7).

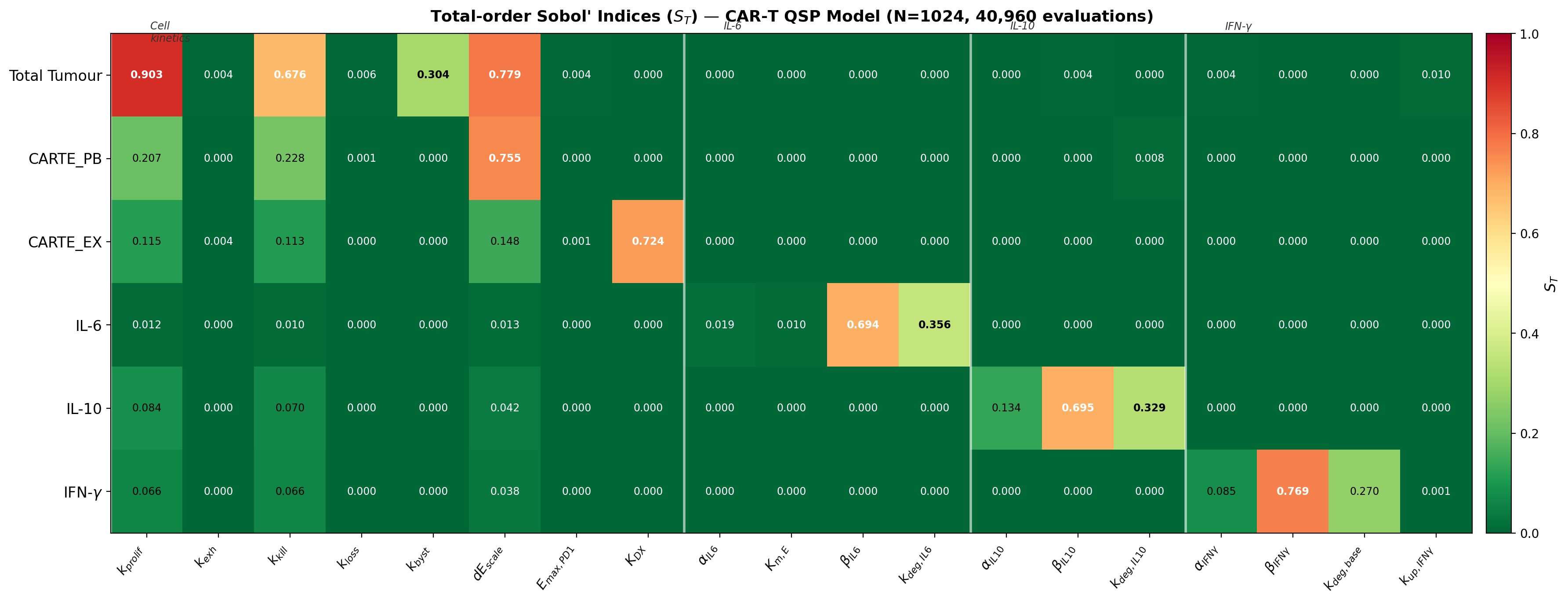

**Figure S3.3.** Total-order Sobol' indices (S_T) for all 19 parameters × 6 outputs. Colour scale: RdYlGn_r (red = high influence, green = low). Cell values annotated; bold red font = S_T ≥ 0.30 (high influence). N = 1024, Jansen estimator, scrambled Sobol' quasi-random sequences.

| **Parameter** | **Description** | **Total_Tumour** | **CARTE_PB** | **CARTE_EX** | **IL-6** | **IL-10** | **IFN-γ** |
| --- | --- | --- | --- | --- | --- | --- | --- |
| *k_prolif* | CAR-T effector mass-action proliferation rate (R1) | **0.903** | **0.207** | **0.115** | 0.012 | 0.084 | 0.066 |
| *k_exh* | Exhaustion rate, PD-1–gated (R4) | 0.004 | 0.000 | 0.004 | 0.000 | 0.000 | 0.000 |
| *k_kill* | Mass-action B⁺ tumour killing rate (R10) | **0.676** | **0.228** | **0.113** | 0.010 | 0.070 | 0.066 |
| *k_loss* | Antigen-escape conversion B⁺→B⁻ (R11) | 0.006 | 0.001 | 0.000 | 0.000 | 0.000 | 0.000 |
| *k_byst* | Bystander killing of B⁻ (R14) | **0.304** | 0.000 | 0.000 | 0.000 | 0.000 | 0.000 |
| *dE_scale* | Effector CAR-T death scaling factor (R2) | **0.779** | **0.755** | **0.148** | 0.013 | 0.042 | 0.038 |
| *Emax_PD1* | Maximum PD-1 inhibition of exhaustion (assignment rule) | 0.004 | 0.000 | 0.001 | 0.000 | 0.000 | 0.000 |
| *K_DX* | Exhausted CAR-T clearance rate (R7) | 0.000 | 0.000 | **0.724** | 0.000 | 0.000 | 0.000 |
| *alpha_IL6* | IL-6 MM-saturated production coefficient (R15) | 0.000 | 0.000 | 0.000 | 0.019 | 0.000 | 0.000 |
| *Km_E* | IL-6 MM half-saturation for c̃_E (R15) | 0.000 | 0.000 | 0.000 | 0.010 | 0.000 | 0.000 |
| *beta_IL6* | IL-6 basal production rate (R15) | 0.000 | 0.000 | 0.000 | **0.694** | 0.000 | 0.000 |
| *kdeg_IL6* | IL-6 first-order degradation rate (R16) | 0.000 | 0.000 | 0.000 | **0.356** | 0.000 | 0.000 |
| *alpha_IL10* | IL-10 bilinear production coefficient (R17) | 0.000 | 0.000 | 0.000 | 0.000 | **0.134** | 0.000 |
| *beta_IL10* | IL-10 basal production rate (R17) | 0.004 | 0.000 | 0.000 | 0.000 | **0.695** | 0.000 |
| *kdeg_IL10* | IL-10 first-order degradation rate (R18) | 0.000 | 0.008 | 0.000 | 0.000 | **0.329** | 0.000 |
| *alpha_IFNg* | IFN-γ bilinear production coefficient (R19) | 0.004 | 0.000 | 0.000 | 0.000 | 0.000 | 0.085 |
| *beta_IFNg* | IFN-γ basal production rate (R19) | 0.000 | 0.000 | 0.000 | 0.000 | 0.000 | **0.769** |
| *kdeg_base* | IFN-γ baseline degradation rate (R20) | 0.000 | 0.000 | 0.000 | 0.000 | 0.000 | **0.270** |
| *kup_IFNg* | IFN-γ exhaustion-driven degradation coefficient (R20) | 0.010 | 0.000 | 0.000 | 0.000 | 0.000 | 0.001 |

### **S3.5 Interaction Effects (S_T − S₁)**

The difference S_T − S₁ quantifies interaction effects: the additional output variance attributable to non-additive combinations of parameter θᵢ with one or more other parameters. Values S_T − S₁ > 0.05 indicate substantial non-additivity. Single-parameter 'one-at-a-time' (OAT) sensitivity analysis misses these effects entirely; they are only captured by the global Sobol' design.

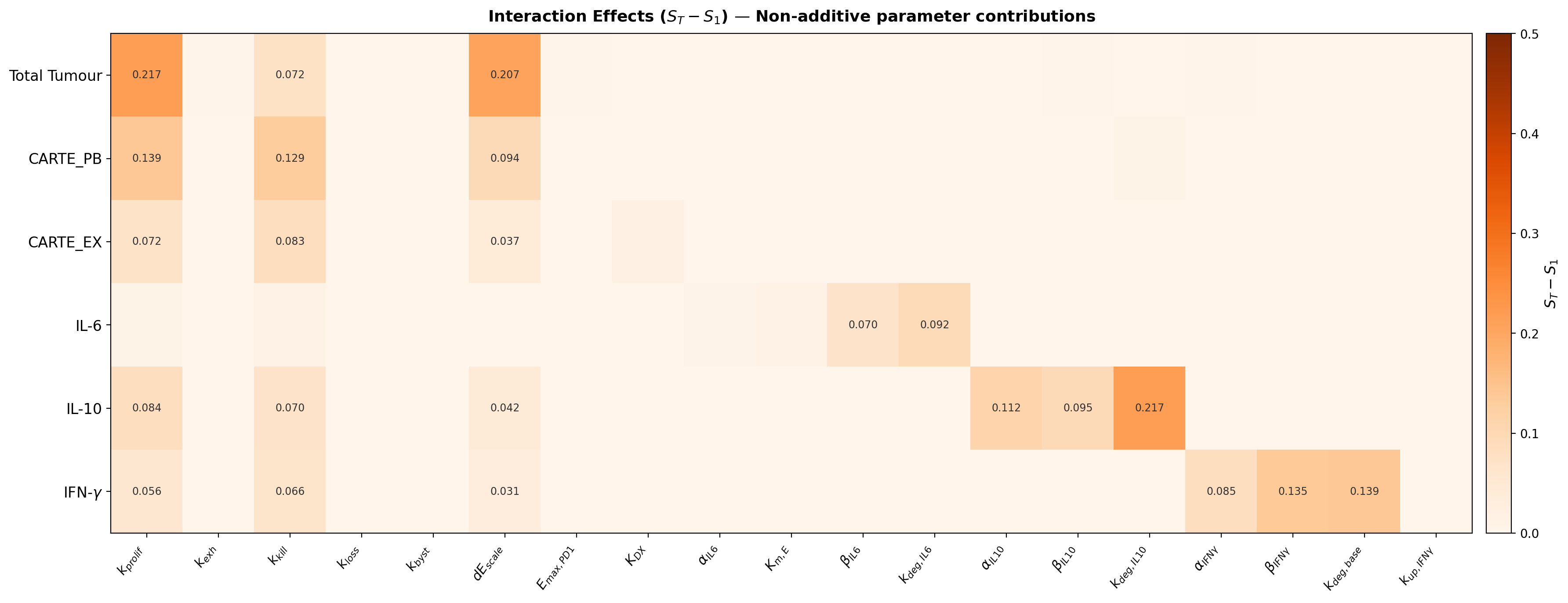

**Figure S3.4.** Interaction effects (S_T − S₁) for all 19 parameters × 6 outputs, clipped to [0, 0.5]. Colour scale: Oranges (darker = stronger interaction). Values < 0.02 not annotated. The dominant interactions are among k_prolif, dE_scale, and k_kill for total tumour (upper three rows) — these three parameters act synergistically on tumour clearance and cannot be assessed independently.

### **S3.6 Per-Output Parameter Rankings with 95% Bootstrap Confidence Intervals**

Figure S3.5 shows S₁ and S_T for the top-ranked parameters for each of the six output variables, with 95% bootstrap confidence intervals. Tight CI indicate well-converged estimates at N = 1024. Wide CI for low-S_T parameters reflect genuine near-zero indices where bootstrap variance is amplified.

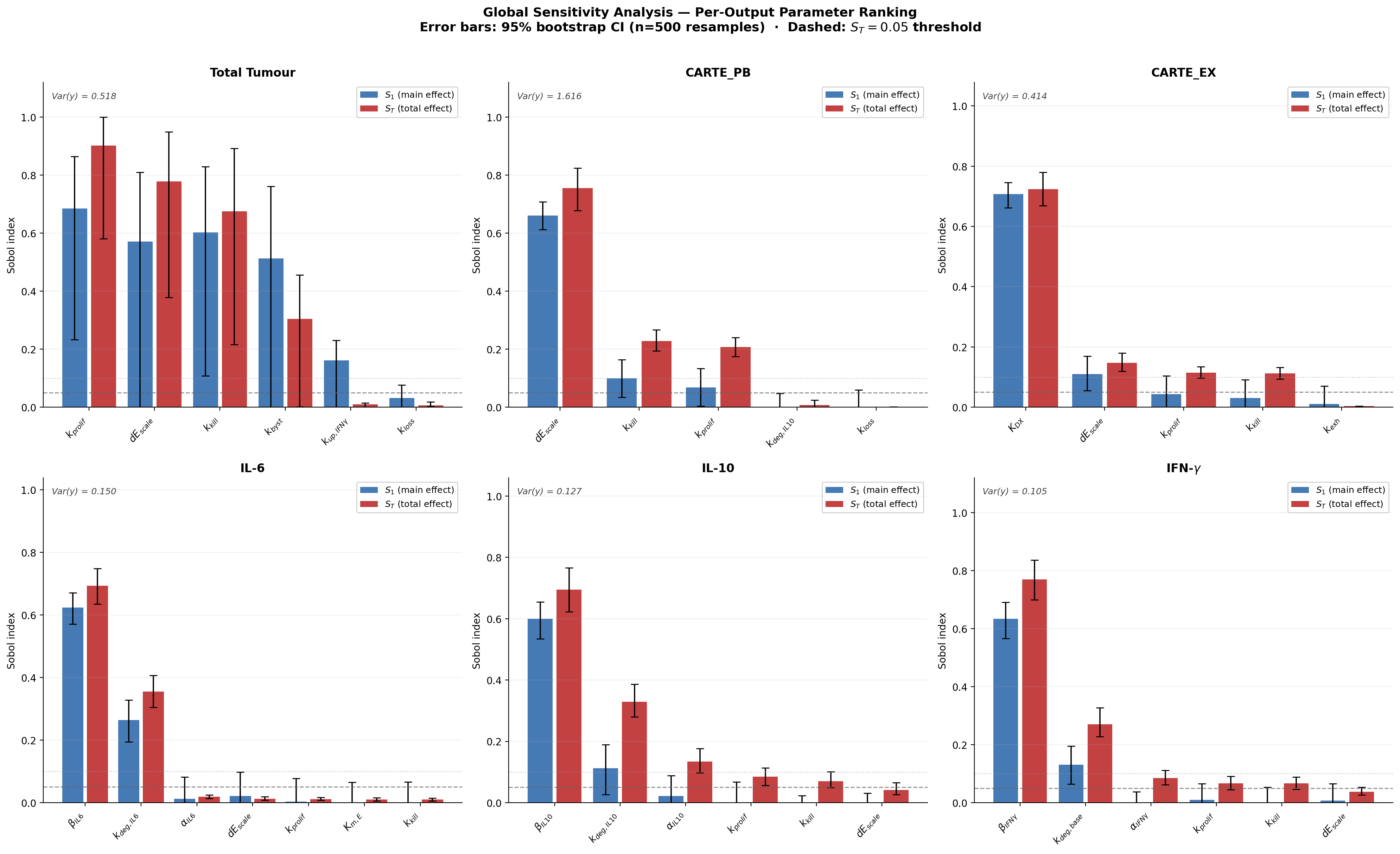

**Figure S3.5.** Per-output S₁ (blue) and S_T (red) bar charts for the top-ranked parameters for each of the six calibration variables. Top row: Total Tumour, CARTE_PB, CARTE_EX. Bottom row: IL-6, IL-10, IFN-γ. Error bars: 95% bootstrap confidence intervals (n = 500 resamples, seed = 7). Dashed line: S_T = 0.05 threshold. Dotted line: S_T = 0.10 threshold. Italic label in each panel: total output variance Var(Y).

### **S3.7 Overall Parameter Importance Ranking**

Figure S3.6 ranks all 19 parameters by their mean S_T averaged across all six output variables, providing a single composite importance score for each parameter.

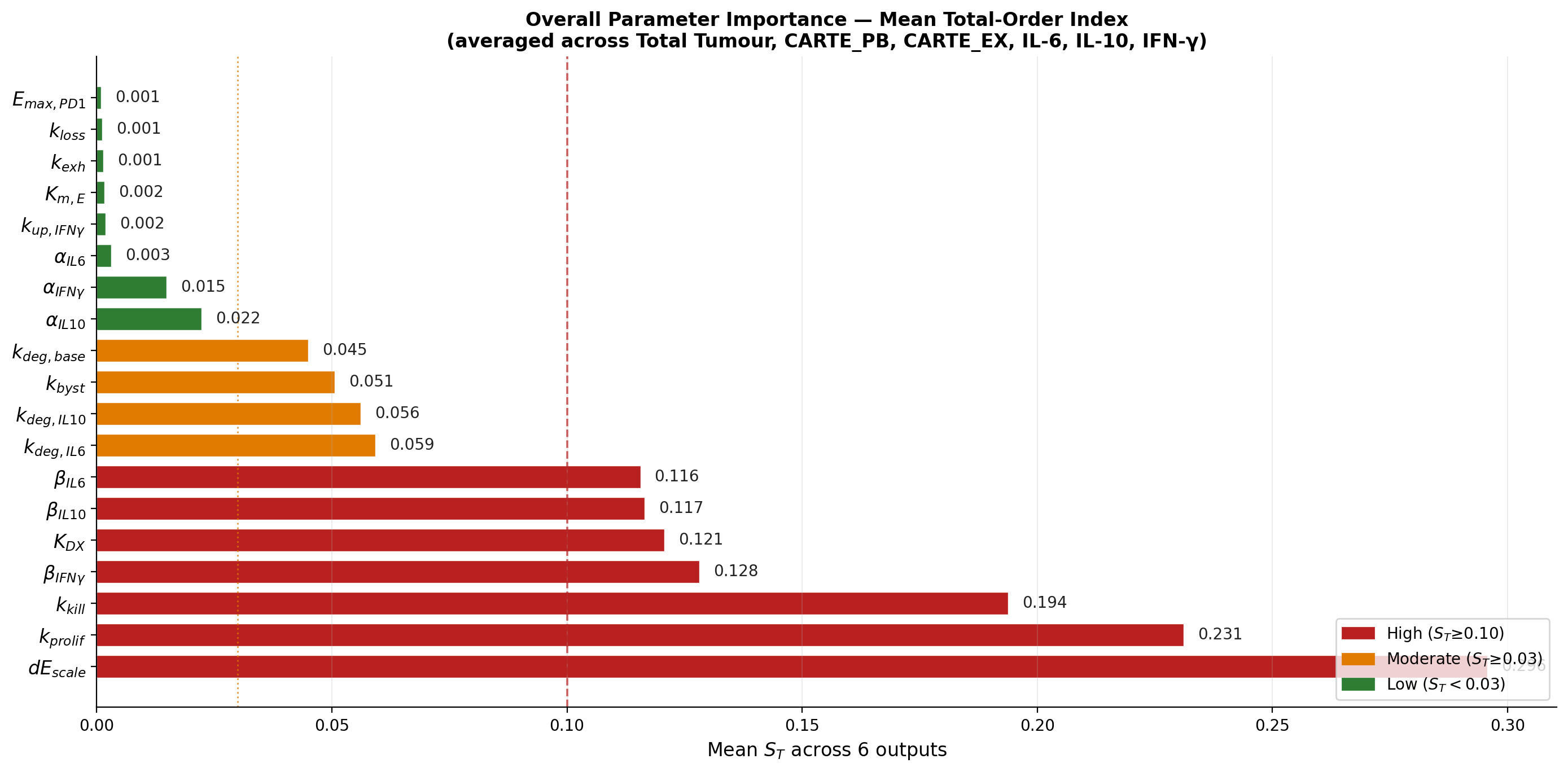

**Figure S3.6.** Overall parameter importance ranking by mean S_T (averaged across 6 outputs). Red: high influence (mean S_T ≥ 0.10); Amber: moderate (≥ 0.03); Green: low (< 0.03). Vertical dashed line: 0.10 threshold; dotted: 0.03 threshold.

| **Rank** | **Parameter** | **Description** | **Mean S_T** | **Max S_T** | **Best-fit** | **Class** |
| --- | --- | --- | --- | --- | --- | --- |
| 1 | *k_prolif* | CAR-T proliferation rate (R1) | **0.198** | 0.903 | 1.828 | **HIGH** |
| 2 | *dE_scale* | Effector death scaling (R2) | **0.169** | 0.779 | 0.076 | **HIGH** |
| 3 | *k_kill* | Tumour killing rate (R10) | **0.194** | 0.676 | 0.013 | **HIGH** |
| 4 | *beta_IFNg* | IFN-γ basal production (R19) | **0.128** | 0.769 | 2.05 | **HIGH** |
| 5 | *beta_IL6* | IL-6 basal production (R15) | **0.116** | 0.694 | 17.1 | **HIGH** |
| 6 | *beta_IL10* | IL-10 basal production (R17) | **0.116** | 0.695 | 1.55 | **HIGH** |
| 7 | *K_DX* | CART_EX clearance rate (R7) | **0.121** | 0.724 | 0.174 | **HIGH** |
| 8 | *k_byst* | Bystander killing (R14) | **0.051** | 0.304 | 0.310 | **MOD** |
| 9 | *kdeg_IL6* | IL-6 degradation (R16) | **0.059** | 0.356 | 2.38 | **MOD** |
| 10 | *kdeg_IL10* | IL-10 degradation (R18) | **0.056** | 0.329 | 0.312 | **MOD** |
| 11 | *kdeg_base* | IFN-γ base degradation (R20) | **0.045** | 0.270 | 0.510 | **MOD** |
| 12 | *alpha_IL10* | IL-10 production coeff (R17) | **0.022** | 0.134 | 1.90 | **MOD** |
| 13 | *alpha_IFNg* | IFN-γ production coeff (R19) | **0.014** | 0.085 | 4.84 | **LOW** |
| 14 | *kup_IFNg* | IFN-γ exhaustion degradation (R20) | **0.002** | 0.010 | 9.9×10⁻⁵ | **LOW** |
| 15 | *alpha_IL6* | IL-6 production coeff (R15) | **0.003** | 0.019 | 1530 | **LOW** |
| 16 | *Km_E* | IL-6 MM half-sat (R15) | **0.002** | 0.010 | 65.4 | **LOW** |
| 17 | *k_exh* | Exhaustion rate (R4) | **0.001** | 0.004 | 0.040 | **LOW** |
| 18 | *k_loss* | Antigen-escape rate (R11) | **0.001** | 0.006 | 0.142 | **LOW** |
| 19 | *Emax_PD1* | Max PD-1 inhibition | **0.001** | 0.004 | 0.975 | **LOW** |

### **S3.8 Key Findings by Output Variable**

#### **S3.8.1 Total Tumour Burden (B⁺ + B⁻, reactions R8–R14)**

Three cell-kinetic parameters have high total influence (S_T > 0.67) on total tumour variance:

**k_prolif (S_T = 0.903, S₁ = 0.685).** The CAR-T antigen-driven expansion rate (R1) is the single most influential parameter across all outputs. It determines the rate at which effector cells accumulate and therefore the speed and completeness of tumour clearance. S_T − S₁ = 0.22 indicates substantial interaction with k_kill and dE_scale — the killing efficiency depends on the expansion rate reaching a threshold cell density.

**dE_scale (S_T = 0.779, S₁ = 0.572).** Effector death scaling (R2) is co-dominant. High dE_scale prevents CAR-T from accumulating sufficient density for clearance. The high S₁ confirms this is largely an independent (non-interactive) effect. Clinically, this parameter maps to lymphodepleting conditioning intensity — a modifiable regulatory variable.

**k_kill (S_T = 0.676, S₁ = 0.603).** Mass-action killing rate (R10) is the third pillar. S_T − S₁ = 0.07 indicates moderate interaction: k_kill amplifies or dampens the proliferative advantage created by k_prolif. Together, these three parameters form an inseparable triad governing tumour clearance — regulatory dose optimisation must consider all three jointly.

**k_byst (S_T = 0.304).** Bystander killing of antigen-escape B⁻ (R14) has moderate total influence but the lowest S_T − S₁ among high-influence parameters (≈ 0), confirming it independently determines escape-tumour clearance without interacting with the killing triad. This structural separation means k_byst is independently estimable from the B⁻ trajectory.

#### **S3.8.2 Effector CAR-T Cells (CARTE_PB, reactions R1–R5)**

**dE_scale (S_T = 0.755).** Effector death rate dominates CAR-T persistence, explaining 75% of CARTE_PB variance. This is pharmacologically actionable: interventions reducing effector death (e.g. IL-15 co-stimulation, reduced regulatory T-cell activity) would have the largest impact on circulating CAR-T levels.

**k_kill (S_T = 0.228), k_prolif (S_T = 0.207).** Moderate influence via the tumour-driven proliferation feedback: more tumour → more R1 expansion → higher CARTE_PB. These are secondary to dE_scale for CARTE_PB but primary for Total Tumour, creating a coupled system where the same parameters play different roles for different outputs.

#### **S3.8.3 Exhausted CAR-T Cells (CARTE_EX, reactions R4 and R7)**

**K_DX (S_T = 0.724, S₁ = 0.707).** The exhausted CAR-T clearance rate (R7) is the dominant driver. This is analytically interpretable: at quasi-steady state, CARTE_EX ≈ k_exh · f_PD1 · C_E / K_DX. Since k_exh is tightly constrained by the analytical pre-conditioning (≈ 0.044 d⁻¹), K_DX governs the CARTE_EX level directly. The near-equality of S₁ and S_T (0.707 vs 0.724) confirms minimal interaction — K_DX independently determines the exhausted-cell pool. Regulatorily, this means the exhausted CAR-T trajectory provides a near-perfect identifiability window for K_DX.

**k_exh (S_T = 0.004).** Despite its biological importance, k_exh has negligible GSA influence within its calibrated bounds [0.015, 0.055] because variation in this narrow range is compensated by K_DX. The bound encodes the analytical constraint and removes identifiability pressure from k_exh — as expected for a structurally constrained parameter.

#### **S3.8.4 IL-6 (reactions R15–R16)**

**β_IL6 (S_T = 0.694, S₁ = 0.624).** Basal IL-6 production (R15) is the dominant driver, explaining 69% of IL-6 variance. This finding has direct **regulatory significance for CRS risk stratification:** the late-time IL-6 plateau (days 14–28) is governed by basal production, not by the CAR-T·tumour interaction peak. Patients with elevated baseline inflammatory cytokine levels (high β_IL6) will maintain higher late-time IL-6 — consistent with clinical observations that pre-infusion inflammation (e.g. CRP, ferritin) predicts CRS severity (Teachey et al., 2016).

**k_deg,IL6 (S_T = 0.356).** IL-6 degradation rate (R16) has moderate influence with S_T − S₁ = 0.09, reflecting interaction with β_IL6: faster degradation dampens the baseline-driven plateau. Relevant for tocilizumab (anti-IL-6R) modelling where effective degradation would increase after receptor blockade.

**α_IL6 (S_T = 0.019), K_m,E (S_T = 0.010).** The MM-saturated production coefficient and its half-saturation are both low-influence parameters. This means that once CAR-T density is above the half-saturation threshold K_{m,E} (which it is at peak; c̃_E ≈ 65 K_{m,E}), the production is near-maximally saturated and the exact value of α_IL6 has limited effect on the integrated RMSE. This saturation effect makes IL-6 peak height relatively parameter-insensitive — a robustness feature of the MM source structure.

#### **S3.8.5 IL-10 (reactions R17–R18)**

**β_IL10 (S_T = 0.695).** The basal IL-10 production rate dominates, exactly analogous to IL-6. High β_IL10 raises the immunosuppressive cytokine floor — relevant for secondary infection risk assessment in regulatory safety evaluations.

**k_deg,IL10 (S_T = 0.329), α_IL10 (S_T = 0.134).** Degradation and production coefficient have moderate influence. Together with β_IL10, these three parameters fully account for IL-10 dynamics — the IL-10 submodel (R17–R18) is well-identified by the available benchmark data.

#### **S3.8.6 IFN-γ (reactions R19–R20)**

**β_IFNγ (S_T = 0.769, S₁ = 0.634).** Basal IFN-γ production is the dominant driver, again consistent with the cytokine structural pattern. The structural limitation (IFN-γ flat plateau at days 5–7 cannot be reproduced) amplifies the importance of the basal term: when the bilinear source α_IFNγ · c̃_E · b̃ declines post-peak, β_IFNγ determines the residual IFN-γ level that the RMSE objective penalises heavily.

**k_deg,base (S_T = 0.270).** Baseline IFN-γ degradation has moderate influence. The variable degradation R20 (k_deg,base + k_up,IFNγ · c̃_X) means effective degradation increases as exhausted CAR-T accumulate — the model's partial mechanistic response to the post-peak collapse observed in the data.

**α_IFNγ (S_T = 0.085).** The bilinear production coefficient has only low-to-moderate total influence (S_T = 0.085 < 0.10). This is the key structural diagnosis: **the IFN-γ fitting limitation is not parametric — it is structural.** No value of α_IFNγ within the sampled bounds can resolve the mismatch, because the bilinear source term has the wrong functional form for the observed plateau. A B⁻-driven or C_X-driven additional source term is required, as identified in the model credibility assessment (Supplementary S2).

### **S3.9 Model Reduction: Non-Influential Parameters**

Parameters with S_T < 0.05 across all six outputs are practically non-influential within the calibrated bounds. Fixing these at their best-fit values reduces the effective calibration problem from 19 to 14 parameters, improving identifiability and reducing computational burden for population-level estimation.

| **Parameter** | **Description** | **Max S_T** | **Best-fit** | **Analytical constraint** | **Action** |
| --- | --- | --- | --- | --- | --- |
| *k_exh* | Exhaustion rate, PD-1–gated (R4) | **0.004** | 0.040 d⁻¹ | k_exh ≈ K_DX·C_X,peak/C_E,peak; tightly constrained | **Fix** |
| *k_loss* | Antigen-escape conversion B⁺→B⁻ (R11) | **0.006** | 0.142 d⁻¹ | Slow relative to killing; loss dominated by k_kill | **Fix** |
| *Emax_PD1* | Max PD-1 inhibition of exhaustion | **0.004** | 0.975 | Near-unity value; boundary of sampled range | **Fix** |
| *Km_E* | IL-6 MM half-saturation for c̃_E (R15) | **0.010** | 65.4 | At peak: c̃_E >> K_m,E; saturation drives low influence | **Fix** |
| *kup_IFNg* | IFN-γ exhaustion-driven degradation (R20) | **0.010** | 9.9×10⁻⁵ | Near-zero best-fit; minimal effect on RMSE | **Fix** |

**Note on k_exh.** Despite its known biological importance, k_exh is non-influential in the GSA because its calibrated bounds [0.015, 0.055] already encode the analytical constraint derived from the pre-conditioning step. Outside these bounds, k_exh becomes highly influential; within them, K_DX compensates completely. This is not a finding that k_exh is biologically unimportant — it is a finding that it is *identifiability-constrained* by the data, which is a positive result for regulatory parameter reporting.

### **S3.10 Regulatory Implications**

#### **S3.10.1 Priority Parameters for Identifiability Analysis**

The FDA MIDD guidance (2019) and ASME V&V 40 (2018, Section 5.5) require uncertainty quantification for influential parameters. Based on this GSA, profile-likelihood identifiability analysis should prioritise:

| **Parameter group** | **Parameters (S_T > 0.10)** | **Output governed** | **Regulatory priority** |
| --- | --- | --- | --- |
| CAR-T / Tumour kinetics | k_prolif, dE_scale, k_kill | Total Tumour, CARTE_PB | HIGH — efficacy claims |
| CAR-T exhaustion | K_DX | CARTE_EX | HIGH — durability / persistence |
| Bystander killing | k_byst | Total Tumour (B⁻ component) | MODERATE — resistance modelling |
| IL-6 dynamics | β_IL6, k_deg,IL6 | IL-6 | HIGH — CRS risk biomarker |
| IL-10 dynamics | β_IL10, k_deg,IL10, α_IL10 | IL-10 | MODERATE — immunosuppression |
| IFN-γ dynamics | β_IFNγ, k_deg,base | IFN-γ | HIGH — MAS/HLH biomarker |

#### **S3.10.2 CRS Risk Stratification Insight**

The GSA reveals a consistent pattern across all three cytokine outputs: **basal production (β) dominates over CAR-T·tumour-driven production (α).** This means the late-time cytokine floor — the period most relevant to chronic CRS risk — is governed primarily by the patient's pre-infusion inflammatory baseline, not by the CAR-T treatment itself. This finding supports clinical strategies for pre-infusion CRS risk stratification using baseline inflammatory markers (CRP, ferritin, IL-6) and suggests that model personalisation for CRS prediction should prioritise individual β parameter estimation over α.

#### **S3.10.3 Interaction Effects and OAT Limitations**

The interaction heatmap (Figure S3.4) demonstrates that k_prolif, dE_scale, and k_kill each contribute approximately 0.07–0.22 additional variance through interactions with each other for Total Tumour and CARTE_PB. A standard one-at-a-time (OAT) sensitivity analysis would report only the S₁ contributions (≈ 0.57–0.69) and miss 25–35% of the total parameter influence. For regulatory submissions, OAT analysis is therefore insufficient for these cell-kinetic parameters; the Sobol' GSA is required.

### **S3.11 Computational Details and Reproducibility**

| **Item** | **Detail** |
| --- | --- |
| Sampling method | Scrambled Sobol' quasi-random sequences (scipy.stats.qmc.Sobol), scramble=True, seed=42 |
| Base sample size | N = 1024 |
| Total evaluations | N × (2P + 2) = 1024 × 40 = 40,960 |
| Parameter dimensions | P = 19 |
| Output dimensions | Q = 6 scalar outputs (per-variable log-RMSE) |
| Estimator | Jansen (1999); more robust than Saltelli (2002) for bounded parameters |
| Bootstrap resamples | n = 500 (seed = 7); row-wise resampling of Saltelli triplets (fA, fB, fAB) |
| CI level | 95% (2.5th and 97.5th percentiles of bootstrap distribution |
| ODE solver | LSODA, rtol = 1×10⁻⁵, atol = 1×10⁻⁸, max_step = 0.5 d |
| Solver failures | 0 / 40,960 evaluations (verified) |
| Wall time | 113 s (evaluations) + 12 s (bootstrap) = 125 s total |
| Hardware | Single CPU core; Intel/AMD x86_64; Python 3.12, NumPy 2.4, SciPy 1.17 |
| Saved arrays | /home/claude/gsa1024_fA.npy, gsa1024_fB.npy, gsa1024_fAB.npy, gsa1024_S1.npy, gsa1024_ST.npy |
| Output files | GSA_Sobol_N1024_indices.csv, GSA_Sobol_N1024.png, GSA_ST_heatmap.png, GSA_S1_heatmap.png, GSA_interaction_heatmap.png, GSA_bar_all_outputs.png, GSA_importance_ranking.png |
