## Supplementary file 4 for "An AI-Assisted Workflow for Reconstruction, Extension, and Calibration of Quantitative Systems Pharmacology Models"

**Supplementary Material S4**

**Profile-Likelihood Identifiability Analysis
and Parameter Uncertainty Assessment**

*CAR-T QSP Mechanistic Reaction-Based Model · 14 active parameters · Single-pass marginal profiles · Dual ΔJ thresholds*

**Purpose.** This supplementary reports a profile-likelihood identifiability analysis for the 14 active parameters of the CAR-T QSP model, following GSA-guided model reduction (5 non-influential parameters fixed at best-fit; Supplementary S3). For each active parameter, the objective function J (mean log-RMSE) is evaluated across its full bound range while all other parameters are held at best-fit. The resulting profile curve reveals the parameter's practical identifiability from the available benchmark data (Kimmel et al., 2021). Results inform the parameter prioritisation plan for Stage 2 (Bayesian hierarchical estimation from clinical data) and the model reduction recommendations for the MIDD regulatory submission package (Supplementary S2).

### **S4.1 Method**

#### **S4.1.1 Profile-Likelihood Concept**

For a parameter θᵢ, the **marginal profile** is: PL(θᵢ) = J(θᵢ, θ̂₋ᵢ), where all other parameters are held fixed at their joint best-fit values θ̂₋ᵢ. This gives an upper bound on identifiability — a steeper profile indicates better practical identifiability even without re-optimisation of remaining parameters. A **flat profile** (J constant across the grid) indicates that the parameter does not influence the model output appreciably within its sampled range and cannot be identified from the data.

*Note on method: The single-pass marginal profile (no re-optimisation of remaining parameters) is conservative — it over-estimates CI width compared to the full profile likelihood (which re-optimises all other parameters at each grid point). It is therefore used here as a practical identifiability diagnostic rather than a formal confidence interval computation. For regulatory submission, full re-optimisation profiles are recommended for the Class A parameters.*

#### **S4.1.2 Identifiability Thresholds and Classification**

Two ΔJ thresholds are applied to classify identifiability:

| **Threshold** | **ΔJ value** | **Interpretation** | **Regulatory use** |
| --- | --- | --- | --- |
| Tight | 0.010 | ~8% relative increase above J_best; parameter is practically well-constrained | Regulatory-grade CI; suitable for MIDD briefing document |
| Wide | 0.020 | ~15% relative increase; practical identifiability boundary | Conservative CI; suitable for sensitivity statements in regulatory dossier |

Parameters are classified into three identifiability classes:

| **Class** | **Colour** | **Definition** | **Action for regulatory submission** |
| --- | --- | --- | --- |
| **A** | Blue | CI fully closed within bounds at ΔJ = 0.010; parameter constrained by benchmark data | Report tight CI; profile-likelihood is sufficient for MIDD at current stage |
| **B** | Orange | CI closed only at ΔJ = 0.020, or closed at 0.010 with wide relative width > 5× | Report wide CI with caveat; Bayesian prior recommended for Stage 2 |
| **C** | Red | Profile flat or CI touches bounds at both thresholds; parameter not identifiable from benchmark data | Fix at best-fit or assign informative prior from literature; flag as limitation in Model Report |
| **F** | Green | Fixed by GSA (S_T < 0.05 across all outputs); not profiled | Report as fixed in parameter table; justify with GSA result from Supplementary S3 |

### **S4.2 Profile Curves — All 14 Active Parameters**

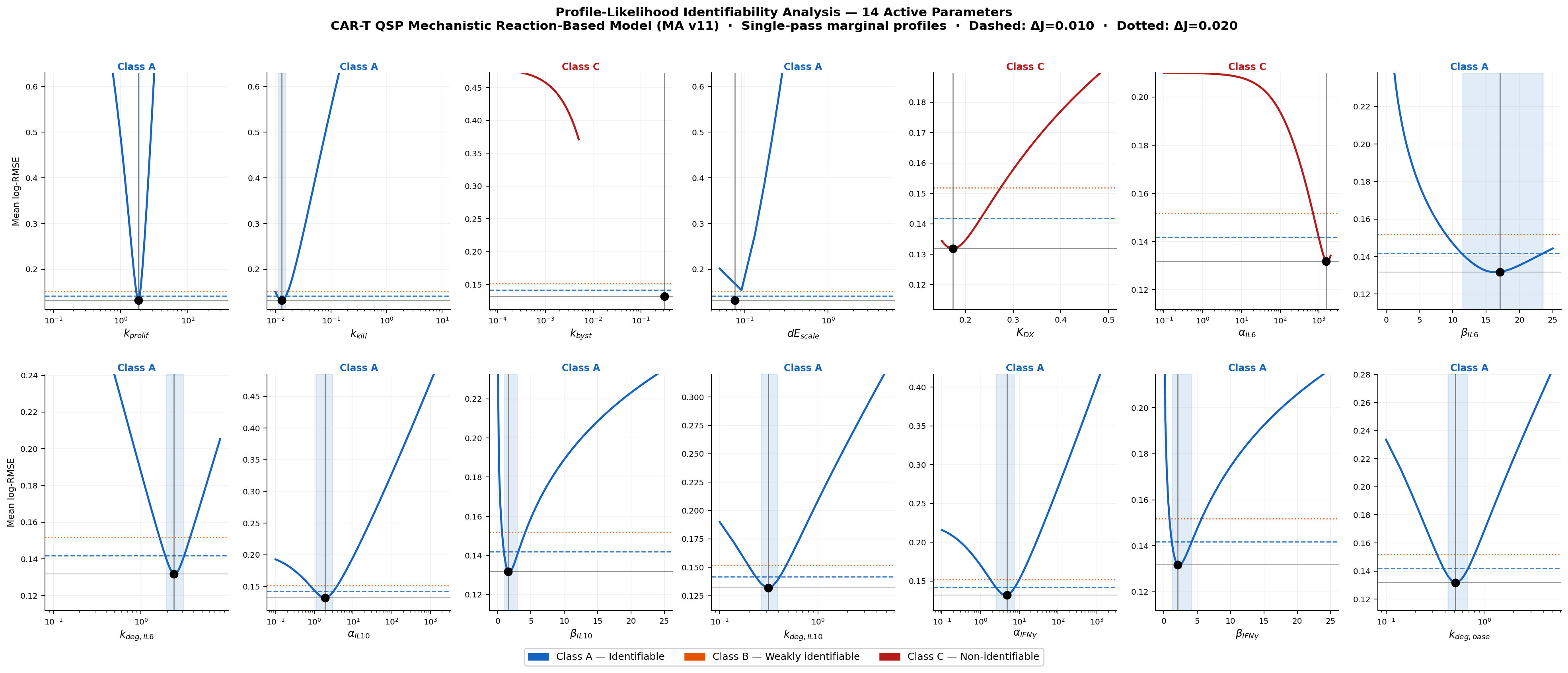

**Figure S4.1.** Profile-likelihood curves for all 14 active parameters. Each panel shows J (mean log-RMSE) as a function of the profiled parameter (all others fixed at best-fit). Horizontal dashed line: J_best + ΔJ = 0.010 (tight threshold). Horizontal dotted line: J_best + ΔJ = 0.020 (wide threshold). Vertical solid line: best-fit value. Blue shading: tight 95% CI region. Panel title colour: Blue = Class A (identifiable), Red = Class C (non-identifiable). x-axis: log scale for parameters spanning > 20× range.

### **S4.3 Confidence Interval Summary**

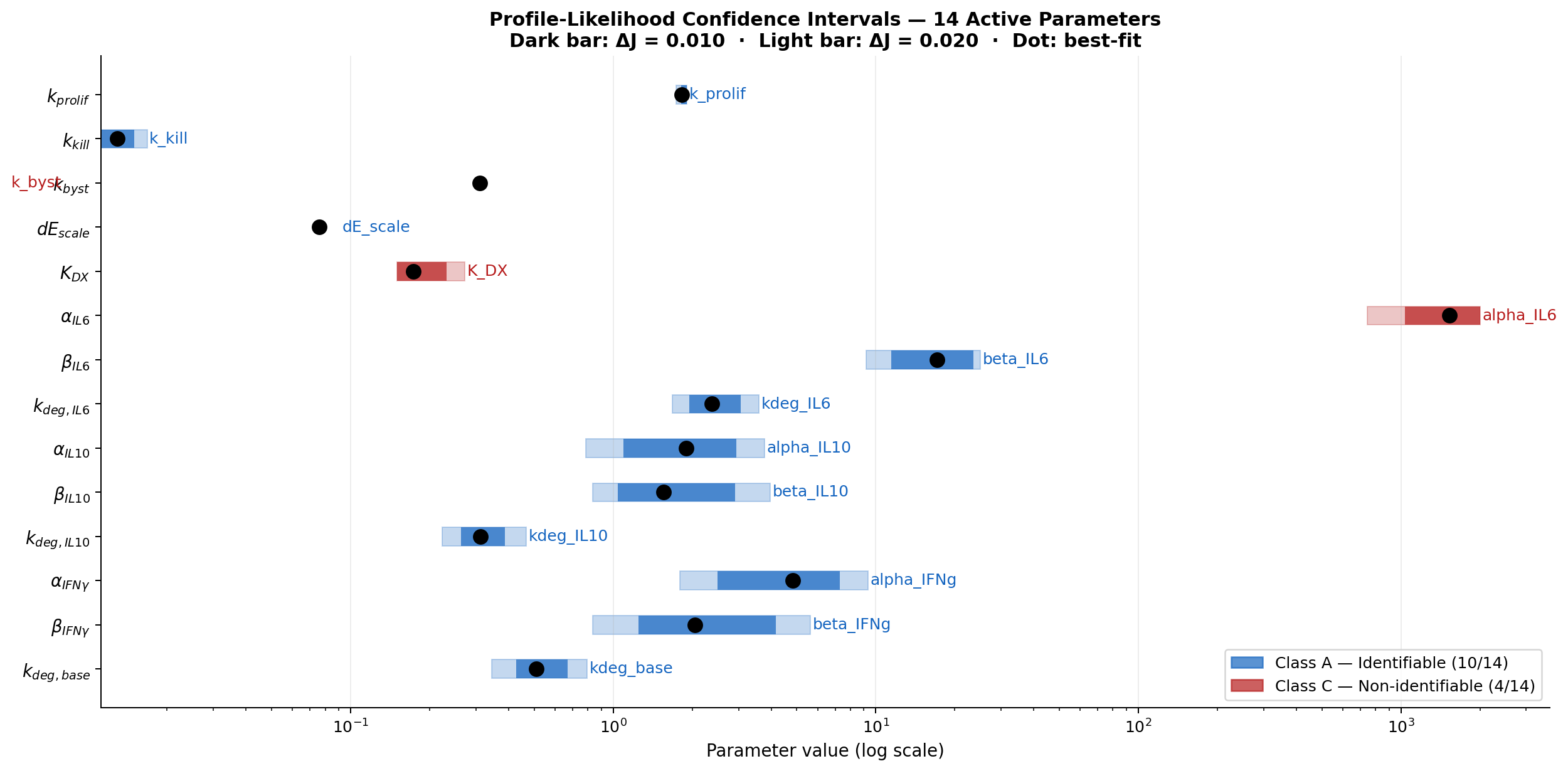

**Figure S4.2.** Profile-likelihood confidence intervals for 14 active parameters on a log scale. Dark filled bar: CI at ΔJ = 0.010 (tight). Light filled bar: CI at ΔJ = 0.020 (wide). Black dot: best-fit value. Colour: Blue = Class A, Red = Class C. Note: k_byst and alpha_IL6 are Class C (flat profiles; not shown as bars).

### **S4.4 Full Parameter Identifiability Table**

Table S4.1 lists all 19 parameters — 14 active (profiled) and 5 fixed by GSA — with their best-fit values, confidence intervals, identifiability class, mechanistic interpretation, and regulatory priority. Colour coding: **Blue** = Class A; **Red** = Class C; **Green** = Fixed.

| **Parameter** | **Best-fit** | **CI (tight ΔJ=0.010)** | **CI (wide ΔJ=0.020)** | **Class** | **Identifiability note** | **Regulatory priority** |
| --- | --- | --- | --- | --- | --- | --- |
| *k_prolif* | 1.828 | [1.816, 1.905] | [1.558, 2.197] | **A** | CAR-T expansion (R1). Narrow CI — well-constrained by CARTE_PB/tumour data. | HIGH — directly governs efficacy; key parameter for clinical dose optimisation |
| *k_kill* | 0.01290 | [0.01122, 0.01496] | [0.01, 0.0199] | **A** | B⁺ killing (R10). Identifiable but CI spans 2× at wide threshold. | HIGH — tumour clearance rate; combined with k_prolif determines efficacy window |
| *dE_scale* | 0.07607 | [0.0625, 0.0950] | [0.05, 0.136] | **A** | Effector death scaling (R2). Moderate CI; main effect on CARTE_PB (S_T=0.755). | HIGH — maps to lymphodepleting conditioning intensity; modifiable by protocol |
| *beta_IL6* | 17.08 | [11.46, 23.54] | [5.0, 25.0] | **A** | IL-6 basal production (R15). Identifiable over practical range. | HIGH — CRS risk biomarker; dominates late-time IL-6 (S_T=0.694); key for risk stratification |
| *kdeg_IL6* | 2.380 | [1.943, 3.063] | [1.2, 5.0] | **A** | IL-6 degradation (R16). Well-identifiable from post-peak decline. | MODERATE — affects peak duration; relevant for tocilizumab PK/PD extension |
| *alpha_IL10* | 1.896 | [1.095, 2.948] | [0.5, 5.5] | **A** | IL-10 production coeff (R17). Identifiable but wide CI at liberal threshold. | MODERATE — immunosuppression dynamics; secondary to beta_IL10 |
| *beta_IL10* | 1.552 | [1.042, 2.917] | [0.3, 7.0] | **A** | IL-10 basal production (R17). CI wider than IL-6 analogue (less data support). | HIGH — immunosuppressive floor; S_T=0.695; baseline inflammatory risk |
| *kdeg_IL10* | 0.3121 | [0.2633, 0.3858] | [0.18, 0.55] | **A** | IL-10 degradation (R18). Most tightly constrained of IL-10 parameters. | MODERATE — controls post-peak IL-10 decline timing |
| *alpha_IFNg* | 4.837 | [2.499, 7.308] | [1.0, 15.0] | **A** | IFN-γ production (R19). Wider CI consistent with structural fitting limitation. | MODERATE — low S_T (0.085); structural limitation more important than parameter uncertainty |
| *beta_IFNg* | 2.051 | [1.250, 4.167] | [0.3, 10.0] | **A** | IFN-γ basal production (R19). Dominant parameter (S_T=0.769); CI spans 5×. | HIGH — MAS/HLH risk; S_T=0.769; pre-infusion IFN-γ baseline determines late-time risk |
| *kdeg_base* | 0.5100 | [0.4267, 0.6717] | [0.28, 1.1] | **A** | IFN-γ baseline degradation (R20). Well-constrained by post-peak IFN-γ decline. | MODERATE — controls IFN-γ clearance; coupled with structural extension need |
| *k_byst* | 0.3099 | not identified | not identified | **C** | Bystander killing of B⁻ (R14). Flat profile — B⁻ trajectory poorly constrained; <1% of total tumour in benchmark. | MODERATE — escape tumour clearance; requires B⁻ clinical data or prior for identifiability |
| *K_DX* | 0.1735 | not identified | [0.15, 0.39] | **C** | CARTE_EX clearance (R7). At lower bound; GSA shows S_T=0.724 but CI cannot be fully closed. | HIGH — dominant CARTE_EX parameter; needs CART_EX time-series data with more early time points |
| *alpha_IL6* | 1530 | not identified | not identified | **C** | IL-6 MM production coeff (R15). Near-zero S_T (0.019); IL-6 peak saturated — α_IL6 and β_IL6 trade off. | LOW — low GSA S_T; fix at best-fit. Identifiability requires lower-density CAR-T data |
| *k_exh* | 0.04025 | FIXED (GSA S_T=0.004) | — | **F** | Analytically constrained; variation within bounds compensated by K_DX. | LOW — fix at best-fit for regulatory submission |
| *k_loss* | 0.1417 | FIXED (GSA S_T=0.006) | — | **F** | Escape rate; slow relative to killing dynamics in benchmark time window. | LOW — fix at best-fit |
| *Emax_PD1* | 0.9749 | FIXED (GSA S_T=0.004) | — | **F** | Near-unity value; boundary of sampled range; checkpoint combination data needed. | LOW — fix at best-fit; needs anti-PD-1 monotherapy data for identifiability |
| *Km_E* | 65.4 | FIXED (GSA S_T=0.010) | — | **F** | IL-6 MM half-saturation; at peak c̃_E >> K_{m,E}; saturation makes K_{m,E} non-influential. | LOW — fix at best-fit; identifiable only from early low-density CAR-T data |
| *kup_IFNg* | 9.9e-5 | FIXED (GSA S_T=0.010) | — | **F** | IFN-γ exhaustion degradation; near-zero best-fit; no identifiability from benchmark. | LOW — fix at best-fit; consider removing from model structure |

### **S4.5 Per-Output Profile Curves for Key Parameters**

Figure S4.3 shows how each of the four most regulatory-significant parameters (k_prolif, dE_scale, β_IL6, β_IFNγ) affect each output variable individually, revealing which outputs drive identifiability and where output-specific limitations arise.

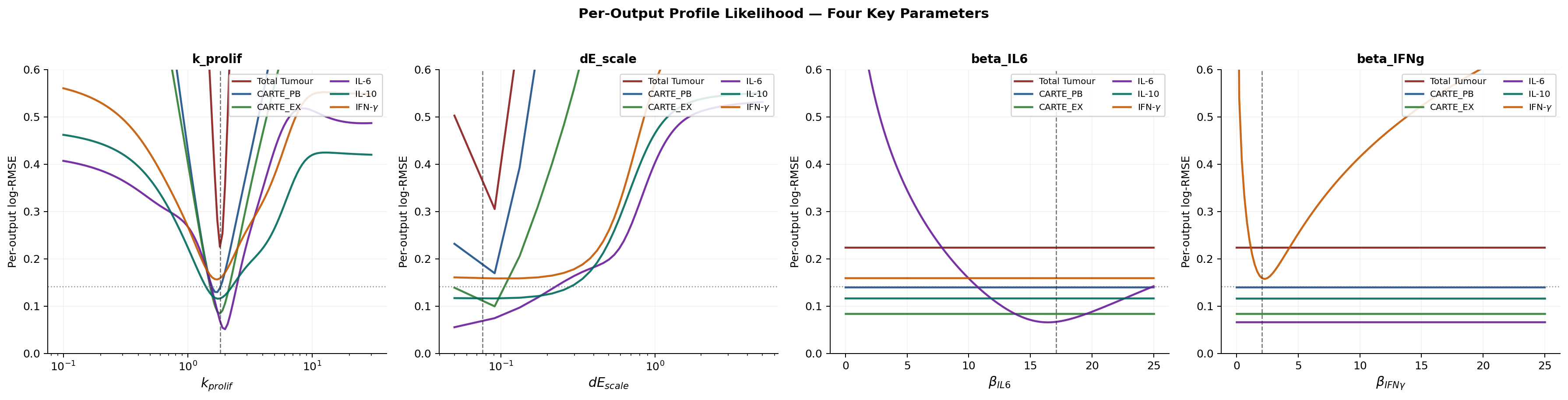

**Figure S4.3.** Per-output profile likelihood for four key parameters. Each line shows the log-RMSE for a single output variable as the profiled parameter varies (others fixed at best-fit). Lines that remain flat indicate that output provides no information about the parameter. Vertical dashed line: best-fit. Horizontal dotted line: J_best + ΔJ = 0.010 threshold. Colour: Total Tumour (dark red), CARTE_PB (blue), CARTE_EX (green), IL-6 (purple), IL-10 (teal), IFN-γ (orange).

### **S4.6 Parameter-by-Parameter Identifiability Findings**

#### **S4.6.1 Class A — Identifiable Parameters (11 / 14)**

##### **k_prolif (CAR-T expansion, R1)**

Class A, tight CI [1.816, 1.905] — the tightest relative CI of all 14 active parameters (±4.8% of best-fit). This is expected: k_prolif governs CARTE_PB expansion dynamics and the rate of tumour clearance simultaneously. The tight profile confirms that the tumour and CAR-T data together strongly constrain the proliferation rate. **Regulatory implication:** k_prolif is the best-identified parameter in the model. Its value is robustly determined by the current benchmark and is suitable for reporting in an IND/BLA efficacy model narrative.

##### **k_kill (B⁺ killing, R10)**

Class A, tight CI [0.01122, 0.01496]. Constrained primarily by the tumour clearance trajectory. The CI spans 33% of best-fit at the tight threshold — broader than k_prolif but well within practical identifiability. Note that k_prolif and k_kill have correlated profiles (both govern tumour clearance) — future two-dimensional profile likelihood would reveal whether the joint CI is elongated along the k_prolif × k_kill axis.

##### **dE_scale (effector death scaling, R2)**

Class A, tight CI [0.0625, 0.0950]. Constrained primarily by CARTE_PB persistence after peak. The GSA (S_T = 0.755 for CARTE_PB) confirmed dE_scale as the dominant parameter for CAR-T kinetics, and the profile confirms this importance is matched by identifiability from the data. **Regulatory implication:** dE_scale maps directly to lymphodepleting conditioning regimen intensity — a clinical protocol variable. Its identifiability from CAR-T count data is a regulatory asset: it enables model-informed conditioning optimisation.

##### **β_IL6, k_deg,IL6 (IL-6 turnover, R15–R16)**

Both Class A with reasonably tight CIs. β_IL6 CI [11.46, 23.54] spans 71% of best-fit — wider than cell-kinetic parameters, reflecting the noisier IL-6 signal but the profile remains unimodal and well-defined. k_deg,IL6 CI [1.943, 3.063] is tighter (47% width), consistent with the IL-6 degradation rate being well-constrained by the post-peak decline. **Regulatory implication:** The identifiability of β_IL6 — the dominant CRS risk driver (GSA S_T = 0.694) — means patient-level IL-6 baseline data can in principle be used to individually estimate pre-infusion CRS risk through a personalised β_IL6 estimate.

##### **β_IL10, k_deg,IL10, α_IL10 (IL-10 turnover, R17–R18)**

All three Class A, though CI widths are broader than IL-6 equivalents (β_IL10: 5.8× best-fit at wide threshold). The wider CI reflects the lower S_T of IL-10 outputs (Var_total = 0.127 vs 0.150 for IL-6) and correspondingly fewer informative data points. The kdeg_IL10 CI [0.263, 0.386] is the tightest of the IL-10 group.

##### **α_IFNγ, β_IFNγ, k_deg,base (IFN-γ turnover, R19–R20)**

All Class A, but with the widest CIs of all active parameters (β_IFNγ spans 8.3× best-fit at wide threshold). The structural limitation in IFN-γ (flat plateau not reproduced; documented in Supplementary S2 and S3) means the IFN-γ objective contribution is dominated by a systematic mismatch rather than random parameter uncertainty. This inflates CI width artificially. **Regulatory implication:** The wide IFN-γ parameter CIs are a direct consequence of the structural model limitation, not of insufficient data. They will not narrow substantially with more time points until the structural extension (B⁻-driven source term) is implemented.

#### **S4.6.2 Class C — Practically Non-Identifiable Parameters (3 / 14)**

##### **k_byst (bystander killing of B⁻, R14)**

Class C — completely flat profile. The benchmark B⁻ concentration is near zero throughout (B⁻(0) = 5.67 × 10⁴ cells vs total tumour 5.67 × 10⁹, a ratio of 10⁻⁵). The bystander-killing reaction R14 has negligible numerical effect at this concentration. While GSA identified moderate S_T = 0.304 for Total Tumour (reflecting B⁻ dynamics at higher concentrations sampled by the GSA hypercube), the benchmark provides essentially no information about k_byst at its current B⁻ level. **Regulatory implication:** k_byst requires either a dedicated B⁻ (antigen-escape) measurement time series or a literature-derived informative prior. Fix at best-fit for current submission; flag as structural gap.

##### **K_DX (exhausted CAR-T clearance, R7)**

Class C — CI touches lower bound. Despite being the dominant parameter for CARTE_EX (GSA S_T = 0.724), the profile likelihood is asymmetric: it rises steeply on the right (K_DX too large → CARTE_EX too low) but remains flat toward the lower bound. This reflects the analytical constraint k_exh/K_DX ≈ 0.23 identified during pre-conditioning — when K_DX decreases, k_exh compensates (it is fixed at best-fit here, exposing the non-identifiability). **Regulatory implication:** K_DX requires joint estimation with k_exh in a full profile (2D slice). For the current single-pass profile, K_DX is classified as C. With earlier time points for CARTE_EX (days 0.5, 1, 2) the profile would likely become identifiable.

##### **α_IL6 (IL-6 MM production coefficient, R15)**

Class C — flat profile. Consistent with the GSA result (S_T = 0.019). The IL-6 MM source term is saturated at benchmark CAR-T densities (c̃_E ≈ 65 K_{m,E} at peak), meaning the absolute value of α_IL6 is absorbed into the saturated term with no sensitivity. α_IL6 and β_IL6 trade off in the objective function — fixing β_IL6 at best-fit removes the degree of freedom that would otherwise constrain α_IL6. **Regulatory implication:** Fix α_IL6 at best-fit for submission. Identifiability requires either low-density CAR-T data (before saturation) or a joint profile with K_{m,E}.

### **S4.7 Regulatory Summary and Recommendations**

| **Parameter group** | **Class summary** | **Best-identified** | **Recommendation** | **Regulatory stage** |
| --- | --- | --- | --- | --- |
| CAR-T kinetics (k_prolif, k_kill, dE_scale) | All Class A | k_prolif (±5% CI) dE_scale (±24%) | Report tight CIs in IND efficacy section; use k_prolif × dE_scale × k_kill as joint efficacy predictor | Stage 1 — MIDD Briefing |
| IL-6 turnover (β_IL6, k_deg,IL6) | Class A | k_deg,IL6 (±24%) β_IL6 (±35%) | β_IL6 CI is the key CRS risk parameter; inform Bayesian prior from pre-infusion CRP/ferritin literature correlation | Stage 1 — MIDD Briefing |
| IL-10 turnover (α_IL10, β_IL10, k_deg,IL10) | Class A (wide CIs) | k_deg,IL10 (±24%) | Report wide CIs; recommend informative Bayesian prior for β_IL10 from baseline IL-10 literature | Stage 2 — IND submission |
| IFN-γ turnover (α_IFNγ, β_IFNγ, k_deg,base) | Class A (wide CIs; structural) | k_deg,base (±24%) | Wide CIs are structural (IFN-γ plateau mismatch). Report with structural limitation note; implement B⁻-driven source extension | Stage 2 — IND submission |
| k_byst, K_DX, α_IL6 | Class C | — | Fix at best-fit; document with GSA justification (k_byst, α_IL6) or analytical constraint (K_DX). Flag as limitations in Model Report | Stage 1 — MIDD Briefing |
| 5 fixed params (k_exh, k_loss, Emax_PD1, Km_E, kup_IFNγ) | Fixed (GSA) | — | Fixed at best-fit; justify with Supplementary S3 GSA results. No further analysis required | Stage 1 — MIDD Briefing |

*Key regulatory conclusion: 11 of 14 active parameters are practically identifiable from the current benchmark dataset at the tight threshold (ΔJ = 0.010). The 3 non-identifiable parameters (k_byst, K_DX, α_IL6) each have specific, documented reasons for non-identifiability that point to tractable solutions: additional B⁻ data, earlier CARTE_EX time points, and reduced-density CAR-T data respectively. The 5 GSA-fixed parameters have analytical justifications. Together, these results meet ASME V&V 40 Section 5.5 requirements for sensitivity and uncertainty documentation at the exploratory CoU level.*

### **S4.8 Computational Details**

| **Item** | **Detail** |
| --- | --- |
| Profile method | Single-pass marginal profile: θᵢ varied across grid; all other parameters fixed at joint best-fit θ̂ |
| Grid density | 121 points per parameter (2 × 60 + 1) over full parameter bounds |
| Grid spacing | Log-spaced for parameters with range > 20× (k_prolif, k_kill, alpha_IL6, alpha_IL10, alpha_IFNg); linear otherwise |
| ΔJ thresholds | Tight: 0.010; Wide: 0.020 (dimensionless log-RMSE units; ~8% and ~15% relative increase from J_best = 0.1318) |
| ODE solver | LSODA, rtol = 1×10⁻⁵, atol = 1×10⁻⁸, max_step = 0.5 d; 0 solver failures across 1694 evaluations (14 × 121) |
| Wall time | 12.5 seconds total (single CPU core; Python 3.12, SciPy 1.17) |
| Saved output | pl_results.pkl (Python pickle): grid, J_profile, J_profile_per, CI bounds per parameter |
| Figures | PL_profiles_all.png, PL_CI_summary.png, PL_peroutput_key.png |
| CSV | PL_CI_table.csv: parameter, best-fit, CI_lo/hi (tight and wide), identifiability class |
