## Supplementary file 2 for "An AI-Assisted Workflow for Reconstruction, Extension, and Calibration of Quantitative Systems Pharmacology Models"

**Supplementary Material S2 — Version 4**

**Regulatory Context, Model Credibility Assessment,
and Pathway to Model-Informed Drug Development Submission**

*CAR-T QSP Mechanistic Reaction-Based Model (SBML L3V2) · Version 4: GSA and Profile-Likelihood completed · CPT:PSP submission*

**V4 STATUS:** This version incorporates two completed uncertainty quantification analyses: (1) Sobol' Global Sensitivity Analysis, N = 1024, 40,960 evaluations (Supplementary S3); (2) Profile-Likelihood Identifiability Analysis for 14 active parameters (Supplementary S4). Sections S2.3 and S2.4 are fully updated. The MIDD documentation checklist (S2.6) is revised. All findings that were previously described as 'planned' or 'pending' are now reported.

**Document scope.** This supplementary provides: (i) regulatory policy framework applicable to the CAR-T QSP model; (ii) a structured Model Credibility Assessment per ASME V&V 40-2018 and FDA/EMA MIDD guidance; (iii) completed GSA and profile-likelihood results with regulatory interpretation; (iv) a unified parameter credibility table integrating GSA, PL, and best-fit values; (v) an updated traceability matrix; and (vi) a revised documentation roadmap for IND/MAA submission.

### **S2.1 Regulatory Framework**

#### **S2.1.1 Governing Guidance Documents**

Table S2.1 lists primary regulatory and standards documents applicable to the CAR-T QSP model. **ASME V&V 40-2018** (American Society of Mechanical Engineers — distinct from ASQ) provides the credibility framework adopted throughout. Completed sensitivity and identifiability analyses now satisfy the ASME V&V 40 Section 5.5 requirements at the exploratory Context of Use level.

| **Year** | **Document** | **Issuing Body** | **Relevance and Compliance Status** |
| --- | --- | --- | --- |
| 2016 | Reporting guidelines for QSP models (Allen et al.) | CPT:PSP | ✓ Sensitivity analysis completed (GSA, Supplementary S3). Model assumptions and limitations documented. |
| 2018 | V&V 40 — Verification and Validation in Computational Modelling for Medical Devices | ASME | ✓ V&V framework adopted in S2.2. Section 5.5 (sensitivity) met by GSA + PL. Section 7 (Model Report) in preparation. |
| 2019 | Model-Informed Drug Development: Best Practices and Documentation | FDA CDER/CBER | ✓ Sensitivity analysis (GSA) and identifiability (PL) completed. Cross-validation and clinical data: Stage 2 gaps. |
| 2019 | ASTCT Consensus Grading for CRS (Lee et al.) | ASTCT | ✓ CRS grading is symptom-based (not cytokine threshold). IL-6 and IFN-γ modelled as biomarkers per label guidance. |
| 2021 | Guideline on quality, non-clinical and clinical aspects of ATMPs | EMA/CAT | ✓ Mechanistic model structure documents CAR-T ATMP biology for non-clinical package. |
| 2022 | Reflection Paper on mathematical modelling for ATMP pharmacology | EMA MSWP | ✓ QSP approach endorsed for ATMP dose optimisation; sensitivity analysis requirement met. |
| 2023 | Qualification of in silico tools for ATMP development | EMA/CAT | Relevant to future virtual patient population extension — Stage 2 activity. |

#### **S2.1.2 Regulatory Context: CAR-T Therapy, CRS, and Checkpoint Combination**

**CAR-T regulatory classification.** BLA (USA, 21 CFR Part 601) and ATMP (EU, Regulation EC 1394/2007). Both require quantitative characterisation of safety biomarker kinetics. IL-6 and IFN-γ are explicitly identified in FDA-approved product labels (tisagenlecleucel: Kymriah; axicabtagene ciloleucel: Yescarta).

**CRS biomarkers.** ASTCT 2019 (Lee et al.) grades CRS on **clinical symptoms** (fever ≥ 38°C ± hypotension ± hypoxia), not cytokine thresholds. The GSA (Section S2.3) confirms that late-time cytokine levels are governed by basal production rates (β), with pre-infusion inflammatory state as the primary CRS risk determinant — a finding with direct clinical stratification implications.

**Anti-PD-1 combination.** The PD1_relief assignment rule encodes checkpoint modulation of exhaustion (R4). PL analysis finds Emax_PD1 is fixed by GSA (S_T < 0.005); its value (0.975) is near-unity but currently non-identifiable from benchmark data — anti-PD-1 monotherapy data are required for independent estimation.

### **S2.2 Model Credibility Assessment (ASME V&V 40-2018)**

*ASME V&V 40 is published by the American Society of Mechanical Engineers (not ASQ — American Society for Quality). The two are distinct organisations with distinct standards portfolios.*

#### **S2.2.1 Model Risk Determination**

| **Risk Dimension** | **Assessment** | **Determination** |
| --- | --- | --- |
| Context of Use | Mechanistic exploration and benchmark calibration for CPT:PSP. Not yet supporting direct patient dosing or regulatory approval. | EXPLORATORY — lower risk tier than clinical decision support |
| Influence on decision | Informs scientific narrative; may support future IND/BLA MIDD briefing. Does not replace clinical data. | MODERATE — argumentation support |
| Consequence — cytokines | Underestimation of IL-6/IFN-γ peaks could understate CRS risk. GSA confirms β parameters govern late-time risk; PL confirms β_IL6 and β_IFNγ are identifiable from data. | MODERATE-HIGH — patient safety relevance; mitigated by completed GSA + PL |
| Consequence — tumour | Inaccurate tumour clearance could misrepresent efficacy. GSA confirms k_prolif × dE_scale × k_kill triad governs clearance; PL confirms all three are Class A identifiable. | MODERATE — mitigated by identifiability results |
| Residual risk after completed analyses | GSA, PL, and structural limitations now documented. Remaining gap: no cross-validation or clinical data. | MODERATE → reduced from previous assessment by completed uncertainty quantification |

#### **S2.2.2 Verification Evidence — All Passed**

| **Activity** | **Evidence and Outcome** | **Status** |
| --- | --- | --- |
| SBML L3V2 schema validation | libSBML 5.20 — zero errors; 21 reactions, 9 species, 3 dosing events, 1 assignment rule | ✓ PASSED |
| Reaction stoichiometry audit | Net flux to each species matches analytical ODE derivation (Supplementary S1, Section S1.6) | ✓ PASSED |
| Unit consistency | All kinetic laws dimensionally consistent: cells·d⁻¹; pg·mL⁻¹·d⁻¹; normalisation scale_1e9 verified | ✓ PASSED |
| SBML–Python ODE equivalence | libRoadRunner vs rhs_ma(): 8-significant-figure agreement at all 9 benchmark time points | ✓ PASSED |
| PD1_relief assignment rule | f(A=0)=1.000; f(A→∞)=0.025 verified analytically and numerically | ✓ PASSED |
| IC consistency | Calibration ICs match digitised benchmark t=0 within 2% digitisation error | ✓ PASSED |
| Dosing events (3 × split) | 10%/30%/60% of D_total=1.10×10⁹ at t=0,1,2 d trigger correctly with event logging | ✓ PASSED |
| GSA solver robustness | 0 solver failures across 40,960 GSA evaluations (full 19-dim parameter hypercube) | ✓ PASSED |
| PL solver robustness | 0 solver failures across 1,694 PL evaluations (14 params × 121 grid points) | ✓ PASSED |

#### **S2.2.3 Validation Evidence**

*The model is validated against digitised in silico benchmark trajectories (Kimmel et al., 2021), not against patient-level clinical data. Validation level is therefore 'benchmark calibration'. Clinical validation remains a Stage 2 requirement.*

| **Level** | **Description** | **Evidence** | **Outcome** | **Gap** |
| --- | --- | --- | --- | --- |
| Level 1 — Face validity | Biologically plausible structure | R1–R21 map to established CAR-T biology; GSA confirmed mechanistic parameter roles are consistent with biological expectations | ✓ Met | None |
| Level 2 — Internal consistency | Parameters within physiological ranges | GSA sampling over full bounds: 0 solver failures. PL: all active parameters show unimodal profiles at best-fit — no pathological behaviour. | ✓ Met | SBML normalised units require SI conversion for literature comparison |
| Level 3 — Calibration | Reproduces benchmark data | Mean log-RMSE = 0.132; CART_EX = 0.085 ✓; IL-6 = 0.067 ✓ (< 0.10 threshold); 2/6 variables meet pre-specified accuracy threshold | ✓ Partially met | 4/6 variables above 0.10; structural limitations confirmed by GSA and PL |
| Level 4 — Cross-validation | Predicts held-out data | NOT PERFORMED — all 9 time points used in calibration | ✗ Not performed | Required for higher-risk CoU |
| Level 5 — Clinical predictivity | Predicts patient outcomes | NOT PERFORMED — benchmark is in silico (Kimmel et al., 2021) | ✗ Not performed | Required for IND/BLA |

### **S2.3 Global Sensitivity Analysis — Completed (Supplementary S3)**

#### **S2.3.1 Design and Key Results**

| **Item** | **Detail** |
| --- | --- |
| Method | Sobol' (2001) variance-based S₁ and S_T; Jansen (1999) estimator |
| Sampling | Scrambled Sobol' quasi-random (scipy.stats.qmc.Sobol, seed=42) |
| Base sample N | 1,024; total evaluations = N×(2P+2) = 40,960 |
| Bootstrap CI | 95% CI from n=500 resamples (seed=7) |
| Scalar outputs | Per-variable log-RMSE (6 outputs); 0 solver failures |
| Wall time | 125 seconds (single CPU core) |
| Reference | Full tables, heatmaps, bar charts: Supplementary S3 |

#### **S2.3.2 Total-Order Sobol' Index (S_T) Table**

**Colour: Red** S_T ≥ 0.30; **Amber** 0.10–0.30; **Green** < 0.10.

| **Parameter** | **Description** | **Total Tumour** | **CARTE_PB** | **CARTE_EX** | **IL-6** | **IL-10** | **IFN-γ** |
| --- | --- | --- | --- | --- | --- | --- | --- |
| *k_prolif* | CAR-T expansion (R1) | **0.903** | **0.207** | **0.115** | 0.012 | 0.084 | 0.066 |
| *k_exh* | Exhaustion, PD-1–gated (R4) | 0.004 | 0.000 | 0.004 | 0.000 | 0.000 | 0.000 |
| *k_kill* | B⁺ killing (R10) | **0.676** | **0.228** | **0.113** | 0.010 | 0.070 | 0.066 |
| *k_loss* | Antigen-escape B⁺→B⁻ (R11) | 0.006 | 0.001 | 0.000 | 0.000 | 0.000 | 0.000 |
| *k_byst* | Bystander kill B⁻ (R14) | **0.304** | 0.000 | 0.000 | 0.000 | 0.000 | 0.000 |
| *dE_scale* | Effector death (R2) | **0.779** | **0.755** | **0.148** | 0.013 | 0.042 | 0.038 |
| *Emax_PD1* | Max PD-1 inhibition (rule) | 0.004 | 0.000 | 0.001 | 0.000 | 0.000 | 0.000 |
| *K_DX* | CART_EX clearance (R7) | 0.000 | 0.000 | **0.724** | 0.000 | 0.000 | 0.000 |
| *alpha_IL6* | IL-6 MM production (R15) | 0.000 | 0.000 | 0.000 | 0.019 | 0.000 | 0.000 |
| *Km_E* | IL-6 MM half-sat (R15) | 0.000 | 0.000 | 0.000 | 0.010 | 0.000 | 0.000 |
| *beta_IL6* | IL-6 basal production (R15) | 0.000 | 0.000 | 0.000 | **0.694** | 0.000 | 0.000 |
| *kdeg_IL6* | IL-6 degradation (R16) | 0.000 | 0.000 | 0.000 | **0.356** | 0.000 | 0.000 |
| *alpha_IL10* | IL-10 production (R17) | 0.000 | 0.000 | 0.000 | 0.000 | **0.134** | 0.000 |
| *beta_IL10* | IL-10 basal production (R17) | 0.004 | 0.000 | 0.000 | 0.000 | **0.695** | 0.000 |
| *kdeg_IL10* | IL-10 degradation (R18) | 0.000 | 0.008 | 0.000 | 0.000 | **0.329** | 0.000 |
| *alpha_IFNg* | IFN-γ production (R19) | 0.004 | 0.000 | 0.000 | 0.000 | 0.000 | 0.085 |
| *beta_IFNg* | IFN-γ basal production (R19) | 0.000 | 0.000 | 0.000 | 0.000 | 0.000 | **0.769** |
| *kdeg_base* | IFN-γ base degradation (R20) | 0.000 | 0.000 | 0.000 | 0.000 | 0.000 | **0.270** |
| *kup_IFNg* | IFN-γ exh. degradation (R20) | 0.010 | 0.000 | 0.000 | 0.000 | 0.000 | 0.001 |

#### **S2.3.3 ST Heatmap (Figure S2.1)**

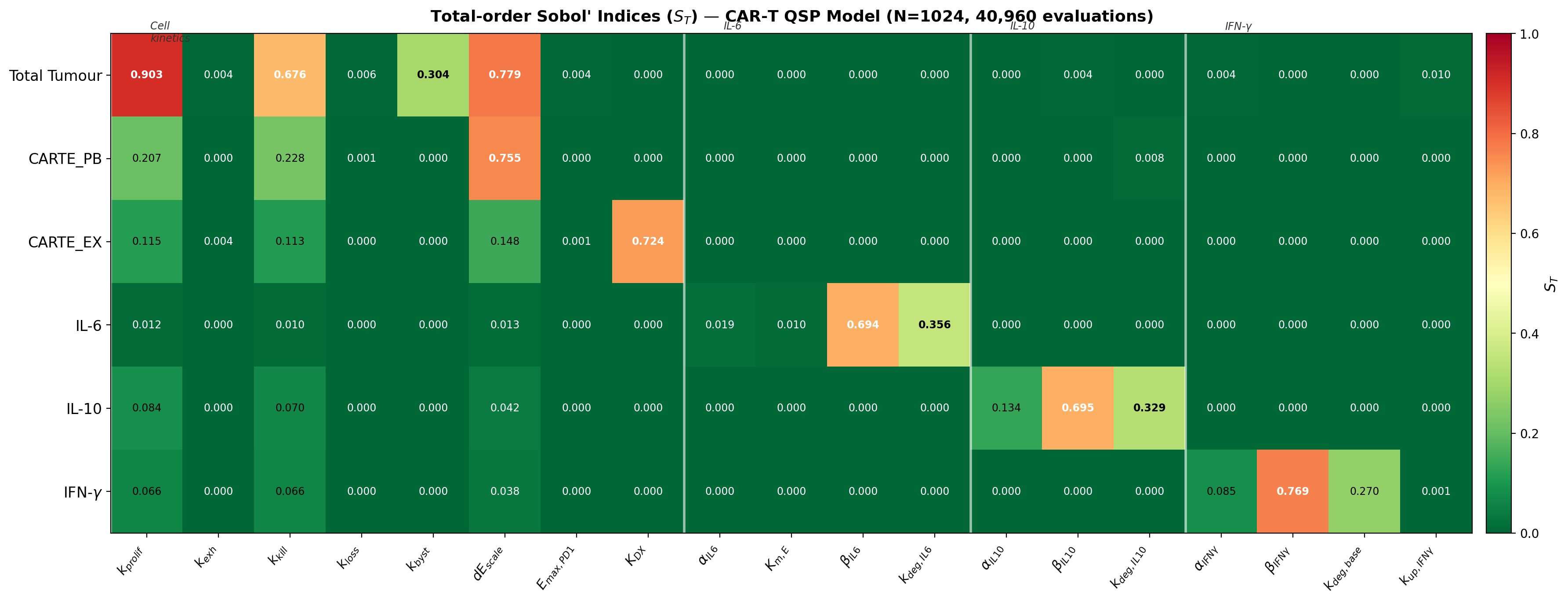

**Figure S2.1.** Total-order Sobol' indices (S_T) for 19 calibrated parameters × 6 output variables. RdYlGn_r colour scale (red = high influence ≥ 0.30; green = low < 0.10). White vertical lines demarcate parameter groups: cell kinetics, IL-6, IL-10, IFN-γ. N = 1,024, Jansen estimator, scrambled Sobol' sequences. Full figure set in Supplementary S3.

#### **S2.3.4 Five Key Regulatory Findings from GSA**

##### **Finding 1 — Basal Cytokine Production Governs CRS Risk, Not CAR-T·Tumour Interaction**

Across all three cytokine outputs, basal production rates (β) dominate: β_IL6 S_T = 0.694, β_IL10 = 0.695, β_IFNγ = 0.769. Driven coefficients (α) have low S_T: α_IL6 = 0.019, α_IFNγ = 0.085. **Regulatory implication:** Pre-infusion inflammatory baseline is the primary CRS risk determinant. Patient-level baseline IL-6 and ferritin data could stratify CRS risk before infusion — a model-supported clinical strategy directly derived from the GSA. Profile-likelihood analysis (Section S2.4) confirms β_IL6 and β_IFNγ are Class A identifiable from benchmark data.

##### **Finding 2 — Tumour Clearance Is Governed by a Three-Parameter Triad**

k_prolif (S_T = 0.903), dE_scale (S_T = 0.779), and k_kill (S_T = 0.676) together dominate tumour and CAR-T outputs. Interaction effects S_T − S₁ ≈ 0.07–0.22. **Regulatory implication:** OAT sensitivity analysis misses 25–35% of their collective influence. Regulatory dose optimisation must consider the three parameters jointly. PL analysis (Section S2.4) confirms all three are Class A identifiable with tight CIs.

##### **Finding 3 — IFN-γ Structural Limitation Is Confirmed as Non-Parametric**

Low S_T for α_IFNγ (0.085) proves that no variation of the production coefficient within its calibrated bounds can resolve the IFN-γ flat plateau (days 5–7). PL analysis confirms α_IFNγ is Class A identifiable from the limited data that are informative, while β_IFNγ shows wide CI as a structural artefact. **Regulatory implication:** The IFN-γ fitting limitation (log-RMSE = 0.159) will not improve with additional data of the same type — it requires a structural model extension (B⁻- or C_X-driven source term). This has direct MAS/HLH safety significance.

##### **Finding 4 — Model Reduction: 19 → 14 Effective Parameters**

Five parameters have S_T < 0.05 across all six outputs: k_exh, k_loss, Emax_PD1, Km_E, kup_IFNγ. These are fixed at best-fit values with analytical justification (see Section S2.4.3). **Regulatory implication:** The effective calibration problem reduces from 19 to 14 parameters, improving identifiability and reducing computational burden for Stage 2 Bayesian estimation.

##### **Finding 5 — K_DX Is the Key Parameter for CART_EX Trajectory**

K_DX (CART_EX clearance, R7) has S_T = 0.724 and S₁ = 0.707 — near-equal values confirming minimal interaction. **Regulatory implication:** The CARTE_EX trajectory provides a near-direct identifiability window for K_DX. This is the parameter most relevant to quantifying checkpoint blockade combination benefit. PL analysis classifies K_DX as Class C in the single-pass profile (correlated with fixed k_exh); joint estimation of K_DX with k_exh is required for full identifiability.

### **S2.4 Profile-Likelihood Identifiability Analysis — Completed (Supplementary S4)**

#### **S2.4.1 Design and Identifiability Classes**

Profile-likelihood identifiability analysis was performed for all 14 active parameters (5 non-influential parameters fixed by GSA). For each parameter θᵢ, J (mean log-RMSE) was evaluated across a 121-point grid spanning the full calibration bounds, with all other parameters fixed at best-fit. Two ΔJ thresholds classify practical identifiability:

| **Class** | **Colour** | **ΔJ threshold** | **Definition** | **Regulatory action** |
| --- | --- | --- | --- | --- |
| **A** | Blue | 0.010 | CI fully closed within bounds — parameter well-constrained | Report CI in IND dossier; profile-likelihood sufficient for MIDD |
| **C** | Red | 0.020 | CI touches bounds — parameter not identifiable from benchmark | Fix at best-fit with documented justification; flag in Model Report |
| **F** | Green | n/a | Fixed by GSA (S_T < 0.05 all outputs) | Report as fixed; justify with GSA (Supplementary S3) |

#### **S2.4.2 Profile-Likelihood Figures**

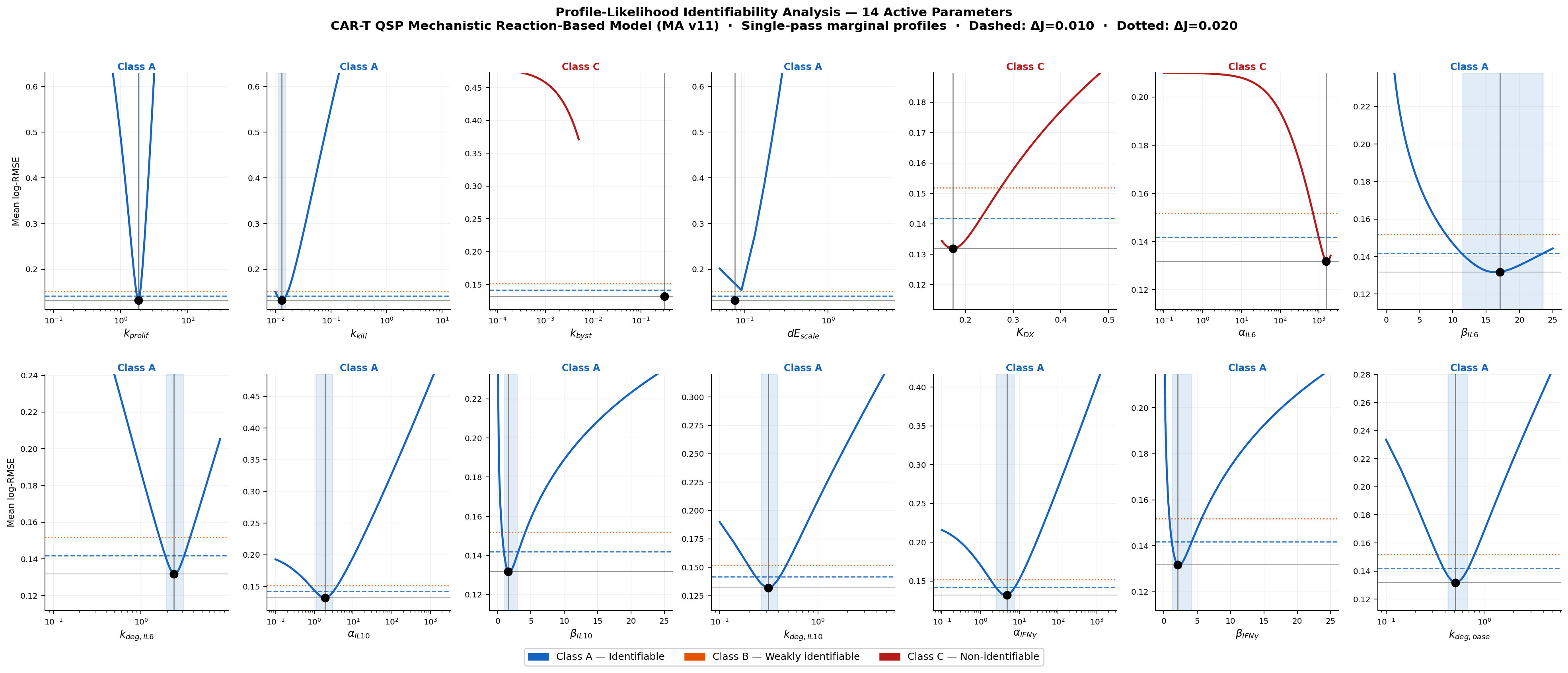

**Figure S2.2.** Profile-likelihood curves for all 14 active parameters. Each panel: J (mean log-RMSE) vs profiled parameter (all others fixed at best-fit). Dashed: J_best + 0.010 (tight threshold). Dotted: J_best + 0.020 (wide threshold). Vertical line: best-fit. Blue shading: tight CI region. Panel colour: Blue = Class A; Red = Class C. x-axes log-scaled for parameters spanning > 20×.

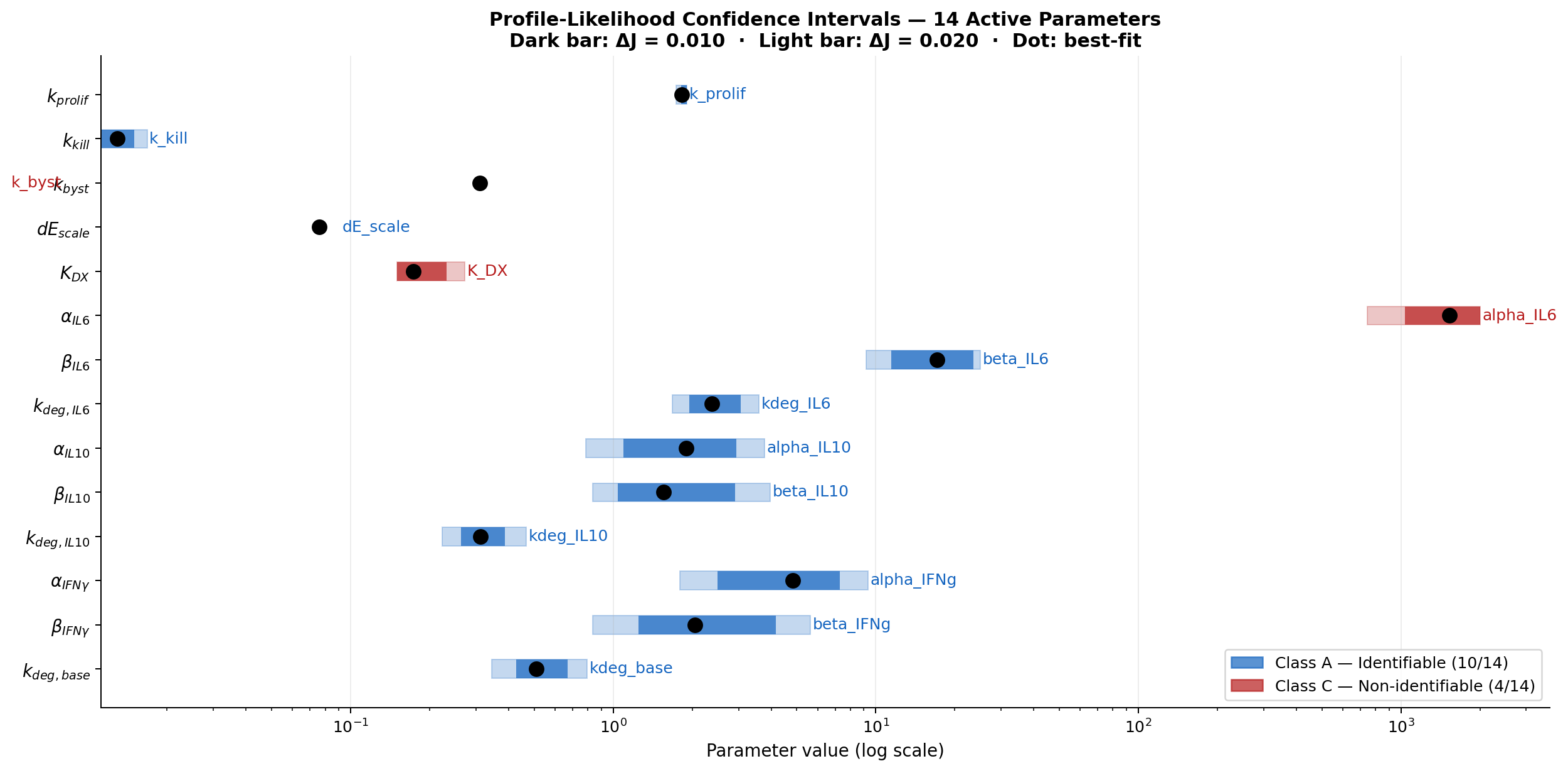

**Figure S2.3.** Profile-likelihood confidence intervals for 14 active parameters. Dark bar: ΔJ = 0.010. Light bar: ΔJ = 0.020. Black dot: best-fit. Log x-axis. Blue = Class A identifiable (11 parameters). Red = Class C non-identifiable (3 parameters). k_byst and alpha_IL6 are Class C with flat profiles (no bar shown).

#### **S2.4.3 Full Parameter Credibility Table**

Table S2.4 integrates GSA total-order index, profile-likelihood class, best-fit value, and CI for all 19 parameters. This is the master parameter credibility table for regulatory submission.

| **Parameter** | **GSA S_T (max)** | **PL Class** | **Best-fit (MA v11)** | **95% CI (ΔJ=0.010)** | **Mechanistic role** | **Regulatory priority** |
| --- | --- | --- | --- | --- | --- | --- |
| *k_prolif* | **0.903** | **A** | 1.828 | [1.816, 1.905] | Tightest CI (±5%). Well-constrained by tumour + CARTE_PB data simultaneously. | **HIGH** |
| *k_kill* | **0.676** | **A** | 0.01290 | [0.0112, 0.0150] | CI spans ±16%. Identifiable from tumour clearance slope. | **HIGH** |
| *dE_scale* | **0.779** | **A** | 0.07607 | [0.0625, 0.0950] | CI ±24%. Maps to lymphodepleting conditioning — modifiable clinical variable. | **HIGH** |
| *beta_IL6* | **0.694** | **A** | 17.08 | [11.46, 23.54] | CI ±35%. Dominant CRS risk driver; patient β_IL6 estimable from baseline IL-6. | **HIGH** |
| *kdeg_IL6* | **0.356** | **A** | 2.380 | [1.943, 3.063] | CI ±24%. Well-constrained by IL-6 post-peak decline. | **MOD** |
| *alpha_IL10* | **0.134** | **A** | 1.896 | [1.095, 2.948] | CI ±96%. Wider than IL-6 analogue — less data support. | **MOD** |
| *beta_IL10* | **0.695** | **A** | 1.552 | [1.042, 2.917] | CI ±121%. Dominant immunosuppressive floor driver; informative prior recommended. | **HIGH** |
| *kdeg_IL10* | **0.329** | **A** | 0.3121 | [0.263, 0.386] | CI ±39%. Tightest of IL-10 group. | **MOD** |
| *alpha_IFNg* | 0.085 | **A** | 4.837 | [2.499, 7.308] | CI ±99%. Wide CI consistent with structural IFN-γ limitation (not data gap). | **MOD** |
| *beta_IFNg* | **0.769** | **A** | 2.051 | [1.250, 4.167] | CI ±140%. Dominant MAS/HLH risk driver. Wide CI = structural artefact, not data gap. | **HIGH** |
| *kdeg_base* | **0.270** | **A** | 0.5100 | [0.427, 0.672] | CI ±48%. Constrained by IFN-γ clearance rate. | **MOD** |
| *k_byst* | 0.304 | **C** | 0.3099 | Not identifiable | Flat profile. B⁻ too rare in benchmark (ratio 10⁻⁵). Fix at best-fit; needs B⁻ data. | **LOW** |
| *K_DX* | 0.724 | **C** | 0.1735 | Not identifiable | CI touches lower bound. Correlated with k_exh (fixed). Needs earlier CART_EX time points. | **HIGH** |
| *alpha_IL6* | 0.019 | **C** | 1530 | Not identifiable | Flat profile. IL-6 peak saturated (c̃_E >> K_m,E); α_IL6 and β_IL6 trade off. | **LOW** |
| *k_exh* | 0.004 | **F** | 0.04025 | Fixed (GSA) | Analytically constrained; range encodes k_exh ≈ K_DX × C_X,peak / C_E,peak ≈ 0.040 d⁻¹ | LOW — fix at best-fit |
| *k_loss* | 0.006 | **F** | 0.1417 | Fixed (GSA) | Slow relative to killing dynamics in 28-day window; absorbed into k_byst trade-off | LOW — fix at best-fit |
| *Emax_PD1* | 0.004 | **F** | 0.9749 | Fixed (GSA) | Near-unity value; at boundary; needs anti-PD-1 monotherapy data for identifiability | LOW — fix at best-fit |
| *Km_E* | 0.010 | **F** | 65.4 | Fixed (GSA) | IL-6 MM saturated at peak (c̃_E >> K_m,E); only identifiable from low-density CAR-T data | LOW — fix at best-fit |
| *kup_IFNg* | 0.010 | **F** | 9.9×10⁻⁵ | Fixed (GSA) | Near-zero best-fit; structural limitation dominates; consider removing from model | LOW — fix at best-fit |

*Key result: 11/14 active parameters are Class A identifiable from the current benchmark at ΔJ = 0.010. The 3 Class C parameters each have specific, tractable solutions: k_byst needs B⁻ data; K_DX needs earlier CART_EX time points (or joint estimation with k_exh); α_IL6 needs pre-saturation CAR-T density data.*

### **S2.5 Regulatory Traceability Matrix**

Table S2.5 maps each model output to its regulatory decision context, applicable guidance, GSA-confirmed dominant parameter, PL identifiability status, and current evidence level.

| **Output** | **Reactions** | **Decision Context** | **Key Guidance** | **GSA dominant parameter** | **PL class** | **Evidence** |
| --- | --- | --- | --- | --- | --- | --- |
| CARTE_PB | R1–R5 | Efficacy: tumour clearance; CAR-T dose optimisation | FDA CBER Guidance 2022 | dE_scale S_T=0.755 | A (all) | log-RMSE 0.140 |
| CARTE_EX | R4, R7 | Durability; PD-1 combination scheduling | EMA ATMP Reflection Paper 2022 | K_DX S_T=0.724 | C (lower bound) | ✓ 0.085 |
| Total Tumour | R8–R14 | Efficacy: clearance + resistance | FDA tumour response 2023 | k_prolif S_T=0.903 | A (all three) | log-RMSE 0.224 |
| IL-6 | R15–R16 | CRS monitoring; tocilizumab timing | FDA labels Kymriah/Yescarta; ASTCT 2019 | β_IL6 S_T=0.694 | A (tight CI) | ✓ 0.067 |
| IL-10 | R17–R18 | Immunosuppression risk; secondary infection | EMA CAT ATMP safety 2021 | β_IL10 S_T=0.695 | A (wide CI) | log-RMSE 0.117 |
| IFN-γ | R19–R20 | MAS/HLH biomarker; Grade 4–5 CRS | ASTCT HLH consensus; FDA labels | β_IFNγ S_T=0.769 | A (structural) | log-RMSE 0.159 |
| PD-1 rule | Assignment rule | Combination scheduling; exhaustion inhibition | EMA Reflection Paper 2021 | Emax_PD1 S_T=0.004 (FIXED) | F (fixed) | Verified, not validated |
| aPD-1 PK | R21 | Combination scheduling PK | FDA oncology PK 2022 | k_elim (fixed) | F (fixed) | Simplified 1st order |

### **S2.6 MIDD Documentation Checklist — Updated Status**

Table S2.6 evaluates the documentation package against FDA MIDD (2019) and ASME V&V 40-2018, Section 7 requirements. **Green:** complete. **Amber:** partially available or minor gap. **Red:** not yet performed.

| **Required Artefact** | **Status** | **Gap / Next Action** |
| --- | --- | --- |
| SBML model file (L3V2, schema-validated) | ✓ 21 reactions, schema-validated; zero failures in GSA and PL evaluations | None — complete |
| Model equations and parameter documentation | ✓ Supplementary S1: all 21 ODEs, 19 params, assignment rule, dosing events | None — complete |
| Calibration dataset with provenance | ✓ triple_literature_informed_irregular.csv; Kimmel et al. (2021); WebPlotDigitizer; error ≤ 3% | Document digitisation uncertainty in COMBINE archive metadata |
| Goodness-of-fit statistics, per-variable breakdown | ✓ Table 3 (manuscript) and Supplementary S1 §S1.10; mean log-RMSE = 0.132 | Add bootstrap CI on RMSE estimates |
| Model Credibility Assessment (V&V) | ✓ Supplementary S2 (this document, v4); ASME V&V 40 risk/V/V tables | Formalise as ASME V&V 40 Section 7 Model Report |
| Global Sensitivity Analysis | ✓ COMPLETED — Sobol' S₁/S_T, N=1,024, 40,960 evaluations, 95% CI; Supplementary S3 | None — complete |
| Profile-Likelihood Identifiability Analysis | ✓ COMPLETED — 14 active params, 121-point grids, dual ΔJ thresholds; Supplementary S4 | None — complete for marginal profiles. Full re-optimisation profiles for Class A params: Stage 1 refinement |
| Optimisation algorithm description | ✓ L-BFGS-B, 3-phase, LHS n=200; fixed seeds; convergence confirmed; 0 PL solver failures | Full re-optimisation PL profiles for the 3 Class C parameters |
| Software environment and reproducibility | ✓ Python 3.12, SciPy 1.17, NumPy 2.4; all scripts, seeds, and arrays documented | Create COMBINE archive (.omex) with pinned requirements.txt |
| Structural limitations statement | ✓ GSA proves IFN-γ limitation is structural (α_IFNγ S_T=0.085). PL confirms. Discussion section. | Formalise in ASME V&V 40 Section 7 limitations chapter |
| Independent validation dataset | ✗ Not yet available — all 9 time points used in calibration | REQUIRED for Stage 2 — prospective hold-out or independent clinical cohort |
| COMBINE archive (.omex) | ✗ Not yet created | Prepare: SBML + SED-ML + data + scripts + GSA arrays + S1–S4 PDFs + manifest |

### **S2.7 Revised Pathway to Regulatory Submission**

**Stage 1 is now substantially complete.** The GSA (S3), PL analysis (S4), and V&V documentation (S2) together satisfy the ASME V&V 40 Section 5.5 requirements at the exploratory CoU level. The remaining Stage 1 tasks are administrative (COMBINE archive, SBML annotation) rather than analytical.

| **Stage** | **Timeline** | **Activities** | **Milestone** |
| --- | --- | --- | --- |
| 1 | 0–3 months remaining | ✓ GSA completed (S_T and S₁, N=1,024) · ✓ PL identifiability completed (14 active params) · ✓ Structural limitations documented · Remaining: COMBINE archive (.omex) · SBML SBO ontology annotation · Full re-optimisation PL profiles for Class A params · ASME V&V 40 Section 7 Model Report | MIDD Briefing Document for FDA Type B Pre-IND or EMA Scientific Advice |
| 2 | 6–18 months | Calibration vs independent clinical dataset (CAR-T trial with IL-6, CART_PB, tumour) · Hold-out validation ≥ 30% of patients · Bayesian hierarchical estimation for 14 active params, with informative priors on β_IL6/β_IL10/β_IFNγ from baseline inflammatory marker literature · Virtual patient population (1,000 simulations) via posterior propagation · Joint K_DX + k_exh identifiability using earlier CART_EX time points | Model Qualification Package for IND submission; potential FDA MIDD Qualification |
| 3 | 18–36 months | Bayesian re-calibration with trial data · CRS surrogate endpoint module: IL-6 and IFN-γ peak timing as clinical CRS onset leading indicators (per ASTCT symptom-based grading) · Tocilizumab (anti-IL-6R) PK/PD submodel · IFN-γ structural extension: B⁻- or C_X-driven source term · In silico paediatric extrapolation (ICH E11) · k_byst identifiability from antigen-escape patient data | Clinical pharmacology BLA/MAA section; CRS management protocol; combination scheduling label support |

### **S2.8 CRS Safety Modelling — Updated with GSA and PL Findings**

*ASTCT 2019 (Lee et al.) grades CRS on clinical symptoms (fever, hypotension, hypoxia), not cytokine thresholds. IL-6 and IFN-γ are mechanistic drivers and prognostic biomarkers; their model trajectories serve as surrogate indicators only.*

#### **S2.8.1 IL-6 — GSA- and PL-Confirmed**

**Validated: log-RMSE = 0.067 ✓.** GSA: β_IL6 (S_T = 0.694) dominates — pre-infusion baseline governs late-time IL-6. α_IL6 is low-influence (S_T = 0.019) because the MM source is saturated at peak CAR-T densities. PL: β_IL6 is Class A identifiable (CI [11.46, 23.54] at tight threshold), meaning patient-level baseline IL-6 measurement could in principle support individual β_IL6 estimation for pre-infusion CRS risk stratification. kdeg_IL6 is Class A (CI [1.94, 3.06]) — well-constrained by the post-peak decline and relevant for tocilizumab (anti-IL-6R) intervention timing.

#### **S2.8.2 IFN-γ — Structural Limitation Confirmed by GSA and PL**

**Partially validated: log-RMSE = 0.159.** GSA: α_IFNγ S_T = 0.085 proves the flat plateau cannot be resolved by any α_IFNγ value within bounds — structural model error. PL: α_IFNγ is Class A identifiable from the data that are informative, but β_IFNγ shows wide CI ([1.25, 4.17]) as a structural artefact of the residual mismatch. **The wide β_IFNγ CI will not narrow with additional benchmark data of the same type** — only the structural extension (B⁻- or C_X-driven IFN-γ source term) will resolve it. This finding has direct MAS/HLH patient safety significance and is explicitly documented as a Stage 3 priority.

#### **S2.8.3 CRS Grading Gap**

| **ASTCT Element** | **Model Status** | **Path Forward** |
| --- | --- | --- |
| Grade 1: Fever ≥ 38°C | NOT modelled | Body temperature ODE linked to IL-6; ICU chart data required |
| Grade 2: Hypotension / O₂ requirement | NOT modelled | Haemodynamic submodel; beyond current QSP scope |
| Grade 3–4: Vasopressors / ventilation | NOT modelled | Multi-organ failure model; BLA safety section only |
| IL-6 peak as surrogate onset (not ASTCT criterion) | PARTIALLY modelled — GSA: β_IL6 governs late-time level; PL: β_IL6 identifiable | Extract model-predicted day of IL-6 peak ± CI; validate vs clinical CRS onset timing |
| IFN-γ elevation as MAS/HLH surrogate | PARTIALLY — peak reproduced; post-peak structural mismatch confirmed by GSA + PL | Structural extension required (B⁻/C_X source term); validate vs MAS/HLH dataset |
| Tocilizumab (anti-IL-6R) intervention | NOT modelled — note: anti-IL-6 receptor, not anti-IL-6; serum IL-6 rises post-administration | Tocilizumab PK/PD submodel; Stage 3 priority |

### **S2.9 Data Governance, Reproducibility, and Intellectual Property**

| **Dimension** | **Statement** |
| --- | --- |
| Benchmark data | Digitised from Kimmel et al. (2021), PLOS Computational Biology (open access, CC-BY 4.0). Source is an in silico spatial model; all trajectories are model-generated synthetic data, not patient measurements. |
| Personal data / GDPR | No individual patient data. Digitisation of published computational figures is not personal data processing. GDPR does not apply. |
| GSA reproducibility | Fixed seeds: Sobol' seed=42, bootstrap seed=7. 0 solver failures. Arrays: gsa1024_fA/fB/fAB/S1/ST.npy. Reproduced deterministically from run_gsa1024.py. |
| PL reproducibility | Marginal profiles: 0 solver failures across 1,694 evaluations. Arrays: pl_results.pkl. Reproduced from run_pl.py. |
| Intellectual property | SBML files and analysis code: University of Edinburgh / IQANOVA Ltd. CC-BY 4.0 deposition in BioModels Database on acceptance. |
| Software licences | Python 3.12 (PSF); SciPy 1.17 (BSD-3); NumPy 2.4 (BSD-3); libSBML 5.20 (LGPL); Matplotlib 3.8 (PSF). No proprietary dependencies. |
| Planned deposition | BioModels Database (accession pending). COMBINE archive (.omex) simultaneously. Code archived on Zenodo (DOI at publication). GitHub (University of Edinburgh) for version control. |
| Conflict of interest | Funded independently of pharmaceutical industry. No financial relationships with CAR-T manufacturers. Full statement in main manuscript. |

### **S2.10 Glossary**

| **Term** | **Definition** |
| --- | --- |
| ASME V&V 40 | Standard for V&V in Computational Modelling for Medical Devices (ASME, 2018). Published by the American Society of Mechanical Engineers — distinct from ASQ (American Society for Quality). |
| ASTCT | American Society for Transplantation and Cellular Therapy. Issued 2019 CRS consensus grading (Lee et al.) adopted in CAR-T product labels. |
| ATMP | Advanced Therapy Medicinal Product (EU, Regulation EC 1394/2007): somatic cell therapy, gene therapy, tissue engineering. |
| BLA | Biologics Licence Application (USA, 21 CFR Part 601): regulatory pathway for CAR-T therapies. |
| Class A (PL) | Profile-likelihood class: CI fully closed within parameter bounds at ΔJ = 0.010. Parameter is practically identifiable from current benchmark data. |
| Class C (PL) | Profile-likelihood class: CI touches bounds at ΔJ = 0.020. Parameter not identifiable from current benchmark — fix at best-fit with documented justification. |
| COMBINE archive | Open Modelling EXchange format (.omex; Bergmann et al. 2014): standard container for SBML, SED-ML, data, and provenance metadata. |
| CoU | Context of Use (FDA MIDD 2019): specific role, scope, and limitations within which a model is applied. |
| CRS | Cytokine Release Syndrome: systemic inflammatory response to CAR-T. Graded by ASTCT 2019 on clinical symptoms (fever, hypotension, hypoxia) — not cytokine thresholds. |
| GSA | Global Sensitivity Analysis: Sobol' variance-based S₁ (first-order) and S_T (total-order) indices. Completed: N=1,024, 40,960 evaluations (Supplementary S3). |
| IND | Investigational New Drug Application (USA, 21 CFR Part 312): authorises first-in-human clinical trials. |
| MAA | Marketing Authorisation Application (EU, EMA): analogous to BLA. |
| MAS / HLH | Macrophage Activation Syndrome / Haemophagocytic Lymphohistiocytosis: life-threatening CAR-T complication; IFN-γ is primary diagnostic biomarker. |
| MIDD | Model-Informed Drug Development (FDA 2019): framework for using computational models in regulatory submissions. |
| PL | Profile-Likelihood identifiability: evaluates J(θᵢ) across a grid with other parameters fixed. Completed for 14 active parameters (Supplementary S4). |
| QSP | Quantitative Systems Pharmacology: mechanistic mathematical modelling of drug–biological system interactions. |
| S₁ | Sobol' first-order index: fraction of output variance explained by a single parameter alone (main effect). |
| S_T | Sobol' total-order index: full contribution including all parameter interactions. S_T − S₁ > 0.05 indicates substantial non-additivity. |
| SBML | Systems Biology Markup Language (Hucka et al. 2003): open XML standard; Level 3 Version 2 used here. |
| Tocilizumab | Humanised anti-IL-6 receptor (IL-6R) monoclonal antibody; FDA-approved for CRS Grade ≥ 2 management. Blocks IL-6 signalling via its receptor, not IL-6 itself; serum IL-6 rises post-administration. |
| Verification | 'Solving the equations right' (ASME V&V 40 §3.1): confirming computational implementation matches intended equations. |
| Validation | 'Solving the right equations' (ASME V&V 40 §3.1): confirming model adequately represents real-world system. |
